# An image is worth a thousand species: mapping plant biodiversity with citizen science, remote sensing, and deep learning

**DOI:** 10.1101/2022.08.16.504150

**Authors:** Lauren Gillespie, Megan Ruffley, Moisés Expósito-Alonso

## Abstract

Anthropogenic habitat destruction and climate change are altering the composition of plant communities worldwide^1,2^. However, traditional species distribution models cannot detect rapid, local plant species changes due to their low spatial and temporal resolution^3,4^, and remote sensing models can only identify changes in coarse vegetation categories^5,6^. Here we combine open-access remote sensing imagery, citizen science observations, and deep learning to create a multi-species prediction model at high spatial and temporal resolution. We train a novel deep convolutional neural network using ∼half a million observations within California to simultaneously predict the presence of over 2,000 plant species at meter-level resolution. This model—*deepbiosphere*—accurately performs many key biodiversity monitoring tasks, from fine-mapping geographic distributions of individual species and communities, to detecting rapid plant community changes in space and time. *Deepbiosphere* shifts the paradigm for species distribution modeling, providing a roadmap for inexpensive, automatic, and scalable detection of anthropogenic impacts on species worldwide.

---

Human-driven land use and climate change are impacting the ranges of plant species worldwide^1,2,7^ at a rapid pace^8^. This resulting geographic shift in plant biodiversity^9^ impacts critical ecosystem services such as carbon sequestration^10^, primary productivity^11^, and climate regulation^12^. However, the scale at which these losses occur is mismatched with that of classical methods for monitoring plant biodiversity. Plot-based monitoring networks lack coverage to capture biodiversity change at short timescales^13^, remote sensing-based change detection algorithms^5,6^ aggregate species into coarse vegetation categories, and species distribution models fail to capture rapid ecosystem transitions in space^14^ and time^4^ through their use of low-resolution decadal climatic variables^15^ (**Fig. S1**). New high-resolution methods are thus needed to quantify rapid spatiotemporal shifts of plant biodiversity in response to land and climate change^16^.

Here we introduce a deep learning-based method for high spatial and temporal resolution species-level plant biodiversity monitoring from publicly available remote sensing imagery, climate data, and citizen science observations. We show that this method improves the prediction of thousands of plant species ranges simultaneously, generates meter-level resolution maps of species ranges, detects spatial and temporal changes in biodiversity, and improves the mapping of both native vegetation and agricultural land use.

We focused on the species-diverse and heterogeneous region of California, where the flora has been extensively studied^17,18^, high-quality remote sensing imagery abounds^19^, and citizen science observations are dense enough to provide sufficient data for training. To compare our approach to classical species distribution models, starting with ∼1 million observations from the Global Biodiversity Information Facility^20^ we curated a dataset of over 650,000 research-grade primarily *iNaturalist* citizen science observations for 2,221 vascular plant species in California^21^ (see **Methods**). We then paired each observation with a 256 x 256 pixel, 1-m-resolution RGB-Infrared aerial image from the National Aerial Imagery Program (NAIP)^22^ (**Figs. 1A, S2, Table S1, Methods**; open-source code to reproduce dataset: github.com/moiexpositoalonsolab/deepbiosphere). In contrast to previous biodiversity datasets^23^, this dataset was multi-label; that is, each aerial image was linked to a list of species observations rather than a single species (**Supplementary Methods** [**SM**] **1.2**). We speculated that predicting multiple species simultaneously could significantly improve remote-sensing-based species distribution models^16^, as biotic interactions and habitat preferences may be captured through species co-occurrences, while rare species ranges may be more accurately predicted from their associations with more common species^24–26^.

**Fig. 1.**
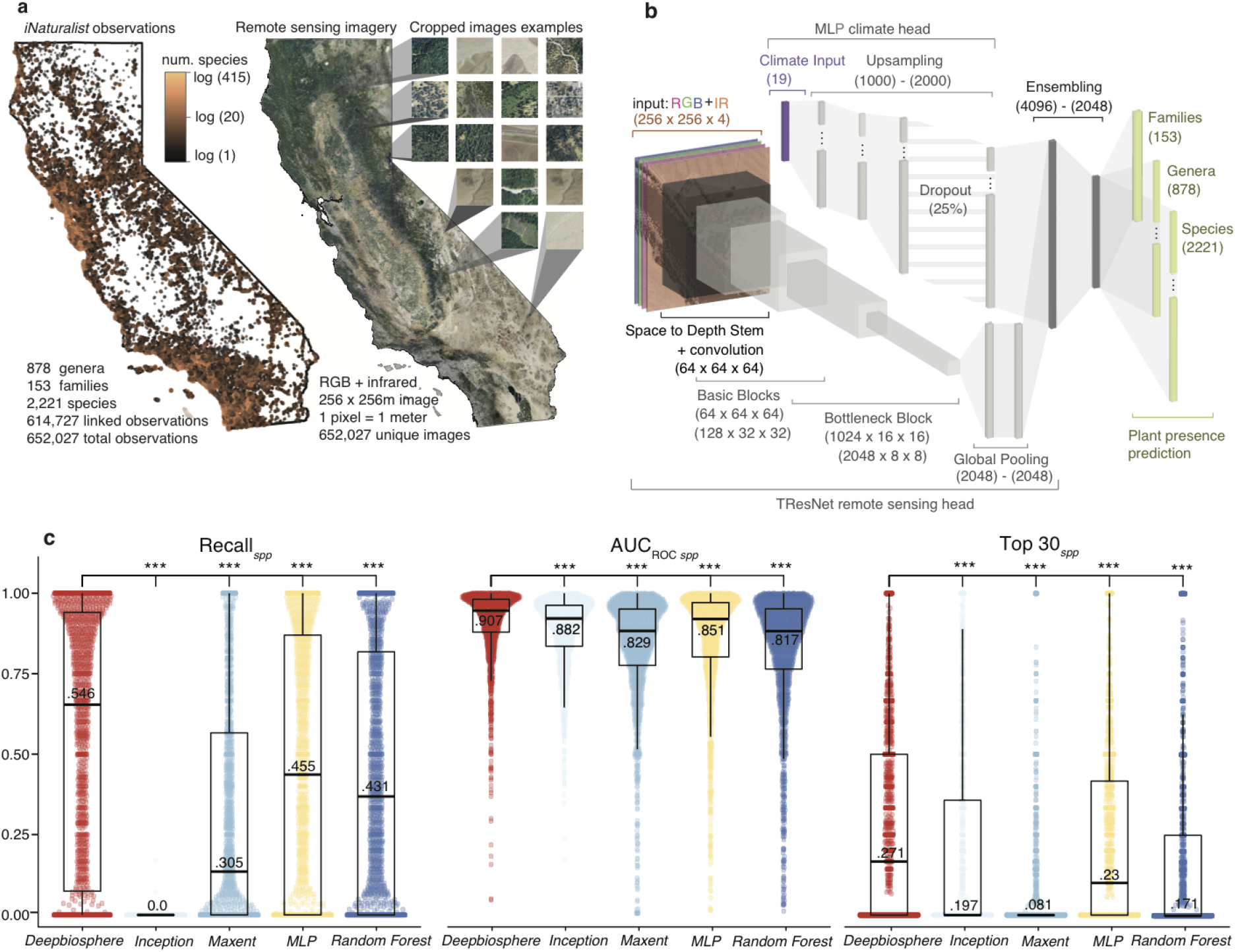
Training a deep neural network to predict plant biodiversity. **a**, Visualization of the dataset used for comparing plant species distribution models. Over 650,000 *iNaturalist* observations of 2,221 plant species were linked to 256 x 256m images cropped from 2012 National Aerial Imagery Program (NAIP)^19^ remote sensing imagery and climate data. **b**, The *deepbiosphere* architecture, which combines a convolutional neural network (CNN)^29^ trained using remote sensing imagery with a multi-layered perceptron (*MLP*) network head trained using climate variables to predict plant families, genera, and species. **c**, Comparison of *deepbiosphere’s* performance to classical climate-based species distribution models (SDMs) including *Maxent, Random Forest*, as well as *MLP*, and NAIP-based *Inception V3* CNN architecture^32^ using three common accuracy metrics. Metrics are reported per-species for the 1,541 species shared between the training and testing set with average score annotated on each boxplot (**Fig. S4A**). *spp* = per-species, AUC = area under the curve; ROC = receiver operating characteristic curve. Stars indicate results from unpaired student’s *t*-test, with *** indicating a *P*-value < 10^-3^.

To capture spatial variation in habitat and land use for accurate plant range mapping^27,28^, we employed a multilabel-optimized residual deep convolutional neural network (CNN) architecture called *TResNet*^*29*^. Training a modified *TResNet* model (**Table S2**) on our citizen science dataset presented significant challenges because these data often over-represent common species and densely populated regions (**Fig. S3**). To control for these sampling biases, we created a new variation of the popular binary cross entropy loss function that differentially downweighs the network learning from absent species based on the estimated per-location incompleteness of the species presence data (**SM 3.2.1**). Therefore, our new loss function emphasizes learning from the few species that are present in an image as much as from the species that are absent (as “absent” species may indeed be present though not recorded on *iNaturalist*). We compared this new loss function to other popular functions on a test split of the data (**Fig. S4A, SM 1.3.1**), and our tailored loss function performed best across a suite of accuracy metrics designed to test different model skills, including precision, recall, area under the receiver operating characteristic curve, and top 30 accuracy^30,31^ (**Table S6, SM 2**).

Since climate also predicts species ranges—albeit at larger spatial scales^28^—next we simultaneously processed both low-resolution climate data and high-resolution remote sensing data by designing a novel CNN architecture that combines a feed-forward multilayer perceptron (MLP) with the *TResNet* architecture^29^ (**Fig. 1B, Table S3**). This multi-headed architecture outperformed both classical species distribution models and a previous *Inception V3*-based CNN approach^32^ (**Table S4**) that used a standard computer vision architecture, loss function, and single-label objective (**Fig, 1C, S6-7, Tables 1, S7, Methods**). Our multi-headed architecture further outperformed CNNs trained only with remote sensing data as well as MLPs trained only with climate data (**Tables S5-7**). This architecture also excelled at extrapolation during a ten-fold spatial cross-validation assessment (**Fig. S4B, Tables 2, S8**). This top-performing CNN—named *deepbiosphere*—can predict the presence of thousands of plant species simultaneously from remote sensing imagery, making it a one-of-a-kind, high-resolution joint species distribution model.

**Table 1.** Comparing the accuracy of species distribution models across uniform test set examples. Median [IQR] of per-species accuracy metrics for each species distribution model and baseline random or frequency-based predictions. The testing examples withheld from the training set were at least 1.3 km away from any training point and were sampled from across all of California (**Fig. S4A, Methods**). For accuracy results per-image, see **Table S7**.

| Model name | Resolution | Loss | AUC<sub>ROC spp</sub> | AUC<sub>PRC spp</sub> | Recall<sub>spp</sub> | Prec<sub>spp</sub> | F1<sub>spp</sub> | Accuracy | mAP | Top 30<sub>spp</sub> |
| --- | --- | --- | --- | --- | --- | --- | --- | --- | --- | --- |
| *Deepbiosphere* | 256m | Scaled BCE | **0.946**<br>[0.88–0.98] | **0.036**<br>[0.01–0.11] | **0.654**<br>[0.07–0.94] | **0.018**<br>[0.0–0.05] | **0.035**<br>[0.00–0.09] | **0.776** | **0.161** | **0.167**<br>[0.0–0.5] |
| *Climate MLP* | ~1,000m | Scaled BCE | 0.920<br>[0.80–0.97] | 0.0342<br>[0.01–0.09] | 0.436<br>[0.0–0.87] | 0.013<br>[0.0–0.04] | 0.025<br>[0.0–0.08] | 0.684 | 0.115 | 0.1 [0.0–0.42] |
| *Inception*<sup>32</sup> | 256m | CE | 0.921<br>[0.84–0.96] | 0.020<br>[0.01–0.06] | 0.0<br>[0.0–0.0] | 0.0<br>[0.0–0.0] | 0.0 [0.0–0.0] | 0.005 | 0.153 | 0.0<br>[0.0–0.36] |
| *Maxent* | ~1,000m | N/A | 0.883<br>[0.78–0.95] | 0.018<br>[0.00–0.07] | 0.135<br>[0.0–0.57] | 0.005<br>[0.0–0.03] | 0.009<br>[0.0–0.06] | 0.276 | 0.026 | 0.0<br>[0.0–0.0] |
| *Random Forest* | ~1,000m | N/A | 0.882<br>[0.76–0.95] | 0.024<br>[0.00–0.09] | 0.368<br>[0.0–0.82] | 0.009<br>[0.0–0.04] | 0.017<br>[0.0–0.07] | 0.394 | 0.057 | 0.0<br>[0.0–0.25] |
| *Random* | NA | N/A | 0.501<br>[0.49–0.51] | 0.002<br>[0.00–0.01] | 0.5<br>[0.48–0.52] | 0.002<br>[0.00–0.00] | 0.003<br>[0.00–0.01] | 0.502 | 0.007 | 0.0<br>[0.0–0.02] |
| *Frequency* | NA | N/A | 0.5<br>[0.5–0.5] | 0.002<br>[0.0–0.01] | 0.0<br>[0.0–0.0] | 0.0<br>[0.0–0.0] | 0.0<br>[0.0–0.0] | 0.066 | 0.038 | 0.0<br>[0.0–0.0] |

*MLP* = multilayer perceptron; BCE = binary cross-entropy; CE = cross-entropy; *spp* = per-species; AUC<sub>ROC</sub> = area under receiver operating curve; AUC<sub>PRC</sub> = area under precision-recall curve; Prec = precision; mAP = mean average precision.

**Table 2.** Comparing the accuracy of selected species distribution models on latitudinal cross-validation blocks. Median [IQR] of the accuracy metrics across ten latitudinal cross-validation blocks (**Fig. S4B, Methods**). For accuracy results per-image, see **Table S8**.

| Model name | Resolution | AUC<sub>ROC spp</sub> | AUC<sub>PRC spp</sub> | Recall<sub>spp</sub> | Prec<sub>spp</sub> | F1<sub>spp</sub> | Accuracy | mAP | Top 30<sub>spp</sub> |
| --- | --- | --- | --- | --- | --- | --- | --- | --- | --- |
| *Deepbiosphere* | 256m | **0.820**<br>[0.79–0.83] | **0.072**<br>[0.07–0.08] | **0.340**<br>[0.32–0.36] | **0.052**<br>[0.05–0.06] | **0.079**<br>[0.08–0.08] | **0.684**<br>[0.64–0.72] | **0.189**<br>[0.17–0.21] | **0.140**<br>[0.13–0.16] |
| *Climate MLP* | ~1,000m | 0.687<br>[0.68–0.73] | 0.054<br>[0.05–0.06] | 0.218<br>[0.19–0.24] | 0.039<br>[0.03–0.04] | 0.058<br>[0.05–0.06] | 0.493<br>[0.473–0.53] | 0.114<br>[0.10–0.13] | 0.090<br>[0.08–0.11] |
| *MaxEnt* | ~1,000m | 0.714<br>[0.71–0.73] | 0.053<br>[0.05–0.06] | 0.326<br>[0.26–0.48] | 0.036<br>[0.03–0.038] | 0.054<br>[0.04–0.06] | 0.426<br>[0.36–0.64] | 0.039<br>[0.03–0.05] | 0.050<br>[0.04–0.05] |
| *Random Forest* | ~1,000m | 0.702<br>[0.69–0.71] | 0.053<br>[0.05–0.06] | 0.445<br>[0.34–0.58] | 0.036<br>[0.03–0.04] | 0.057<br>[0.05–0.06] | 0.510<br>[0.437–0.69] | 0.054<br>[0.05–0.06] | 0.086<br>[0.08–0.09] |
| *Frequency* | N/A | 0.5<br>[0.5–0.5] | 0.053<br>[0.05–0.06] | 0.014<br>[0.01–0.02] | 0.002<br>[0.001–0.002] | 0.003<br>[0.002–0.003] | 0.103<br>[0.07–0.12] | 0.095<br>[0.057–0.10] | 0.021<br>[0.02–0.023] |

*MLP* = multilayer perceptron; *spp* = per-species; AUC<sub>ROC</sub> = area under receiver operating curve; AUC<sub>PRC</sub> = area under precision-recall curve; Prec = precision; mAP = mean average precision.

Unlike previous methods, *deepbiosphere* can generate meter-resolution biodiversity predictions (**Fig. S8, SM 4.1.1**), which should allow for the detection of keystone species even in highly fragmented and heterogeneous landscapes. To test this, we focused on the iconic redwood forests of California (*Sequoia sempervirens*, 2,349 observations in dataset). These forests were heavily logged in the mid-20th century, leaving only 5% old-growth forest scattered across remote pockets of California^33^. Nearly half of this remaining mature forest is contained in Redwood National and State Parks and can be visually identified from aerial imagery as a dark strip running from north to south along Redwood Creek (**Fig. 2A**). Given a few examples (**Fig. S9, Table S9)**, human annotators can detect this old-growth redwood forest from aerial images with high accuracy but at a slow pace, averaging 5 seconds per-image (**Figs. 2B, S10, Methods, SM 4.1.2**). Surprisingly, the *deepbiosphere*-generated map of probability of redwood presence detected a much broader distribution of redwood than the human-annotated maps, capturing the same old growth remnants as human annotators plus areas beyond (**Fig. 2C**). Comparing these results to the official National Park Service (NPS) alliance-level vegetation map for the area^34^ (**Fig. 2D**) revealed that most of the region is in fact covered by redwood forest, with *deepbiosphere* detecting both the young regrowth after clear-cutting (blue regions) in addition to the old-growth remnants easily detected by human annotators (red regions). Further comparison against classical species distribution models and another recent CNN-based approach—*Inception V3* (ref. ^32^)*—* revealed that *deepbiosphere* better predicts the full extent of known redwood forest distribution (**Fig. 2F, S11**) and test observations (**Table S10**). Thus, *deepbiosphere* more accurately predicts the true extent of redwood forest compared to previous approaches and is nearly 100X more efficient than manual human annotations.

**Fig. 2.**
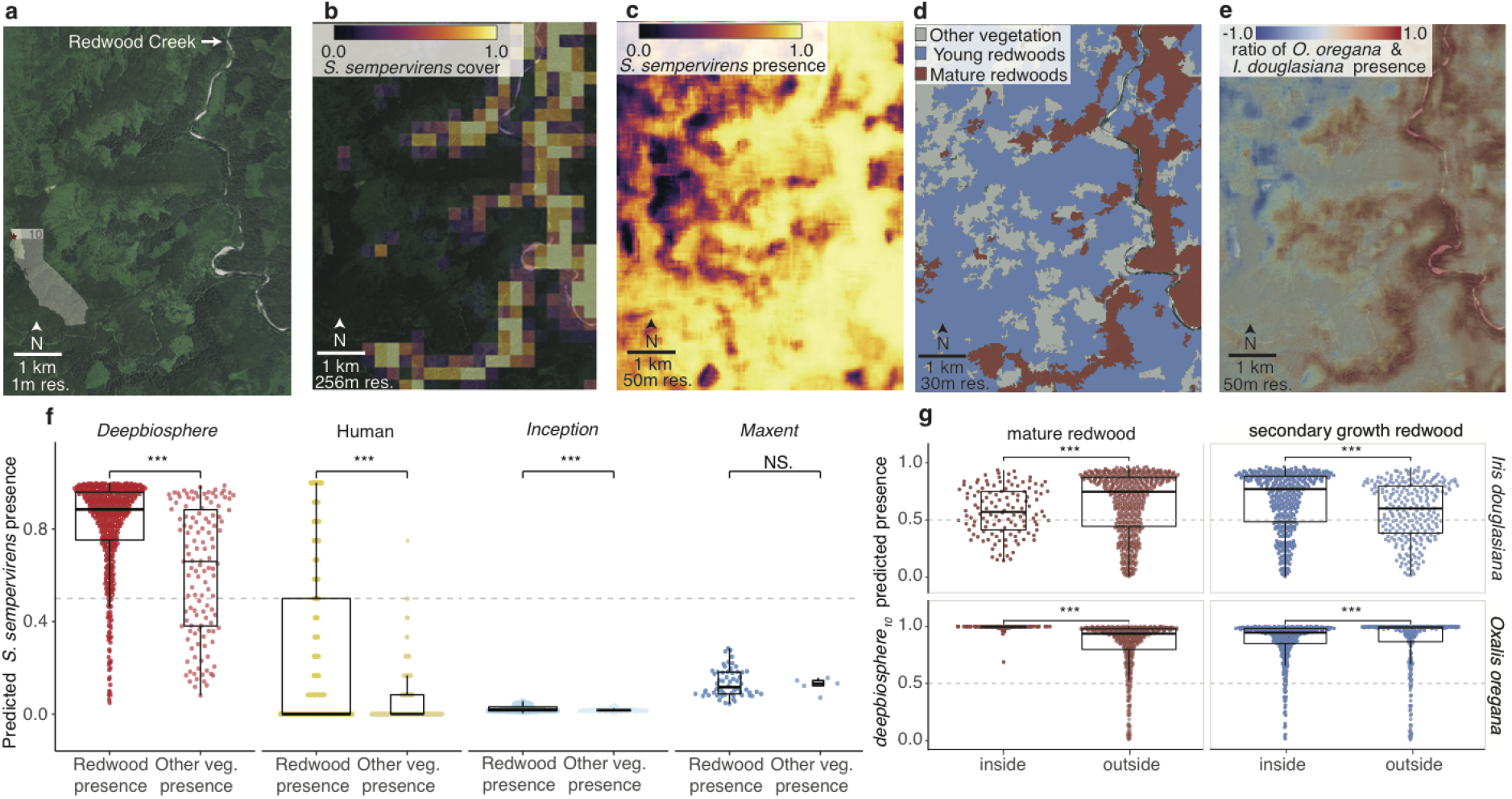
*Deepbiosphere* predictions of field-validated coastal redwood forest species. **a**, NAIP aerial imagery of Tall Trees redwood grove (*Sequoia sempervirens*) in California’s Redwoods National and State Parks. The region contains some of the last old-growth redwood remnants in the world, visible as the dark green line along Redwood Creek. **b**, Human annotation of redwood forest cover, based on examples of other old-growth redwood groves (**Fig. S9**). Human annotators can distinguish mature groves (**Fig. S10**). **c**, *Deepbiosphere* predicted presence of *S. sempervirens*. **d**, Official National Park Service (NPS) vegetation map^34^ highlighting mature redwood (dark red) and young regrowth (blue) vegetation classes. **e**, *Deepbiosphere* difference in predicted presence of two understory species: *Oxalis oregana*, which has a preference for mature redwood stands, and *Iris douglasiana*, which has a preference for secondary-growth redwood forest^34^. **f**, Comparison of the predicted presence of *S. sempervirens* at every pixel for *deepbiosphere* and other models based on whether the associated vegetation class for said pixel from **d** is redwoods-dominant. *Deepbiosphere* and *Maxent* were fitted without any examples from the region, while *Inception* did see examples from the study area during training. **g**, *Deepbiosphere* predicted presence of two understory species in **e** at every pixel, annotated by NPS vegetation class type^34^ from **d** (**Methods**). Stars indicate an unpaired student’s *t*-test with *** indicating a *P*-value < 10^-3^ and N.S. indicating a *P*-value of ≥ 10^-1^ .

While many large plant species have been mapped using remote sensing^35,36^, we were curious whether *deepbiosphere* could also predict understory species that are not directly visible from co-existence patterns. We focused on two charismatic understory herbs found in Redwood National and State Parks: redwood sorrel (*Oxalis oregana*, 1,063 observations in dataset) and Douglas iris (*Iris douglasiana*, 763 observations in dataset). *Deepbiosphere* predicted redwood sorrel as more likely to be present mainly in mature redwood groves, matching field-validated associations with the cool and moist understories of these old-growth trees^34^ (**Fig. 2E, 2G, S12B, SM 4.1.3**). Meanwhile, Douglas iris was predicted as more likely to be present mainly in young redwood regrowth, confirming its known preference for the semi-shade of young redwood forests^34^ (**Fig. 2E, 2G, S12D**). These associations are further supported by analyses of other well-known understory species associated with either mature or regrowth redwood forests^34^ (**Figs. S12-13**). *Deepbiosphere’s* ability to correctly map the distribution of both canopy trees and small herbaceous plants extends to other habitats, including Southern California’s mediterranean ecosystems where previously mapped vegetation distributions^37,38^ match *deepbiosphere’s* predictions (see case studies of *Quercus lobata, Q. berberidifolia, Ceanothus cuneatus, Bromus diandrus, Arctostaphylos glandulosa, Adenostoma fasciculatum*, **Fig. S14-16**). Together, these results demonstrate that remote-sensing imagery linked to citizen science observations contain useful information on the key drivers of individual species distributions—such as local topography, land use, and co-occurring species^24,28^—and allow *deepbiosphere* to predict species ranges at fine spatial scales.

Given *deepbiosphere’s* ability to predict local species presence, we asked if it could also detect spatial and temporal trends in whole-community turnover at high resolution. To study spatial changes in biodiversity, we focused on north Marin County (**Fig. 3A**) as this region sits at the boundary between the Marine Coastal Forest and Mediterranean California ecoregions, with a heterogeneous mix of native vegetation, urban areas, and agricultural land (**Fig. S17C, S18**). We specifically tested whether an image-processing-based edge detection algorithm could delineate community-level spatial changes (e.g., ecotones) using *deepbiosphere’s* species-level predictions. For each 256 x 256 m image in the region, we generated a vector of 2,221 species probabilities from *deepbiosphere* and calculated the average Euclidean distance between this vector and that of each of the eight neighboring images. We refer to this unitless estimation of species turnover as *spatial community change* (**Fig. S19, Methods, SM 5.1**). *Deepbiosphere’s* spatial community change map for north Marin county captured both mountain-to-valley and developed-to-undeveloped edges (**Fig. 3B**), transitions that are usually not well-captured by kilometer-resolution climate or elevation maps (**Fig. S1**). Encouragingly, *deepbiosphere’s* spatial community change predictions correlate strongly with the official Marin County fine-scale vegetation map^39^ (Pearson’s *r* = 0.45, *P*-value < 2.2 × 10^-16^, **Figs. 3B, 3C**), specifically with the number of unique vegetation classes present in each 256 x 256 m image (**Fig. S17C, S18**). This prediction accuracy far exceeds edge detection results based on raw remote sensing data (Pearson’s *r* = 0.24, *P*-value < 2.2 × 10^-16^, **Fig. 3D, Fig. S17D**) and is not simply a holdover effect of observation density, as spatial community change does not correlate with the density of *iNaturalist* observations in the study area (**Fig. S17E-F**). These findings suggest that, when aggregated, *deepbiosphere’s* species-level predictions capture important spatial patterns in biodiversity, such as species turnover across neighboring communities.

**Fig. 3.**
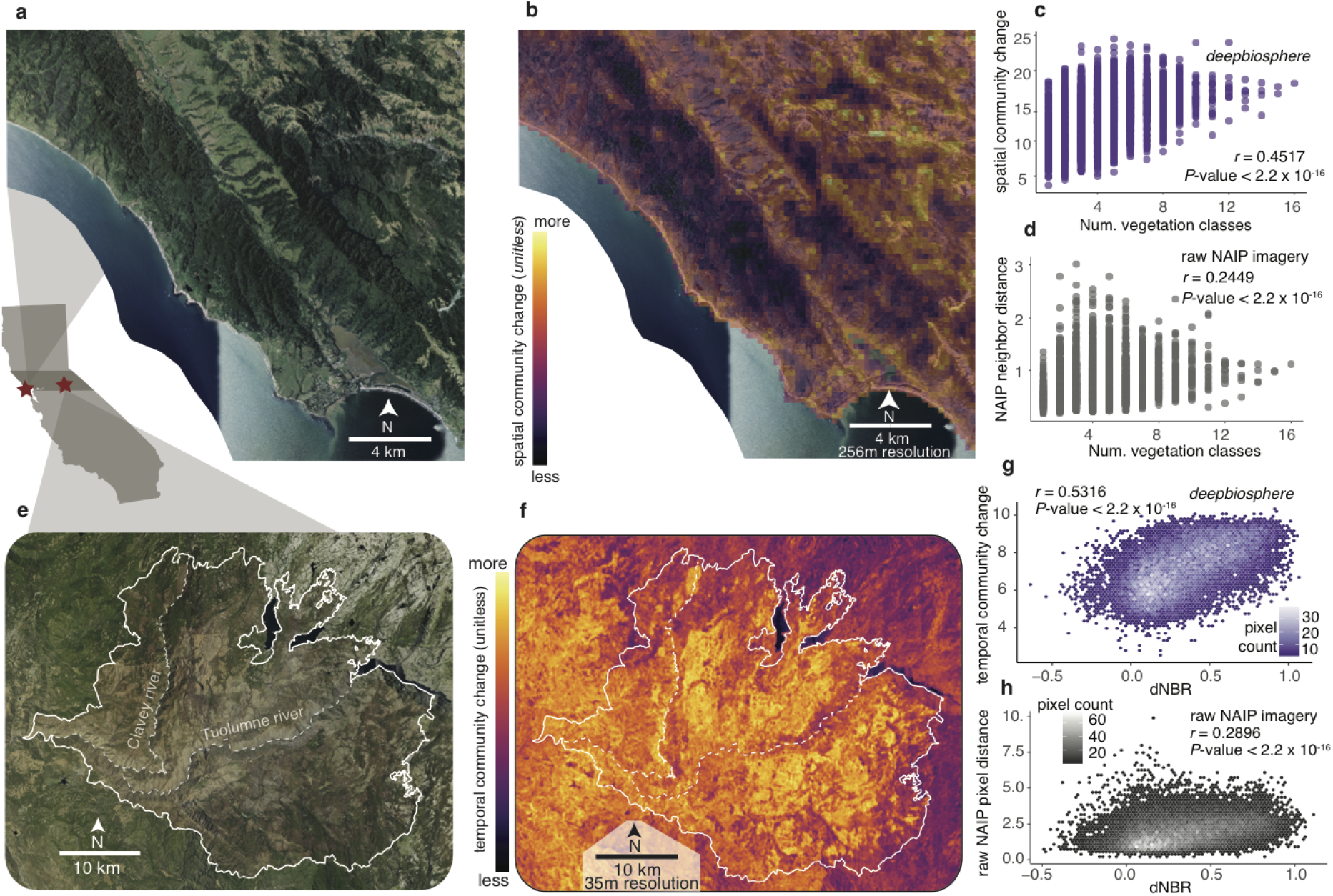
Detection of high-resolution spatial and temporal community changes using *deepbiosphere*. **a**, NAIP aerial imagery of northern Marin county at the boundary of two major ecoregions. **b**, *Deepbiosphere* spatial community change calculated using an edge-detection algorithm applied to the predicted presence of all 2,221 plant species (**Fig. S19, Methods, SM 5.1**). **c**, Comparison of the number of unique fine-scale vegetation types^39^ present in each 256 x 256 m plot and *deepbiosphere*’s spatial community change predictions. **d**, Comparison of the number of unique fine-scale vegetation types^39^ present at each 256 x 256 m plot and an edge-detection algorithm run on raw NAIP RGB-I images. **e**, 2014 NAIP aerial imagery of Sierra foothills after severe Rim Fire of 2013 (fire boundary in white). **f**, *Deepbiosphere* temporal community change calculated using the Euclidean distance between predicted species presence in 2012 and 2014 (**Fig S21, Methods, SM 5.2**). **g**, Comparison of empirical burn severity metric—difference in normalized burn ratio (dNBR, **Fig. S20C**)^40^—with *deepbiosphere’s* temporal community change from **f. g**, Comparison of dNBR with Euclidean distance between raw NAIP RGBI-I imagery from 2012 and 2014 (**Fig. S20E**).

To study *deepbiosphere*’s capacity to detect rapid temporal changes, we focused on the 2013 Rim wildfire in the western California Sierra foothills (**Fig. 3E**). Remote sensing data, which is often collected at week-to-yearly timescale, should contain signals to detect this rapid temporal change thanks to their fine temporal resolution^16,22^. The Rim fire’s burn scar is clearly visible on NAIP imagery taken after the fire as compared to before (2014 vs 2012, **Fig. S20A-B**). Again using the Euclidean distance metric, we compared *deepbiosphere’s* vectors of 2,221 species probabilities for NAIP imagery taken in 2012 with predictions made using 2014 imagery from the same 256 x 256 m areas, a metric we refer to as *temporal community change* (**Fig. S21, Methods, SM 5.2**). Our metric visually indicated that the most severe burning occurred east of the Clavey river and south of the Tuolumne river (**Fig. 3F**). On average, pixels inside the fire’s boundary had higher temporal community changes than those outside (**Fig. S20F**, unpaired student’s *t*-test, *P*-value < 2.2 × 10^-16^). These findings are corroborated by the difference in Normalized Burn Ratio (dNBR)^40^—an NDVI-like metric routinely used in forestry to map burn severity (**Fig. 20C**)—which significantly correlated with our metric (Pearson’s *r* = 0.53, *P*-value < 2.2 × 10^-16^, **Fig. 3G**).This far exceeds results calculated based on raw remote sensing data, specifically the pixel-wise difference between 2012 and 2014 NAIP imagery, which naturally should contain a signal in the green and infrared band (Pearson’s *r* = 0.29, *P*-value < 2.2 × 10^-16^, **Figs. 20E, 3H**). Finally, *deepbiosphere’s* species-based approach also enabled the detection of individual species change in this disturbed landscape (see case studies of *Populus tremuloides, Eriophyllum confertiflorum* **Figs. S22, S23**). Overall, these findings show *deepbiosphere* can capture rapid ecological transitions at species-level resolution, substantially improving the study of plant species and community dynamics in rapidly changing environments.

Finally, we wondered whether we could use *deepbiosphere’s* ability to finely detect plant communities for important practical applications using a limited number of examples, such as delineating natural vegetation types or mapping croplands. Current machine learning-based strategies for these approaches require large datasets of hundreds of thousands of labeled data points to train models—even when focusing on only a few vegetation or land use classes^41,42^. To use *deepbiosphere* for downstream tasks with only a few thousand examples, we employed the machine-learning concept of feature extraction (**SM 6.1**) in two final case studies. First, we focused on fine-scale vegetation mapping in Redwood National and State Parks where 1,009 field-validated vegetation plots are publicly available and such a map already exists^34^ (vegetation classes were provided at the alliance level [multi-species consortia] from manual relevé botanical surveys, **Figs. 4A, S24B**). To generate our own fine-scale vegetation classifier from these plot data, we extracted feature vectors from the hidden layer parameters of *deepbiosphere* at each plot. Using these feature vectors, we fitted a random forest classifier to distinguish the 30 vegetation classes used by the National Park Service (**SM 6.2, Methods**). We found that using *deepbiosphere*’s feature vectors improved the prediction of vegetation type at test plots across 10 cross-validation trials by an average of 8% compared to the official National Park Service vegetation map^34^ (56% vs. 48% average accuracy, unpaired student’s *t*-test, *P*-value = 4.4 × 10^-4^, **Table S11, Fig. 4C, Methods**) and across many vegetation classes (**Table S12, Figs. S26, S27**). Finally, the 50-m-resolution vegetation map generated from our feature vector classifier closely matched the official National Park Service map^34^ (**Fig. 4B**, 60% top-1, 90% top-5 pixel agreement; most common class 29% of pixels, top-5 most common classes 71% of pixels).

**Fig. 4.**
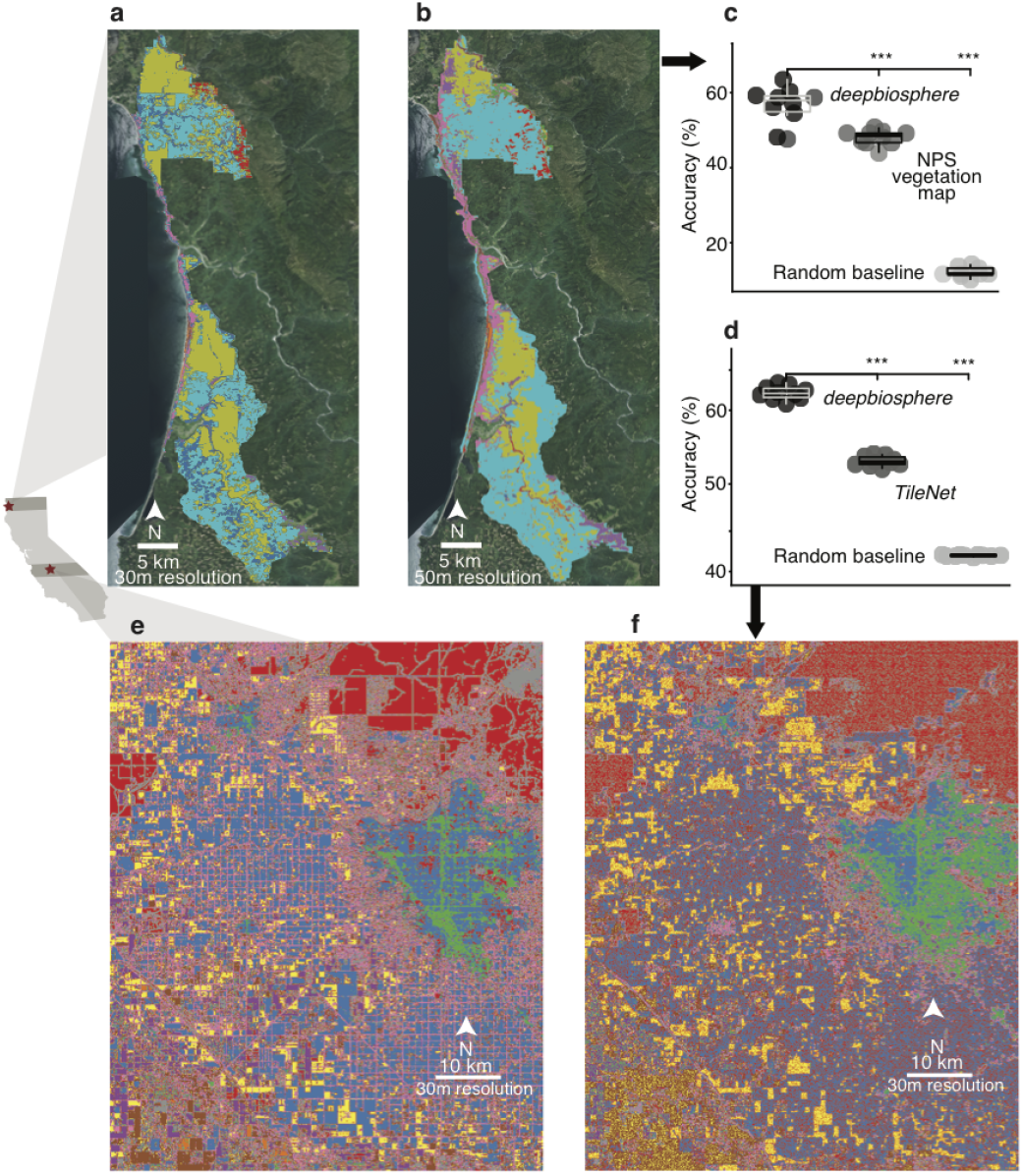
Application of *deepbiosphere* to vegetation and crop mapping. **a**, Official National Park Service (NPS) alliance-level vegetation map of Redwood National and State Park comprising 30 vegetation types (see color legend in **Fig. S25**). **b**, Alliance-level vegetation map created using a random forest classifier fitted from *deepbiosphere*’s learned features, based on a dataset of 1,009 validated field plots (**Fig S24B**). **c**, Ten-fold accuracy of *deepbiosphere-*based random forest vegetation classifier compared to the official NPS vegetation map and random Gaussian noise baseline using 189 previously unseen plots bootstrapped from 489 NPS accuracy assessment field plots (**Methods**). **d**, Accuracy of ten trials of *deepbiosphere*-based random forest cropland classifier compared to a recent unsupervised CNN approach *TileNet* (from ref. ^44^) and a random Gaussian noise baseline at 1,000 previously unseen test locations (**Fig. S28D**). **e**, 2016 USDA cropland data layer for the San Joaquin Valley comprising 69 unique crop types^43^ (see color legend in **Fig. S29**). **f**, Cropland map created using a random forest classifier fitted with *deepbiosphere*’s learned features using 1,000 randomly selected locations from **a** (**Fig. S28D**). Asterisks represent unpaired Student’s *t*-test results with *P*-values as follows: *** < 10^-3^; ** < 10^-2^_;_ * < 10^-1^; N.S. ≥ 10^-1^ .

As a second application of using *deepbiosphere* for downstream mapping with little data, we tried predicting cropland type from *deepbiosphere’s* feature vectors using only 1,000 examples. We focused on the San Joaquin Valley in Central California, home to a diversity of crop types mapped each year by the United State Department of Agriculture^43^ (USDA) (**Fig. S28C**). As before, we fitted a random forest classifier to predict 69 crop types from the 2016 crop map using *deepbiosphere’s* feature vectors at 1,000 randomly sampled training locations (**Fig. S28D, Methods**). Compared to *TileNet*—an unsupervised, deep-learning-based cropland classifier which is state-of-the-art^44^—our feature vector classifier exhibited a nearly 10% higher performance at test locations across 10 trials (62% vs. 54% average accuracy, unpaired student’s *t*-test, *P*-value = 5.21 × 10^-15^, **Table S13, Fig. 4D**) and typically exhibited higher accuracy per-crop class (**Table S14, Fig. S30, S31**). The resulting 30-m-resolution crop type map of San Joaquin Valley generated from our feature vector classifier (**Fig. 4F**) agreed strongly with the official, field-validated USDA cropland data layer^43^ (**Fig. 4E, Fig. S30**, 57.17% top-1, 85.47% top-5 pixel agreement; most common class 42% of pixels, top-5 most common classes 91% of pixels). This is particularly encouraging because domesticated crop species were not present in our biodiversity dataset. These results imply the complex ecological patterns *deepbiosphere* learned from the visual features of remote sensing images can be extrapolated to non-natural habitats, including those transformed and managed by humans. *Deepbiosphere* may therefore also be a powerful tool for urban, crop, and wildlife planning at high spatiotemporal scales.

Motivated by a much-needed paradigm shift in species distribution modeling for biodiversity conservation^16,22^, *deepbiosphere* is a first-of-its-kind deep-learning algorithm that can simultaneously predict the presence of thousands of plant species from remote sensing data at unparalleled resolution. These predictions can be applied in a number of ways, from detecting spatiotemporal biodiversity change to mapping croplands and natural habitats. Since our freely-available, open-source network relies on public remote sensing imagery and simple to collect citizen science observations of plant species, *deepbiosphere* represents a significant step towards public, high-resolution plant biodiversity monitoring tools that can scale globally. Through continual training on ubiquitous satellite imagery and biodiversity data collected on a nearly-daily timescale, we expect that neural networks similar to *deepbiosphere* will soon generate predictions of biodiversity change in real-time worldwide. The features learned by these models will further enable high-quality downstream mapping tasks beyond vegetation and cropland identification from limited examples. Such publicly available algorithms and predictions will be critical to inform nature conservation at local and global scales.

## METHODS SUMMARY

### Species observations

We collected observations from kingdom *Plantae* using *GBIF*.*org* from the years 2015-2022. Only records observed by humans with a coordinate uncertainty radius of less than or equal to 120 m with no flagged geospatial issues were taken from within the state of California. We downloaded a total of 912,380 observations of 5,193 unique plant species^45^ and further filtered the dataset to only include vascular plants, remove duplicate observations of the same species within a 150m radius, remove species that contain all observations located within a 256m radius, remove observations that were not geographically located within the Global Administrative Area boundary of California^46^, and remove observations that were not located within both the climatic and remote sensing imagery rasters. To increase the density of observations and allow for multiple species within a single image, we used neighbor imputation to add any other species observed within an overlapping 256m radius to a given observation (**Fig. S2**, see **Supplemental Methods** [**SM] 1.2** for details). We finally removed any species that had fewer than 500 total observations in the dataset after neighbor imputation, leaving us with a total of 652,027 observations of 2,221 unique plant species^21^ (**Fig. 1A, Table S1**).

### Remote sensing data

To link species observations with images, we utilized aerial imagery from the National Agricultural Imagery Program (NAIP)^19^ which we downloaded for the entire state of California from 2012 and 2014 using Microsoft Azure’s NAIP data blob disk image on its West Europe and Eastern U.S. servers. For training the CNN models, we specifically used the NAIP data from 2012 at 1-m resolution to generate 256 x 256 pixel images, where 1 pixel corresponds to a 1 x 1 m resolution. We used all available bands for training, specifically the RGB and infrared color bands (**Table S1**). The 256 x 256 pixel images were extracted so that the geographic coordinates of the corresponding species observation mapped to the center of the image (**Fig. S2**).

### Climate variables

We used the 19 bioclimatic variables available from WorldClim Version 2 at 30 arc-second (approximately 1km) per-pixel resolution^15^. Variables were downloaded directly from the WorldClim Version 2 repository (http://www.worldclim.com/version2). Before fitting any model, all bioclimatic variables were normalized per-variable to mean 0 and standard deviation of 1 using the entire raster clipped to the outline of California.

### Train/test split and cross-validation

In order to properly validate and compare models, we split the dataset into multiple partitions. The first partition, which was used for hyperparameter tuning and loss comparison, was generated by randomly selecting observations uniformly from across the state (**SM 1.3.1, Fig. S4A**) which we refer to as the *uniform partition* and use the notation *modelname*_*unif*_ to refer to models trained using this partition of the dataset. For this train/test partition, we chose points uniformly across the state to maximize the number of unique ecosystems models would be evaluated on. To ensure the independence of training and testing set data due to spatial autocorrelation, we added all overlapping observations to the test set to guarantee that none of the remote sensing images and observations in the test set were present in the training set. To further ensure that there was no data leakage between the test and train set, only observations which were more than 1,300m away from any other non-overlapping observation were included. We chose an exclusion radius of 1,300m because the climate variable raster pixels converted from arc-seconds to meters can have a diameter of up to 1,200m, so any test set observation within that distance to any observation in the train set would have an identical input value as some observations used during fitting. Ultimately 12,277 observations (1.88% of the dataset) were set aside for testing in this split.

In order to provide cross-validation of the uniform train-test split and to test the extrapolation ability of all models, we also conducted a latitudinal ten-fold spatial holdout block validation by partitioning California into ten one-degree latitudinal bands (**SM 1.3.2, Fig. S4B**) which we refer to as the *spatial partition*, using the notation *modelname*_*k*_ to refer to models trained using points from the k-th spatial block (**Fig. S4B**). Training observations within 1,300m of the test band were removed to prevent data leakage as discussed above. For SDMs fitted with pseudo-absence points, all pseudo-absence points within the test bands were removed to ensure a fair comparison to presence-only models. Ultimately, the percentage of test points per-spatial block ranged from 1.40-25.35% of the entire dataset.

### Deep convolutional neural networks (CNNs) for species distribution models (SDMs)

We chose to use the medium-sized TResNet architecture, a CNN-based residual neural network^47^ which is GPU-optimized for fast inference speeds and is a state-of-the-art architecture for multi-label image classification in the computer vision community^29^. We modified the TResNet architecture to have four input channels in order to support the RGB and infrared NAIP imagery and added three fully connected output layers corresponding to predict three taxonomic ranks (family, genus, and species) to confer some phylogenetic signal during training. All TresNet-based CNNs are trained to predict each of the 2,221 plant species simultaneously. Along with this standard version of the TResNet architecture trained using only the NAIP aerial imagery (TResNet, **Table S2**) we also created our own custom CNN model which combines a TResNet head trained using NAIP imagery (**Table S2)** with a multilayer perceptron head trained using climate inputs (**Table S5)** which we refer to as the Joint TResNet / *deepbiosphere* model (**Table S3, Figure 1B**). Weights were initialized following best practices laid out in the original TResNet paper, using Kaiming He-style for CNN layers and zeroed out BatchNorm and residual connections^47^. For all analyses, TresNet-based CNN outputs were converted to independent probabilities using the sigmoid transformation.

We compared performance of the TResNet architecture trained on a variety of standard loss functions (**SM 3.2.1)**. The loss function compares how well CNN outputs align with a training set of observations and thus determines how well the model fits the data and learns from it. While we report results from a variety of common loss functions for fair comparison to previous work^32^, the final results use a new loss function we called scaled binary cross-entropy (scaled BCE) that overcomes limitations of common functions like cross-entropy (CE) which is best suited for single-label images, binary cross-entropy (BCE) which is best suited for multi-label images where absence of labels are informative, or the recent asymmetric focal loss (ASL) which is best suited for multi-label images where many mis-labels may occur. During training, our custom scaled BCE loss weighs the contribution of the few species present in any given image as much as the contribution of the many species that are absent, and provides the most accurate results (see **Table S6, SM 3.2.1**).

For comparison to previous work using CNNs to rank species presence from remote sensing imagery^32^, we trained an *Inception V3* architecture (*Inception*, **Table S4**) with softmax cross entropy loss using the official architecture implementation and initial weights from pytorch and using both the standard and auxiliary loss during training^48^. We utilize the standard dropout rate of 0.5 and a standard learning rate of 0.01, different but comparable hyperparameters to those used in previous work^32^. For all analyses, the *Inception* outputs were converted to a probability density function using the softmax transformation. While the *Inception* model is trained jointly across all species like *deepbiosphere*, the cross-entropy loss forces the *Inception* CNN to fit a probability density function across species, meaning that it has been trained to predict just one species at a time and making it impossible to use the model as a joint SDM effectively (see **Fig. S11B, SM 3.2.5**).

All CNNs were trained with standard mini-batch stochastic gradient descent using the Adam optimizer^49^. Batch size, learning rate, total number of epochs, memory usage, GPU, and training time are reported for each CNN in **Tables S2-S8**. Learning rates were chosen using a preliminary search across learning rates ranging from 10^-5^ to 10^-1^ and batch sizes were chosen depending on model size relative to the GPU size used for training.

To determine which training epoch to evaluate the deep neural networks, we calculated the loss function value and a subset of accuracy metrics at each epoch and selected the average optimal epoch across these metrics for evaluation (see **Fig. S5, SM 3.2.6**). For *deepbiosphere*_*unif*_, the average optimal epoch was four; for *InceptionV3*_*unif*_ ten; for the *MLP*_*unif*_ forty five. For comparing accuracy across loss functions, we evaluate all CNN SDMs at epoch 19 to standardize comparison across accuracy metrics (**Table S6**). For all further downstream evaluations, the average optimal epoch on the uniform partition test set was used (**Fig. S5**).

### Species Distribution Models based on climate rasters and other baselines

We use the popular *dismo* R package for species distribution modeling^50^ and compared against two popular SDM approaches: *Maxent* and downsampled single stacked random forest (RF). We chose these two models as they consistently had the best performance across dozens of models and hundreds of species in a large benchmarking experiment^51^. Following best practices from ref. ^51^, we removed all but one bioclim variable with a Pearson correlation coefficient higher than 0.8, leaving ten variables in total for modeling including Mean Diurnal Range, Max Temperature of Warmest Month, Minimum Temperature of Coldest Month, Annual Precipitation, Precipitation of Wettest Month, Precipitation of Driest Month, Precipitation Seasonality, Precipitation of Wettest Quarter, Precipitation of Warmest Quarter, and Precipitation of Coldest Quarter. For each species, we generated 50,000 background samples using a circular overlay across all points in the training dataset where the radius of each circle is the median distance between said species’ observations. We used the same number of presence and background points for both the random forest and *Maxent* models and we used the “nothreshold” option for *Maxent* and 1,000 trees with equal bootstrapping of positive and negative samples with replacement, with all other options set using *dismo* default. For a few species, the fitting process failed for *Maxent* and/or RF. For these species, in downstream accuracy analyses we impute an accuracy of 0 for all metrics.

For completeness, we also trained a fully-connected, feed-forward multilayer perceptron (MLP) on all 19 bioclim variables to predict all 2,221 species simultaneously as a climate-only deep learning baseline. The architecture consists of two fully-connected layers with 1,000 neurons each, followed by a dropout layer with a 0.25 dropout rate, then by two layers with 2,000 neurons each, before predicting species, genus, and family (**Table S5**). This MLP was also trained with standard mini-batch stochastic gradient descent using the Adam optimizer^52^ with a batch size of 1,000 and a learning rate of 0.0001. Finally, we also compared performance against two trivial baselines. The random baseline was calculated by drawing random values from a standard normal distribution ten times and averaging the accuracy metrics across these ten trials. The frequency baseline involved calculating the frequency of observations per-species on the training set, rescaling the probabilities to 0.001-1.0 and imputing these frequencies as the predicted probabilities at each test set example.

### Accuracy metrics

We utilized a wide variety of accuracy metrics from across a variety of relevant disciplines, from computer vision to species distribution modeling. For the full list of reported accuracy metrics and their explicit mathematical definitions, see **SM 2**. The reported accuracy metrics can be classified into three broad categories. The first is binary classification metrics, which capture an SDM’s ability to correctly predict the presence or absence of a species given a probability of presence threshold (we use a standard 0.5 threshold). We report precision, recall, and F1 score both per-species and per-image, along with single-label accuracy. The second category of metrics—discrimination metrics—calculate an SDM’s performance across a wide range of presence thresholds and describe the relationship between threshold change and performance change. Discrimination metrics are commonly reported in species distribution modeling work and reported metrics include area under the receiver operating characteristic curve (AUC_ROC_) and area under the precision-recall curve (AUC_PRC_), averaged across species (_*spp*_). The third and final category—ranking metrics—are common in machine learning and computer vision research, and include top-K accuracy across observations and species, plus mean average precision (mAP).

### Case studies of species and ecosystems

For both case studies, locations were chosen using expert knowledge of the respective species ranges and known occurrences from Calflora. Three non-expert human annotators annotated *Sequoia sempervirens* cover and two annotated *Quercus lobata* cover. To calibrate annotators to the task, each annotator received three NAIP images from 2012 and an assigned cover classification using known species occurrences pulled from Calflora^53^ (**Table S9, Fig. S9, S14C, SM 4.1.2**). Annotations took between 30 minutes to two hours per-case study (depending on the efficiency and familiarity of the annotators with the task) and final cover scores were calculated by averaging annotations per-pixel across annotators.

High-resolution species predictions at 50 m resolution were generated from the CNN networks by convolving the 256 x 256 pixel prediction window with a stride of 50 (**Fig. S8, SM 4.1.1**). It is important to note that the versions of *deepbiosphere, Maxent, RF*, and *MLP* used in these case studies were trained without observations or pseudoabsences from the respective spatial cross-validation band where the case study was located (see darkened band inside California inset in **Figs. 2, S14**). Thus, these models did not see any example images or climate variables from the respective regions at train time, with the nearest training examples located between 9–20 km away from the case studies (**Fig. S4B**). Conversely, the *Inception* baseline was trained using multiple observations from within the parks (specifically using the uniform data split, **Fig. 4A, S11B**).

For the redwoods case study, the 2017 National Park Service (NPS) generalized alliance-level map was used for vegetation comparison^34^, with the class “mature redwoods” mapping to the *Sequoia sempervirens* mature forest alliance, the class “young redwoods” mapping to the *Sequoia sempervirens*-(other) YG alliance, and the class “other vegetation” mapping to all other alliance-level classes present in the study area. Per-pixel labels were determined based on which alliance had the largest area overlap with the pixel’s extent.

For the oaks case study, the USDA Forest Service’s 2018 map of existing vegetation in region 5 was used for comparison^38^, specifically the type 1 regional dominance map with species crosswalked to vegetation type using the vegetation class descriptions from CALVEG zone 7^37^. For the per-species analysis, the species to CALVEG mappings are as follows: *Ceanothus cuneatus*: CC, CQ, EX; *Quercus lobata*: QL; *Bromus diandrus*: HG; *Quercus berberidifolia*: CQ; *Arctostaphylos glandulosa*: CQ, SD, *Adenostoma fasciculatum*: QA, CC, CQ, SS, EX. For each species, pixels were marked as “inside” if said pixel intersected with at least one of the associated CALVEG classes for that species.

### Spatial community change metric

For calculating spatial community change using *deepbiosphere*, we designed a novel edge detection algorithm inspired by common edge detection filters from the field of computer vision. Specifically, the averaged one-neighbor Euclidean norm was calculated per-pixel to generate a map of averaged similarity to neighbor pixels using standard 256 m resolution *deepbiosphere* predictions (**SM 5.1**). This algorithm essentially measures the average distance from a given pixel’s species prediction to all its nearest neighbors’ predictions, summarizing how similar or different a given pixel’s predicted species list is from nearby areas (see **Fig**.**S19** for visual walkthrough). To validate *deepbiosphere’s* spatial community change predictions, we utilized the 2018 Marin fine-scale vegetation map^39^ to calculate the number of vegetation classes intersecting each pixel. Pearson’s *r* between the number of intersecting vegetation classes and spatial community change was calculated using the spatially-corrected modified *t*-test from SpatialPack, using the centroid of each pixel as the coordinates per-sample^54^. A similar comparison to the number of intersecting vegetation classes was performed using the averaged one-neighbor Euclidean norm between the normalized raw NAIP pixel values per-band, upsampled to 256 m resolution.

### Temporal community change metric

For calculating temporal community change using *deepbiosphere*, we used the per-pixel Euclidean distance between *deepbiosphere’s* predicted species probabilities made at two different timepoints (**Fig. S21, SM 5.2)**. This change metric essentially measures the magnitude of per-species change (including both increases and decreases) aggregated between the two timepoints. To validate *deepbiosphere’s* temporal community change predictions for the Rim fire, we compared an independently-generated map of difference in normalized burn ratio (dNBR)^40^ to *deepbiosphere*-generated temporal community change predictions made using 2012 and 2014 NAIP imagery. Pearson’s *r* between temporal community change and dNBR was calculated using the spatially-corrected modified *t*-test from SpatialPack, using the centroid of each pixel as the sample coordinates ^54^. For this comparison, we used 256 m, non-strided species predictions from *deepbiosphere* and dNBR upsampled to 256 m resolution to minimize spatial autocorrelation and ensure the memory-intensive spatially-corrected modified *t*-test could run in sufficient time. A similar comparison to upsampled dNBR was performed using the Euclidean distance between the normalized raw NAIP pixel values per-band upsampled to 256 m resolution.

### Using *deepbiosphere’s* features to generate maps from few examples

To generate downstream maps of vegetation and crop type, we employed *deepbiosphere* as a feature extractor by sequentially feeding in 256 x 256 pixel remote sensing images and extracting the 2,048 outputs from the final layer of the CNN (Linear: 2-23 in **Table S3**, rightmost gray bar in **Fig. 1B**) to use as feature vectors for downstream classification (details in **SM 6.1)**. These feature vectors are numeric vectors that describe high-level features in the images succinctly, and can be used as predictor variables for simpler models such as logistic regression or random forest to perform downstream predictions. We showcased *deepbiosphere’s* feature extraction ability using two datasets, a native vegetation map (Redwoods National and State Park, RNSP), and a crop type map (United States Department of Agriculture, USDA).

For native vegetation mapping, we utilized 1,198 field plots provided by the RNSP^34^; 423 were marked as training plots, 489 as testing plots, and 286 additional plots were not used for fitting the final RNSP vegetation map but were used to fit our downstream models (**Fig. S24B**). The RNSP vegetation map was built with 462 field plots (the additional plots came from locations overlapping the 286 additional plots, which were removed as duplicates in our analysis) while we utilized both the 423 training plots and the 286 additional plots. To minimize overfitting and improve accuracy, and to ensure that there were more training points than training features, we augmented the 709 field plots with an additional 300 points bootstrapped from the test set across ten cross-validation trials, leaving a total of 1,009 field plots for fitting the downstream vegetation classifier. Per-trial, this left 189 plots for testing that neither the *deepbiosphere*-based classifier nor the RNSP algorithm saw during train time. To decrease the downstream classifier’s complexity, we only retained *deepbiosphere* features with a non-zero value for at least one field plot, leaving a total of 955 features for fitting. We compared single label accuracy across three standard downstream classifiers: random forest, multilayer perceptron and logistic regression using the standard scikit-learn version 1.1.1 implementations with default parameters except for logistic regression, which utilized the ‘liblinear’ solver^55^. As a trivial baseline, we also generated random gaussian noise and fitted the same downstream classifiers using the field data. The final map (**Fig. 4B**) was created using the top *deepbiosphere*-based random forest predictor across the ten trials using a 50 pixel stride to achieve a 50 m resolution map. Top-K per-pixel accuracy is calculated by rasterizing the RNPS vegetation map to a 50m-resolution map using rasterio^56^, then comparing the RNPS vegetation class at each pixel to the K most highly predicted vegetation classes from the downstream random forest map. Details of dataset cleaning, vegetation class crosswalking, and the final alliance-level classes used to construct the map are found in **SM 6.2**.

For the cropland type mapping example, we partitioned the 2016 USDA cropland data layer map centered in the San Joaquin Valley^43^ into the same train-test-validation blocks used by ref. ^44^ (**Fig. S28D**). We randomly sampled 1,000 geographic locations within the training blocks, ensuring that locations were at least 128 m away from the boundary edge to minimize examples bleeding over into adjacent blocks (**SM 6.3**). We also randomly sampled 1,000 locations within the held-out test blocks for model comparison. Finally, we removed the top row of training blocks to ensure that no location had been previously seen by *deepbiosphere* during training (**Fig S28A**, maroon band). To compare against a previous state-of-the-art unsupervised deep learning approach called *Tile2Vec*^*44*^, we downloaded the *Tile2Vec*-trained *TileNet* CNN architecture and trained weights from the open Github repository (https://github.com/ermongroup/tile2vec). Cropped NAIP images of the required size (256 x 256 pixels for *deepbiosphere*, 50 x 50 pixels for *TileNet*) were generated for each of the 1,000 training and 1,000 testing locations, and the final layer feature vectors were extracted from each (2,048 features for *deepbiosphere* and 512 features for *TileNet*). Three types of downstream classifiers—random forest, multilayer perceptron and logistic regression—were fitted using these feature vectors and the standard scikit-learn version 1.1.1 implementation with default parameters^55^. We compared accuracies across ten instantiations of each classifier using the 1,000 test examples generated from the unseen test blocks as done in ref. ^44^. To generate a full cropland map of the region, we used the highest performing *deepbiosphere*-based random forest classifier and generated predictions using *deepbiosphere*’s feature vectors extracted from the 60 cm 2016 NAIP imagery and a 50-pixel stride to achieve a 30 m resolution cropland map. Top-K per-pixel accuracy was calculated by comparing the USDA cropland data layer map at each pixel to the K most highly predicted crop classes from the downstream random forest map.

## ADDITIONAL INFORMATION

### Data availability

Data is publicly available through GBIF.org and NAIP^19^. Scripts to regenerate paired image-species datasets and to build *deepbiosphere* are available at github.com/moiexpositoalonsolab/deepbiosphere. Model weights for *deepbiosphere* are available at <huggingface.co/to.be.added>, the full biodiversity dataset is available at <datadryad.org/to.be.added>, and the case study maps are available at <arcgis.com/to.be.added>.

### Author contribution

M.E.-A. and L.G. conceived the project. M.E.-A., M.R. and L.G. developed scripts and conducted the research. L.G. prepared the first manuscript draft. M.E.-A., M.R. and L.G. discussed and edited the final manuscript.

## Acknowledgements

We thank Mateo Rojas-Carulla, Carl Johann Simon-Gabriel, and Detlef Weigel for early discussions or support on this project. We thank Avery Hill, Oliver Bossdorf, Kaycie Butler, the Moi Lab and Goodman Lab for comments and discussion. We also thank TomKat Center for Sustainable Energy as a source of support for this work. We further thank Claudia Engel, Annette Jing, and Sifan Liu for assistance with spatial statistics. We thank Lucas Czech, Gabriel Poesia Reis e Silva, Shannon Hately, Jason Thomas, and Catrina Gillespie for help with annotations. Finally, we thank Ken Morefield and Rosalie Yacoub for assistance with acquiring data.

## Funding statement

This research was funded by the Carnegie Institution for Science and by Microsoft Azure AI for Earth compute credits grant (M.E.-A.). This research was also funded by the NSF Graduate Research Fellowship DGE-1656518 (L.G.), the Stanford TomKat Fellowship (L.G.), and NSF Postdoctoral Research Fellowships in Biology Program under Grant No. 2109868 (M.R.).

Part of the computing for this project was performed on the Memex, Calc, and MoiNode clusters of the Carnegie Institution for Science and the Caltech Resnick High Performance Computing Center.

