## Supplemental Information for "An image is worth a thousand species: mapping plant biodiversity with citizen science, remote sensing, and deep learning"

Lauren Gillespie[1,2,*], Megan Ruffley[1], Moises Exposito-Alonso[1,3,4,*]

[1] Department of Plant Biology, Carnegie Institution for Science, Stanford, California, USA
[2] Department of Computer Science, Stanford University, Stanford, California, USA
[3] Department of Biology, Stanford University, Stanford, California, USA
[4] Department of Global Ecology, Carnegie Institution for Science, Stanford, California, USA

This Supplemental includes:

- Supplemental Text 1-6
- Supplemental Figures S1-31
- Supplemental Tables S1-14

#### Table of contents

**1. Building the dataset** **5**
1.1 Collecting species observations 5
1.2 Creating Joint Observations 5
1.3 Generating the test/train splits 6
1.3.1 Uniform partition of dataset 7
1.3.2 Spatial cross-validation partition of dataset 7
1.4 Collecting Remote Sensing Imagery 7
1.5 Bioclimatic Variables 8
1.6 Resolution Limitations of Bioclimatic Variables 8
1.7 Challenges in Using Open-Source Citizen Science Data 9
**2. Accuracy Metrics** **10**
2.1 Binary classification metrics 10
2.2 Discrimination metrics 11
2.3 Ranking metrics 13
**3. Species Distribution Models** **14**
3.1 Limitations of on-site sampling methods 15
3.2 Convolutional neural network-based species distribution models 15
3.2.1 Loss functions 15
3.2.2 *TResNet* CNN architecture 18
3.2.3 Generating a novel climate + remote sensing CNN 18
3.2.5 *Inception V3* Baseline 19
3.2.6 Selecting the appropriate epoch for evaluation 19
3.3 Climate-only Species Distribution Model baselines 20
3.3.1 Maximum Entropy Baseline 20
3.3.2 Random Forest Baseline 21
3.3.3 Climate-only Multilayer Perceptron baseline 21
**4. Individual species case studies** **21**
4.1 Validating deepbiosphere's predictions for individual species 22
4.1.1 Generating high-resolution species range maps with CNN SDMs 22
4.1.2 Human annotation protocols 22
4.1.3 Redwoods case study 23
4.1.4 Oaks case study 23
**5. Mapping spatiotemporal changes with *deepbiosphere*** **24**
5.1 Detecting spatial community turnover 24
5.2 Detecting temporal community turnover 25
**6. Using *deepbiosphere* for downstream spatial mapping tasks** **26**
6.1 Using *deepbiosphere* as a feature extractor for downstream maps 26
6.2 Generating an alliance-level vegetation map of Redwoods National & State Parks 27
6.3 Generating a crop type map of San Joaquin Valley 28
**SUPPLEMENTAL FIGURES** **29**
Fig. S1 | Comparison of the spatial and temporal resolution of remote sensing imagery vs. bioclimatic data. 29
Fig. S2 | Illustration of building the biodiversity dataset 31
Fig. S3 | Biases present in the dataset and their effects on accuracy 32
Fig. S4 | Partitioning the dataset for cross-validation 33
Fig. S5 | Per-epoch accuracy and loss used to calculate the average optimal epoch for downstream evaluation 34
Fig. S6 | Comparison of per-species accuracy metrics across species 35

Fig. S7 | Species abundance to accuracy relationship 36
Fig. S8 | Generating high-resolution predictions with *deepbiosphere* 38
Fig. S9 | Example images from human redwood labeling task. 39
Fig. S10 | Humans can correctly detect mature redwood groves 40
Fig. S11 | Baseline SDMs cannot detect redwoods in Tall Trees Grove 41
Fig. S12 | *Deepbiosphere*-generated species presence maps for six redwood co-occurring species 42
Fig. S13 | Comparison of co-occurring species presence predictions for associated species 43
Fig. S14 | Predictions of dominant species in the Santa Ynez Valley and Mountains of Southwest California. 45
Fig. S15 | *Deepbiosphere*-generated species presence maps for six chaparral indicator species 47
Fig. S16 | Comparison of predicted presence of indicator species to CALVEG vegetation mapping 48
Fig. S17 | *Deepbiosphere* prediction of spatial community change in northern Marin county. 49
Fig. S18 | Legend for fine-scale Marin vegetation map 50
Fig. S19 | Spatial community change algorithm visual explanation. 51
Fig. S20 | Detecting rapid temporal plant community change after a major California wildfire 52
Fig. S21 | Example of temporal Euclidean distance calculation 53
Fig. S22 | Decrease in predicted *Populus tremuloides* presence before and after the Rim Fire. 54
Fig. S23 | Increase in predicted *Eriophyllum confertiflorum* presence before and after the Rim Fire. 55
Fig. S24 | Classifying alliance-level vegetation in Redwoods National & State Parks using *deepbiosphere* 56
Fig. S25 | Class map for alliance-level vegetation map of Redwoods National and State Parks 57
Fig. S26 | Confusion matrix of *deepbiosphere*-based vegetation classifier on held-out field plots 58
Fig. S27 | Confusion matrix of official NPS vegetation map on held-out field plots 59
Fig. S28 | USDA crop-specific land cover classification case study. 60
Fig. S29 | Color key for cropland datalayer map 61
Fig. S30 | Confusion matrix of *deepbiosphere*-based crop type predictions to USDA cropland data layer 62
Fig. S31 | Confusion matrix of TileNet-based crop type predictions to USDA cropland data layer 63
**SUPPLEMENTAL TABLES 64**
Table S1 | Key metrics of dataset. 64
Table S2 | Model summary of *TResNet* architecture 65
Table S3 | Model summary of *Joint TResNet* architecture 66
Table S4 | Model summary of *InceptionV3* architecture 70
Table S5 | Model summary of *bioclimate MLP* architecture 71
Table S6 | Comparing the performance of different loss functions on uniform data split 72
Table S7 | Comparison of *deepbiosphere* to baseline SDMs and previous deep-learning-based approaches 75
Table S8 | Comparison of *deepbiosphere* to baseline SDMs across spatial cross-validation bands. 76
Table S9 | Site details for individual species case studies and human annotation experiments 77
Table S10 | Accuracy of various SDMs for predicting previously unseen occurrences in northern California 78
Table S11 | Accuracy of *deepbiosphere*-based vegetation type prediction compared to official map 80
Table S12 | Per-class accuracy of *deepbiosphere* vegetation type prediction compared to official map 81
Table S13 | Comparison of *deepbiosphere* crop type accuracy to state-of-the-art unsupervised learning method 82
Table S14 | Per-crop accuracy of *deepbiosphere* compared to state-of-the-art unsupervised learning method 83
**SUPPLEMENTAL REFERENCES 84**

### 1. Building the dataset

The methods for dataset collection were inspired from ref [1] and broadly consist of linking a set of species observations linked to corresponding remote sensing imagery. How the citizen science observations and remote sensing imagery were collected is detailed below.

#### 1.1 Collecting species observations

We collected observations from kingdom *Plantae* using *GBIF.org* from the years 2015-2022[2]. Only records observed by humans with a coordinate uncertainty radius of less than or equal to 120m with no flagged geospatial issues were taken from within the state of California. Nearly all of the subsequent observations were public observations uploaded using the iNaturalist app[3]. Any person with a smartphone and who has downloaded the app can upload observations to iNaturalist, meaning that the observations used in this dataset were collected by many thousands of citizen scientists with a wide variety of backgrounds. Accordingly, some observations may be mis-identified.

To minimize mis-identification, only research-grade observations with taxon identifications from at least two community members were included in the dataset[4]. That being said, mis-identified observations or mis-located observations can still slip past these filters, making this data especially challenging to work with[5,6]. However, GBIF takes steps to resolve major mis-identification events between closely related taxa[6]. Finally, a recent case study from San Clemente island in California showed that all examined iNaturalist observations with a positional error of < 10m as listed in GBIF were within 270m of the corresponding species detected from remote sensing imagery[5]. Extrapolating to this dataset, it's reasonable to assume that the <120m positional uncertainty filter we used would place the vast majority of observations within the linked remote sensing image, therefore preserving the geographic relationship between the observed species and the image.

In total we downloaded a total of 912,380 plant observations of 5,193 unique plant species[2]. We further filtered observations to only include vascular plants, which we define vascular plants as all plants in the taxonomic classes of Gnetopsida, Liliopsida, Lycopodiopsida, Magnoliopsida, Pinopsida, Polypodiopsida, Lycopodiopsida, and Ginkgoopsida. We also removed duplicate observations of the same species within a 150m radius, removed species that contain all observations located within a 256m radius, and were not geographically located within the Global Administrative Area boundary of California, or were missing climate or NAIP imagery data. To increase the density of observations in the dataset, we used neighbor imputation to add any other species observed within an overlapping 256m radius to a given observation (**Fig. S2**). We finally removed any species that had fewer than 500 total observations in the dataset after neighbor imputation, leaving us with a total of 652,027 observations of 2,221 unique plant species (**Table S1**).

#### 1.2 Creating Joint Observations

To create the joint species occurrence dataset, we used neighbor imputation of overlapping observations. Two individual observations from the species observations dataset are considered to be locally overlapping if the Euclidean distance between the latitude and longitude of the two points is less than or equal to 256 m. While this technically means that some neighbor observations may not be geographically located within a neighbor's 256 × 256 pixel image, said resolution is on par with accepted spatial scales in theoretical biogeography and empirical community ecology, which have shown that biotic species interaction networks between individual plants can reach scales of thousands of square meters, and both biotic and shared land use features are thought to drive site-level plant distribution[7–9]. Therefore, although there is no strict guarantee that two observations lie within the extent of their overlapping observations' respective NAIP imagery, strong ecological theory supports that the two species may be influencing each other's co-occurrence and subsequent observed co-occurrence.

The reason that we chose to create a joint dataset is twofold. First, there is theoretical evidence to support that biotic interactions (the interactions of two living species both directly and indirectly) are a strong driving factor in the distribution of species at a variety of scales[8] and that overlap data can be seen as a partial observation of these biotic interactions. Second, many species in our dataset have few observations (**Fig. S3A**) which can make it impossible to learn an accurate representation of the species' distribution. However, oftentimes these rarely observed species will inhabit similar habitats to much more commonly observed species. Therefore, building a multi-species observations dataset may enable better modeling of those rare species using shared habitat signatures learned for the more common, overlapping species.

Along with providing overlap data, we also provide higher taxonomic information per-image. Concretely, the species, genus, and family of all overlapping species is utilized for training. Our rationale was that the phylogenetic history embedded within the taxonomic hierarchy of species should also encode a shared ecological niche space for some taxonomic groupings.

#### 1.3 Generating the test/train splits

In order to properly validate and compare models, we split the dataset into multiple partitions. Best practices to ensure the reproducibility of machine learning models is to have a train-test-validation split, ideally with the validation data coming from a separate acquisition process to provide as robust a test as possible. To test models with statistical power, such validation sets on the order of thousands of observations would be necessary. Furthermore, to test our models appropriately, we require test and validation sets that are at least 1,300m away from all training set examples, to prevent data leakage due to spatial autocorrelation in the coarse resolution bioclim data used by baseline models. Next, since the CNN predicts at 256m resolution, observations with low geographic uncertainty are a must. Finally, since citizen science observations tend to be common on publicly accessible land and near major access routes, candidate locations for taking validation observations tend to be located on private, inaccessible land or in very remote locales that are exceedingly difficult to reach on foot. Understandably, finding or collecting such a validation dataset is an exceedingly challenging task and is left for future work. Having those caveats in mind, to robustly test the models with the data we have, we designed two types of test/train splits to pick hyperparameters and test extrapolation accuracy: The first type—the uniform split—was used primarily for choosing an optimal loss function and learning rate, while the second type—spatial cross-validation—was used to validate these choices and test model extrapolation ability.

##### 1.3.1 Uniform partition of dataset

The first partition, which was used for loss function comparison and learning rate selection, was generated by uniformly randomly selecting observations from across the state to be part of the test set and to test interpolation ability (**Fig. S4A**). We chose points uniformly across the state to maximize the number of unique climates where models would be evaluated on. To ensure the independence of training and testing set data due to spatial autocorrelation, we added all overlapping observations to the test set to guarantee that none of the remote sensing images and observations in the test set were present in the training set. In order to ensure that there was no data leakage between the test and train set, only observations which were more than 1,300 m away from any other non-overlapping observation were included. We chose an exclusion radius of 1,300 m because the climate variable raster pixels converted from arc-seconds to meters can have a diameter of up to 1,200 m, so any test set observation within that distance to any observation in the train set would have an identical input value as some observations used during fitting, resulting in data leakage. Ultimately 1.88% of the dataset was set aside for testing using this method. The test dataset has relatively few observations compared to a traditional 80/20% test/train split because the *iNaturalist* observations are very spatially heterogeneous, and tend to be very clustered, meaning that there are few observations that are sufficiently far enough away from observations used for training to be included in the test split.

##### 1.3.2 Spatial cross-validation partition of dataset

An alternative cross-validation to the uniform train-test split was also designed to test the extrapolation ability of models. Specifically, we conducted a ten-fold spatial holdout block validation by partitioning California into ten one-degree latitudinal bands (**Fig. S4B**). Training points within 1,300 m of the test band were removed to prevent data leakage as discussed above (For models utilizing pseudo-absence points, all pseudo-absence points within the test bands were removed to ensure a fair comparison to presence-only models). Ultimately, the percentage of test points per-spatial block ranged from 1.40-25.35% of the entire dataset. Since training ten models from scratch takes a considerable amount of time and resources, we only compared the final *deepbiosphere* model and all baselines. The *Inception* baseline CNN model specifically was not cross-validated as it had poor results on the uniform split of the dataset (**Table S8**), and since it is significantly larger than our *TResNet* CNN architecture, thus training it takes considerable time.

#### 1.4 Collecting Remote Sensing Imagery

The remote sensing data used for this project is from the National Agriculture Imagery Program (NAIP) aerial imagery from 2012 (ref. [10]). We chose this dataset because it contains sun angle-corrected orthophotography data collected during the leaf-on growing season with guaranteed < 10% cloud cover at 1 m-resolution (see ref. [11] for a comprehensive overview of NAIP data). We utilized Microsoft's Planetary Computer for dataset access. Briefly, in this dataset the remote sensing images were generated by cropping a 256 × 256 pixel image from the raw NAIP .tiff files centered at the pixel corresponding to the latitude and longitude of each observation (see **Fig S2** for a visual explanation of data collection). Each pixel in the underlying NAIP data is 1m x 1m in resolution (**Table S1**) and each image has four bands corresponding to wavelengths in the red, green, blue, and infrared regions of the light spectrum.

Our decision to work with four-band RGB-Infrared remote sensing data instead of many-band Landsat data or full-band hyperspectral data was motivated by data accessibility and resolution to capture biological patterns. Although Landsat data is collected worldwide, it has a pixel resolution of 30 m per-pixel, while NAIP data has a 1m-60cm pixel resolution (depending on the year of acquisition). Both Landsat and NAIP are sub-kilometer resolution, but the higher resolution of NAIP data specifically may better capture important local factors for plant distribution modeling such as individual tree crowns, land use, and biotic interactions. Further, the four color and infrared bands (RGB-I) of NAIP also contain the same information as derived vegetation products such as NDVI[12]. Other hyperspectral remote sensing products with a similar level of resolution to NAIP—such as the 224-band, full-spectrum product AVIRIS—have limited coverage at national scale, not even covering all of California currently. Finally, four-band (RGB-I) remote sensing products of a similar resolution are available from private companies like Maxar Technologies and Planet Labs across the entire globe with weekly to nearly daily acquisitions, although these products are not publicly accessible. Therefore, the four-band NAIP imagery provides an optimal mix of spatial resolution and availability to appropriately model plant communities at a theoretically sound scale with the ability to scale these techniques to potentially the rest of the globe.

#### 1.5 Bioclimatic Variables

Beyond remote sensing imagery, we also used WorldClim 2 bioclimatic environmental variables at 30 arc-second resolution[13] (~ 1 km depending on latitude). We chose to use WorldClim because the 19 bioclimatic variables are a popular choice for bioclimatic SDMs and have been used to generate many hundreds to thousands of SDMs. Using WorldClim as the baseline climatic environmental variables to compare our remote sensing-based approach allows for a fairer comparison to previous methods built using similar features.

#### 1.6 Resolution Limitations of Bioclimatic Variables

While downscaling and interpolation techniques exist that allow one to scale these bioclimatic variables to 250-100 meter resolution[14,15]—a similar resolution to our methods—these downscaling and interpolation methods are inherently limited in the amount of novel information they can provide, and introduce a new source of potential bias[16]. Furthermore, there has been evidence that SDMs trained using coarser-grained climate data overestimate some montane species' tolerance to changing climate compared to meter-resolution local scale information[17]. This discrepancy can be easily seen in **Fig. S1** which illustrates the difference between the remotely sensed NAIP imagery and Bioclim in northwest California. One can easily distinguish the various grassland-forest transitions along with land use differences from agriculture. This local information is simply not detectable from the projected 7th band of Bioclim data shown. The 30 x 30 arc-second pixels of the bioclimatic rasters are simply too coarse-resolution to capture the local variations between the various communities in this image, as evidenced by the fine scale vegetation map for the region (**Fig. S1B**). Furthermore, as discussed in Section 1.4, from ecological theory it is expected that fundamental drivers of species distribution should be different at local versus site scale, with bioclimatic data driving local scale distribution versus topographic and land use information driving site scale distribution[9]. Thus it is to be expected that the information contained in both sources to be radically different.

#### 1.7 Challenges in Using Open-Source Citizen Science Data

Creating citizen science-based species distribution models is challenging due to the unevenness of number of observations per species and data quality issues. Despite our data filtering, which included minimum data thresholds for species to be included in the analyses, we still see many dataset imbalances; for example, 20% of species have 1,000 images or less while only 27% of species have 10,000 or more. In addition, 37,300 images contain only one species while only 61,386 images have >100 species, reaching the expected size of a full species checklist for a given 256 m radius area. Moreover, species labels are presence-only, meaning that in most cases, the absence of a species label in an image does not guarantee true absence from the ecosystem, only that it has not been observed on iNaturalist at that location. Overall, this leads to a highly imbalanced dataset, with some habitats and some species underrepresented and hard to study (**Fig. S3**).

Specifically, our dataset of observations suffers from three main types of biases. First, there's spatial bias, in that observations are not uniformly distributed across the landmass of California (see **Fig. 3C, 3D**). Citizen scientists can only take observations where they can themselves access species, which restricts most observations to publicly owned land and convenience plays a large factor in the distribution of observations, with more observations coming from ecoregions with a higher population or more public parks, such as the Southern California Chaparral or Coast Range ecoregions (**Fig. S3D**).

Second, observations from casual users tend to show density bias (**Fig. S3B**), where many observed locations have few other overlapping species reported, while a few observations have many overlapping species reported. Oftentimes, this is a result of observers noticing and documenting a particularly salient individual of a species, like when a specific wildflower is in bloom in the spring. However, rarely will users upload all the plant species they may find in a given small area, meaning that the majority of species present at any given observation location in our dataset are unreported. We refer to these present but unobserved species as *pseudo-absences* in our data. The high pseudo-absence rate of our dataset also means that we cannot consider these species occurrences to represent full species checklist data at each site in our dataset, meaning that the co-occurrence network of each site in our dataset is partially observed, again adding extra challenge to the machine learning task.

Third, as mentioned in Section 1.3, the dataset is also long-tailed in the number of observations per-species, with many species possessing few observations while a few possess a large number of observations in the dataset (**Fig. S3A**). Unfortunately, this "commonness of rarity" is a known phenomenon in plants' distributions and is an expected phenomenon for plant observations[18]. This class imbalance can be problematic for classic machine learning algorithms, since standard accuracy metrics become less informative and models can simultaneously suffer from both under- and over-fitting across classes[19]. However, it should be noted that these challenges are not unique to this dataset alone[20–22], thus algorithms successfully able to learn a generalizable representation of this data should be of interest to both the species distribution modeling and more general machine learning fields.

Despite these challenges, citizen science provides an opportunity to utilize deep learning methods that may require hundreds of thousands to millions of observations to train. Further, the wide array of learning functions and training techniques for deep learning models can still enable the learning of useful patterns even with this incomplete dataset.

### 2. Accuracy Metrics

The work in this paper spans many disparate research communities (i.e. deep learning, computer vision, biogeography) which use a variety of different accuracy metrics. To that note, we report twenty separate accuracy metrics in an effort to be as comprehensive as possible. It should also be noted that what absolute value of an accuracy metric constitutes a "good" model also varies across communities, tasks, and datasets. Just because a model does not have a near-perfect accuracy metric does not necessarily mean it's not a good nor useful model. Depending on the use case, even models with low accuracy metrics can lead to useful insights.

#### 2.1 Binary classification metrics

Binary classification metrics are a very common set of metrics used to compare yes/no binary prediction tasks (or e.g. whether a species is or is not present at a given location). For binary classification-based metrics, probability predictions must be converted to a binary presence / absence output using a threshold value. There is vigorous debate within the SDM community on the proper threshold to pick for presence versus absence[23,24], but for consistency with the computer vision community and the fact that our multi-label domain makes optimal threshold determination non-trivial, we chose to threshold all probabilities ≥0.5 as present and <0.5 as absent. Since our dataset is a multi-label dataset, there are two main ways one can partition presence-absences for accuracy calculation: **across species** and **across images**. We report only per-species metrics in the main text as it provides a richer explanation of the models' ability across species. We calculate these metrics in a multi-label setting, meaning that we use the imputed overlapping species mentioned above in the ground-truth labels when calculating all binary classification-based metrics.

There also exist many different metrics for binary classification, but in this work we focus on four common ones: *precision*, which measures how many species predicted to be present were actually present; *recall*, which measures how many of the true species present are predicted as present; *F1*, which is the harmonic mean of precision and recall and represents a conservative mean of the two (i.e. is more affected by low values); and *accuracy* which simply measures the percent of correctly identified examples of present species. For each metric, both binary and multi-class versions exist, along with single-label and multi-label. In this work, we report the multi-label, multi-class versions of each metric except for accuracy, which is reported as multi-class, single-label. To clarify this technicality, we refer to accuracy as *single-label accuracy* in this work to signal that for each example the model can either be right or wrong, as there is only one species tested at a time.

In the multi-label, multi-class setting, there are multiple axes by which one can aggregate the chosen statistic: the first is by species, which we refer to as *per-species*; the second is by example image, which we refer to as *per-image*. For all metrics, $S$ is the number of unique species in the training split of the dataset, $N$ the number of images in the training split of the dataset, $\bar{y}$ is the multi-label ground truth presences and absences of each species for each image, $y_s$ is the single-label original species associated with each observation, $\hat{y}$ is the SDM's predicted binary present / absent list for each species and each image using a $0.5$ threshold, $tp$ is true positives, $fn$ is false negatives and $fp$ is false positives (NB: true negatives are unknown). Per-species metrics were calculated using scikit-learn version 1.1.1 (ref. [25]) and per-image metrics were implemented by ourselves (see open Github repository for implementation: github.com/moiexpositoalonsolab/deepbiosphere).

$$\text{per-species recall} = \frac{1}{S}\sum_{i=1}^{S}\frac{tp_i}{tp_i + fn_i}$$

$$\text{per-species precision} = \frac{1}{S}\sum_{i=1}^{S}\frac{tp_i}{tp_i + fp_i}$$

$$\text{per-species F1} = \frac{1}{S}\sum_{i=1}^{S}\frac{2 \cdot Rec_i \cdot Prec_i}{Rec_i + Prec_i}$$

For per-image accuracy metrics, we used the definitions as outlined in ref. [26]. We implemented these metrics in Python ourselves.

$$\text{per-image precision} = \frac{1}{N}\sum_{i=1}^{N}\frac{|\bar{y}_i \cap \hat{y}_i|}{|\hat{y}_i|}$$

$$\text{per-image recall} = \frac{1}{N}\sum_{i=1}^{N}\frac{|\bar{y}_i \cap \hat{y}_i|}{|\bar{y}_i|}$$

$$\text{per-image F1} = \frac{1}{N}\sum_{i=1}^{N}\frac{2 \cdot Rec_i \cdot Prec_i}{Rec_i + Prec_i}$$

Finally, for the single-label accuracy, we use the standard multi-class definition of the fraction of examples where the correct species observed at that location was predicted as present. This metric was calculated using our own custom implementation.

$$\text{single-label accuracy} = \frac{\sum_{i=1}^{N} y_{i,s} = \hat{y}_{i,s}}{N}$$

#### 2.2 Discrimination metrics

Binary classification metrics do come with some drawbacks, mainly that the choice of what threshold value to use for determining whether a species is present can have an outsized impact on the accuracy of a model[27]. In order to take into account the effect of thresholds, discrimination metrics measure binary classification ability across a wide range of thresholds in order to both control for the effect of threshold choice and to determine what might be an optimal threshold of presence. Rather than set an arbitrary threshold, they observe how well the model is able to predict across the range of predicted probabilities for that class to see if a higher threshold means more species are correctly predicted without adding in too many false positives. For discrimination metrics, we report the area under the receiver operating characteristic curve ($AUC_{ROC}$) averaged across species, average area under the precision-recall curve ($AUC_{PRC}$) averaged across species. We again use multi-label, neighbor imputed ground truth presences and absences when calculating discrimination-based metrics. We use scikit-learn version 1.1.2 for all discrimination metrics, which utilizes the trapezoidal integration to calculate area under the curve[25]. $tp(\hat{y_s}, i)$ refers to the number of true positive predictions of species $s$ in $\hat{y}$ when using $i$ as the threshold for predicted presence for species $S$, while $fp(\hat{y_s}, i)$, and $fn(\hat{y_s}, i)$ are the same for the number of false positives and false negatives, respectively. $P(\bar{y_s})$ is the true number of actual presences in the ground truth for species $S$, which can also be written as $\sum_1^N \bar{y_s}$ and $N(\bar{y_s})$ is the true number of actual absences in the ground truth data for species $S$, which can also be written as $\sum_1^N 1 - \bar{y_s}$.

$$\text{average } AUC_{ROC} = \frac{1}{S}\sum_{s=1}^{S} \int_{i=\min(\hat{y_s})}^{\max(\hat{y_s})} \frac{1}{2} \cdot \Delta TPR(\hat{y_s}, i, \bar{y_s}) \cdot \Delta FPR(\hat{y_s}, i, \bar{y_s})$$

$$\text{average } AUC_{PRC} = \frac{1}{S}\sum_{s=1}^{S} \int_{i=\min(\hat{y_s})}^{\max(\hat{y_s})} \frac{1}{2} \cdot \Delta Prec(\hat{y_s}, i) \cdot \Delta Rec(\hat{y_s}, i)$$

$$\Delta Prec(\hat{y_s}, i) = \frac{tp(\hat{y_s}, i+1)}{tp(\hat{y_s}, i+1) + fp(\hat{y_s}, i+1)} - \frac{tp(\hat{y_s}, i)}{tp(\hat{y_s}, i) + fp(\hat{y_s}, i)}$$

$$\Delta Rec(\hat{y_s}, i) = \frac{tp(\hat{y_s}, i+1)}{tp(\hat{y_s}, i+1) + fn(\hat{y_s}, i+1)} - \frac{tp(\hat{y_s}, i)}{tp(\hat{y_s}, i) + fn(\hat{y_s}, i)}$$

$$\Delta TPR(\hat{y_s}, i, \bar{y_s}) = \frac{tp(\hat{y_s}, i+1)}{P(\bar{y_s})} - \frac{tp(\hat{y_s}, i)}{P(\bar{y_s})}$$

$$\Delta FPR(\hat{y_s}, i, \bar{y_s}) = \frac{fp(\hat{y_s}, i+1)}{N(\bar{y_s})} - \frac{fp(\hat{y_s}, i)}{N(\bar{y_s})}$$

One major drawback to this approach is that it does not measure the calibration of an SDM's predicted probabilities well, meaning that a model which has extremely low predicted probabilities can still nevertheless have a high $AUC_{ROC}$ if within its range of predicted probabilities it has good sensitivity and specificity for that class. This means that it's possible to have a model that never predicts a species as present with the standard presence/absence threshold of 0.5 yet still has a high average $AUC_{ROC}$. Furthermore, tuning the presence/absence threshold using the ROC curve for such models is non-trivial, since the derived optimal threshold will likely be different across species. To correct for this, we also introduce a calibrated area under the curve score where the chosen thresholds are linearly interpolated values between 0 and 1. We use a trapezoidal approximation of area under the curve, utilizing scikit-learn's implementation of area under the curve with 50 uniformly spaced probabilities between 0 and 1 (ref. [25]).

$$\text{calibrated avg. } AUC_{ROC} = \frac{1}{S}\sum_{s=1}^{S}\int_{i=0}^{1}\frac{1}{2}\cdot\Delta TPR(\hat{y_s}, i, \bar{y_s})\cdot\Delta FPR(\hat{y_s}, i, \bar{y_s})$$

$$\text{calibrated avg. } AUC_{PRC} = \frac{1}{S}\sum_{s=1}^{S}\int_{i=0}^{1}\frac{1}{2}\cdot\Delta Prec(\hat{y_s}, i)\cdot\Delta Rec(\hat{y_s}, i)$$

#### 2.3 Ranking metrics

Compared to binary classification and discrimination metrics, ranking metrics focus solely on how high a given species is ranked by probability of presence compared to other species in the same image. These ranking metrics suffer from the same limitations as the aforementioned discrimination metrics in that they only compare accuracy of probabilities in a relative sense, rather than the absolute. However, they are the most common within the deep learning and computer vision communities, so we choose to report them here for completeness. The first set of ranking-based metrics we report are top K accuracy metrics, a set of single-label metrics. These metrics measure how many times the correct species was correctly predicted within the top K highest-ranked species within a given image (where, for example, K=5 would be the 5 species with the highest probability of presence). Much like the binary classification metrics, top K accuracy can be calculated across images and also across species (which we refer to as $\text{TopK}_{img}$ and $\text{TopK}_{spp}$, respectively), and we report both in the supplemental. However, top K accuracy across species is considered to be a better metric of an SDM's ability to distinguish present species, as it corrects for sampling imbalances across species[1,6].

Machine learning papers oftentimes report top-1 or top-5 accuracy, but given our task is an inherently multi-label one, we choose to report with a larger K than normally seen in computer vision projects with many possible labels, as the expected number of unique plant species at the local scale varies anywhere from five to one hundred and thus on average we are most interested in the composition of these top five to one hundred species. To that note, we report both $\text{TopK}_{img}$ and $\text{TopK}_{spp}$ for K = 1, 5, 30, and 100. Along with top K accuracy, both top K recall and precision also exist but are far less commonly reported and so we do not report them here. Finally, as a single-label metric, these top K accuracy metrics do not use or capture any information about an SDM's ability to correctly label overlapping species, making them less useful for judging an SDM's ability to capture co-occurrence patterns correctly. We implement this metric in Python using the definitions from ref [1]. Here, $\text{rank}(j, \hat{y}_i)$ is defined as the rank of species $j$ observed in image $i$ from the sorted list of probabilities $\hat{y}_i$ predicted by the SDM, $\text{rank}(j, \hat{y}_i) = \|\{k : \hat{y}_{i,k} \geq \hat{y}_{i,j}\}\|_0$.

$$\text{TopK}_{img} = \frac{1}{N}\sum_{i=1}^{N} Acc(\hat{y}_{i,j}, K)$$

$$\text{TopK}_{spp} = \frac{1}{S}\sum_{i=1}^{S} Acc_{spp}(\hat{y}_{i,j}, i, K)$$

$$\text{where } Acc_{spp} = \sum_{m=1}^{N_i} Acc(\hat{y}_{m,j}, K) \text{ and where } Acc = \begin{cases} 1 \text{ if } \text{rank}(j, \hat{y}_i) \leq K \\ \\ 0 \text{ otherwise} \end{cases}$$

Finally there does exist a commonly-used multilabel ranking-based metric called mean average precision (mAP). This metric is also sometimes referred to as label ranking average precision (LRAP). mAP calculates how highly each species is correctly ranked along with how many other present species are ranked higher, averaged across species. mAP is the multilabel version of the mean reciprocal rank metric (MRR) commonly used in document retrieval.

$$\text{mAP} = \frac{1}{N} \sum_{i=1}^{N} \frac{1}{\|\bar{y}_i\|_0} \sum_{j:\bar{y}_{i,j}=1}^{J} \frac{\|L(\hat{y}_{i,1:j})\|_0}{\text{rank}(j, \hat{y}_i)} \text{where } L = \begin{cases} 1 \text{ if } \hat{y}_{i,k} \geq \hat{y}_{i,j} \\ 0 \text{ otherise} \end{cases}$$

### 3. Species Distribution Models

Species distribution models (SDMs) describe how a species is distributed across a given geographic extent. Oftentimes this is accomplished by modeling how the predicted presence of a species across a landscape varies spatially in response to a set of ecologically-meaningful variables. SDMs broadly fall into two rough categories: process-based and correlative. Process-based models attempt to derive fundamental equations based on processes or mechanisms governing a species distribution and build a model of likelihood of occurrence (e.g. dispersal ability, growth rates, demographic characteristics, etc.). Correlative models attempt to infer a species' geographic extent by correlating its known occurrences with a suite of relevant environmental variables and projecting likelihood of occurrence from said correlated variables. However, these definitions are not dichotomous nor mutually exclusive, meaning correlative models may indeed capture some process-based mechanisms intrinsically in their modeling procedure[28].

Further distinction can be made between single species and joint species SDMs, with the former only modeling one species at a time and the latter attempting to model multiple species' distributions simultaneously. Joint SDMs can be further subdivided based upon at what point in the modeling process the species were aggregated. Some joint SDMs are really aggregation of individual SDMs, with the predicted species' presence joined during *post-hoc* analysis[29], while other joint SDMs model all species simultaneously throughout both the model fitting and analysis steps[30]. For future clarification, when referring to a *joint SDM*, we are referring to the latter process.

A final important distinction between different types of SDMs is between functional niche models and realized niche models. The functional niche of a species traditionally was defined as the set of environments where a species individually can sustain itself, while the realized niche is the set of environments where a species can sustain itself in the presence of competition from other biotic sources[31]. Contemporary niche theory has strengthened these definitions to include dispersal dynamics, growth rates, and biotic interactions, all vital processes to a species' dispersal[32]. However, many modern uses of the terms use a simpler, less precise definition of niche that simplifies the fundamental niche to where a species *can* occur and the realized niche to where a species *does* occur[33,34]. Our approach falls somewhere in between, where for some species, like large redwoods, the realized nice is likely being modeled through direct detection of redwood canopy signatures, while for smaller less-observable species, like redwood sorrel, the modeling process is closer to the functional niche than realized.

### 3.1 Limitations of on-site sampling methods

While the best metric of species presence and current biodiversity comes from on-the-ground observation of species, generating comprehensive checklists of species presence at high resolution across large geographic extents is generally infeasible. For example, in ref. [35], rarefaction curves of species richness were generated at various grid sizes and using estimates from this analysis of over 1.5 million plant observations, an estimated 1,000 observations would need to be taken per grid scale at a 15 km resolution to reach a point where the rarefaction curve begins to plateau, with many cells requiring upwards of a magnitude more samples due to California's extremely heterogeneous and endemic plant communities. Using the 1,000 observation estimations, an conservative estimated uniformly sampled 2 million observations would be needed to estimate species richness at 15km spatial blocks, with rapidly increasing numbers of samples as the desired resolution increases. All in all, it would be infeasible to try and perform the same species checklist curation that *deepbiosphere* can perform using onsite methods.

### 3.2 Convolutional neural network-based species distribution models

Convolutional neural networks (CNNs) are a popular machine learning model that have been adapted to successfully model a wide variety of complex, real-world image-related tasks, from image classification to deciphering handwriting[36,37]. Their widespread success lies in their ability to learn and extract arbitrary patterns and features from images without any human input, allowing them to detect and exploit only the most relevant visual features of an image for a given task. This ability to intrinsically learn the most relevant features of an image makes them very flexible for use across a wide variety of image-related tasks and a wide range of image media, from medical imagery of cells to satellite imagery[38,39]. In this work, we seek to predict the presence of thousands of plant species from relevant features found in the local aerial imagery of a location, a task that CNN models are well-designed to perform. While there have been other deep neural network architectures applied to similar visual tasks, for the task of *image classification* (the assigning of a discrete label to a given image) and balancing model size and speed, the classic residual convolutional neural network (ResNet) family of architectures is sufficient[40]. Given that standard classification-based CNNs are more simple, faster to train, and more accessible to a wider array of downstream users than larger and more memory-intensive architectures, we chose to use standard ResNet architectures for this work.

##### 3.2.1 Loss functions

The loss, which determines how correct the CNN model's prediction is for a given example, is task-dependent and there are many different choices for a given task type. In this work, we frame our problem as a classification task, where the goal is to classify each image into one of N classes. We also frame our problem as a multilabel task, which extends the above definition to classify each image into K of $S$ classes (where K is the number of present species in the observation). While many CNNs have been developed for image classification, the vast majority of these architectures have been designed for *single label classification*, where for each image exactly one class should apply; for example, each image may either have a dog or a cat, but an image is never expected to have both. However, what makes our dataset both unique but challenging is that it provides occurrences of all overlapping species in a given image, making it a *multilabel* dataset (see Section 1.2 for more details), since each image is associated with anywhere from one to nearly one hundred overlapping plant species within 256 × 265 m squares. By single-label, we mean that for each training example only the original species observed at that location is included as a positive label, with all imputed neighbors being ignored. When using a multi-label loss, we also include all imputed neighbor species as positive labels in each training example. In all notation $\hat{y}$ refers to the raw outputs from a neural network, $\bar{y}$ refers to the single- or multi-hot vector of known species presence in that example, where 0 means a species is absent and 1 means a species is present, and $S$ is the number of unique species in the dataset.

The most commonly used loss function for training classification CNNs in a single label setting is cross-entropy loss (CE).

$$\text{CE} = -L_+ - L_- \begin{cases} L_+ = \bar{y} \cdot log(f(\hat{y})) \\ L_- = (1-\bar{y}) \cdot log(1 - f(\hat{y})) \end{cases} \quad \text{where} f(\hat{y}_i) = \frac{e^{\hat{y}_i}}{\sum_j^S e^{\hat{y}_j}}$$

It should be noted that softmax-based losses like CE loss were designed for classification of single-label datasets like the ubiquitous ImageNet by modeling a probability distribution across labels[37] (i.e. the sum of probabilities across all possible labels must be 1). However, this approach does not match the multi-label nature of our task well, as in each image the probabilities across species should be independent so that the presence of one species in an image does not imply a decrease in presence of other species. Furthermore, when using a softmax-based transformation of model outputs, probabilities of an individual species are no longer directly comparable across individual observations on account of the lack of independence across labels, and thus is not a valid transformation for calculating binary classification metrics. Nevertheless, to compare to previous work in ref. [41], we still report accuracy metrics for models trained with CE loss using the softmax transformation.

In the multi-label setting, the classic loss used for training CNNs is binary cross entropy (BCE) loss:

$$\text{BCE} = -L_+ - L_- \begin{cases} L_+ = \bar{y} \cdot log(f(\hat{y})) \\ L_- = (1-\bar{y}) \cdot log(1 - f(\hat{y})) \end{cases} \quad \text{where} f(\hat{y}_i) = \frac{1}{1 + e^{-\hat{y}_i}}$$

BCE loss can be intuitively interpreted as training the neural network to maximize the conditional log-likelihood of species presence and thus the network's predictions can be interpreted as the likelihood of species occurrence in any arbitrary image. However, our dataset exhibits two important types of imbalance that make this standard loss not ideal for our task. First, our dataset exhibits strong observation imbalance, with few species possessing many observations and many species possessing few observations (**Fig. S2A**) which is problematic since the standard BCE loss formulation assumes an equal number of observations per-class in the dataset. The second is label imbalance, or the imbalance in the number of present vs. absent species per-image, a byproduct of using citizen science observations with incomplete coverage of all species present in a given 256 × 256 m area (see Section 1.7 for details). Most observations have fewer than five species present in the observation (**Fig. S2B**), and very few observations reach close to the expected true number of plant species present in a given 256m radius. Therefore, in most cases only one-to-one hundred classes from the 2,221 species are present in a given observation and we must assume all locations contain some pseudo-absences in $\bar{y}$. Since BCE loss assumes all absences are true absences, yet pseudo-absences are guaranteed to be present in the dataset, a loss calculation whose main contributions are from absences points is not ideal for our task.

Therefore, we also considered two other losses that handle the contribution of the negative class differently. The first is a recent multi-label classification loss variant of focal loss called asymmetric focal loss (ASL). This loss was explicitly designed and optimized for multilabel tasks by upweighting the loss contributions of the present classes without eliminating the contribution of the absent classes entirely[42]. It does so by breaking the standard BCE definition down per-class, depending on whether the class is present or absent in the observation. The contribution of the absent versus present classes is then differentially applied to the loss using the hyperparameters $\gamma+$ and $\gamma-$. By setting $\gamma > 0$, the loss contribution of all absent classes will be scaled down, decreasing their contribution to the overall loss. Furthermore, very easy negative examples which the network assigns low probability ($\hat{y}_i \ll 0.5$) will be exponentially down-weighted, creating a "soft thresholding" effect.

The loss contribution of absent classes can be further reduced for easy negatives through the addition of "hard thresholding" (the function $p_m$) which fully discards the loss contribution for absent classes with predicted probability below a tunable threshold, $m$. One can think of this as "throwing away" the loss contribution of very easy classes ($\hat{y}_i \lll 0.5$). Furthermore, the shape of loss function is such that very hard negative samples ($\bar{y}_i = 0$ when $\hat{y}_i \approx 1$) have a down-weighted loss contribution as well (see Fig. 3 of Ben-Baruch et. al 2020 for specifics). This corresponds to the scenario where an observation is "mis-labeled," which in our dataset would correspond to a location missing an observation of a clearly present species, thus minimizing the negative effect of density bias seen in our dataset. Finally, the values of $\gamma$ can be dynamically adjusted during online training to maintain symmetry between the probability contribution of absent versus positive classes, to ensure that the loss contribution of negative samples does not "overwhelm" the contribution of the positive classes. Overall, these benefits fit our dataset well.

This soft and hard thresholding is somewhat analogous to the sampling of pseudo-absence points, a required step for the maximum entropy modeling of MaxEnt, but importantly the hyperparameters are not dependent on the spatial extent of a given species, as are the pseudo-absence sampling ranges of MaxEnt, and are instead a product of the distribution of presences versus absences in the dataset, thus avoiding the biases introduced through MaxEnt's sampling process[43]. We used the recommended default values for $\gamma^+ = 1$ , $\gamma^- = 4$ and $m = 0.05$, the default values proposed in ref. [42].

$$\text{ASL} = -L_+ - L_- \begin{cases} L_+ = \bar{y} \cdot (1 - f(\hat{y}))^{\gamma+} \cdot log(f(\hat{y})) \\ L_- = (1 - \bar{y}) \cdot p(f(\hat{y}))^{\gamma-} \cdot log(1 - p(f(\hat{y}))) \end{cases}$$

$$\text{where } f(\hat{y_i}) = \frac{1}{1 + e^{-\hat{y_i}}} \text{ and where } p(\hat{y}) = max(p - m, 0)$$

The second bias-aware loss function we considered is a newly defined loss function we called scaled binary cross-entropy loss (scaled BCE) which simply weights the loss contribution of present versus absent classes by the number of present and absent classes, respectively. The effect is that the magnitude of the contribution to the loss of present labels versus absent labels is approximately equal; in other words the correctness of the predictions for present species matters as much as the correctness of the predictions for absent species when calculating the model's regret.

$$\text{Scaled BCE} = -L_+ - L_- \begin{cases} L_+ = \frac{\bar{y} \cdot log(f(\hat{y}))}{\sum_1^S \bar{y}_i} \\ L_- = \frac{(1-\bar{y}) \cdot log(1-f(\hat{y}))}{\sum_1^S 1-\bar{y}_i} \end{cases} \quad \text{where} f(\hat{y}_i) = \frac{1}{1+e^{-\hat{y}_i}}$$

For completeness, we also compare against ASL loss scaled using the same schema introduced in the scaled BCE loss.

$$\text{Scaled ASL} = -L_+ - L_- \begin{cases} L_+ = \frac{\bar{y} \cdot (1-f(\hat{y}))^{\gamma+} \cdot log(f(\hat{y}))}{\sum_1^S \bar{y}} \\ L_- = \frac{(1-\bar{y}) \cdot p(f(\hat{y}))^{\gamma-} \cdot log(1-p(f(\hat{y})))}{\sum_1^S 1-\bar{y}_i} \end{cases}$$

$$\text{where } f(\hat{y}_i) = \frac{1}{1+e^{-\hat{y}_i}} \text{ and where } p(\hat{y}) = max(p-m, 0)$$

We compared the performance of each of these losses on the uniform test set discussed in Section 1.3.1 using the image-only TResNet architecture. All losses were implemented in PyTorch and can be found in the code repository associated with the project. We found that our novel Scaled BCE loss had the best performance, highlighting the importance of caring about present versus absent classes equally.

##### 3.2.2 *TResNet* CNN architecture

Initially we compared a variety of CNN architectures, including VGGNet, ResNet and TResNet architectures[40,44,45] and early results showed the TResNet architecture to be superior (**Table S2**). This finding is further supported by the TResNet being one of the state-of-the-art architectures for multilabel image recognition tasks in other domains of computer science[45]. We slightly modified the default TResNet architecture to have four input channels in order to support the infrared band and three fully connected output layers instead of the standard single output layer, corresponding to the three taxonomic ranks being predicted: family, genus, and species. For learning rate, we tested a gradient of static learning rates from 0.01 to $10^{-5}$ and found a learning rate of 0.0001 to have the highest accuracy on the uniform test set. However, models trained with ASL loss did slightly better with a slightly higher learning rate, so we report accuracy using a learning rate of 0.0005 for these models.

##### 3.2.3 Generating a novel climate + remote sensing CNN

Finally, we were interested in testing a neural network architecture that could combine both remote sensing and climate sources to improve species modeling using CNNs. Previous work has shown that regional patterns in climate data can be interpreted by CNNs[46] but unfortunately climate rasters are too low resolution to be able to combine them with remote sensing images, and rather each 256 x 256 m will be assigned a single climate value (**Fig. S1**).

We then designed an architecture that combines the extracted features of both the image-only TResNet and the climate-only MLP during training to account for the difference in data scales. We refer to this as the "Joint TResNet" architecture (**Table S3**). The climate-only MLP takes in the raw bioclimatic information pointwise, while the aerial image data is still processed using 2D convolutions. We found that models trained with the climate and aerial image data tended to perform better than models trained with either aerial images or climate variables separately (**Tables S6, S7**).

##### 3.2.5 *Inception V3* Baseline

A recent machine learning competition (GeoLifeCLEF) that inspired our work, used NAIP imagery to predict a single species per image[41] using the *InceptionV3* architecture[47] (**Table S4**). Different from our work, this model employs the standard single-label CE loss, which cannot be used for joint species distribution modeling, and the competition was focused on ranking-based metrics such as top 30 accuracy or top 30 error.

Although the *InceptionV3* network was not designed for the multi-species prediction target of our work, we aimed to compare to it, as it is the CNN model most similar to *deepbiosphere* published so far. The difficulty in comparing *InceptionV3* with *deepbiosphere* is how the output values in *InceptionV3* are treated: The raw model logits can span from $-\infty$ to $+\infty$, which are then mapped to probabilities using a *softmax* probability transformation with the restriction that all probabilities across classes must sum to 1. This makes the probability of classes (species) dependent, as if a class is to rise in probability, another class must fall. This means that if a CNN trained with this loss has a high probability that one species is present, for it to also assign probability to another species it's confident is also present, it must decrease the assigned probability to the species already predicted as present. Naturally this introduces a problematic mutual exclusion when wanting to predict upwards of hundreds of species simultaneously. Further, converting these probabilities to binary thresholds for calculating accuracy metrics becomes tricky. To this note, we still train and compare against the *Inception*-based model used in this work, and report accuracies for probabilities calculated using the *softmax* transformation, meaning that the *Inception* model will naturally have very low predicted probabilities per-species and per-image, and thus all binary classification metrics are 0 for this model.

Another point to note is that in ref. [41] used a learning rate scheduler. We did not implement a learning rate scheduler in our training framework, so we only report accuracies for the model trained with a recommended constant learning rate of 0.01. We also use the auxiliary loss for model training, which previous work did not, which should improve accuracy by preventing vanishing gradients.

##### 3.2.6 Selecting the appropriate epoch for evaluation

Traditionally, the epoch to evaluate deep neural networks is decided based on the point at which the loss value on the test set is minimized. However, for many of the neural networks and test losses we considered, the test loss never minimizes, but many of the accuracy metrics do (**Fig. S5**). Therefore, to determine which epoch of training to evaluate the neural network-based SDMs, we calculated the average optimal epoch (defined as the epoch with maximized accuracy or minimized loss) for a subset of the reported accuracy metrics and loss. Specifically, we consider the accuracy of each deep neural network SDM on mAP, $AUC_{ROC_{spp}}$, $AUC_{PRC_{spp}}$, per-species precision, per-species recall, per-species F1 score, top 30 accuracy per-image, and the total loss. We use only a subset of accuracy metrics because some metrics take longer to calculate and are too inefficient to be recalculated each epoch during training, and also in order to introduce a nice balance of binary classification, discrimination, and ranking metrics. We also include the test set loss as this is the traditional way of determining model overfitting, although for certain loss functions in our domain, the minimization of the test loss doesn't correlate well with the epoch of best performance for most accuracy metrics. Since most metrics do not agree perfectly on which epoch is maximized / minimized, we use the average optimal epoch for all further downstream evaluation (vertical lines in **Fig. S5**). For *deepbiosphere*, the average optimal epoch was four, ten for *InceptionV3*, and forty five for the climate-only *MLP*. For comparing accuracy across loss functions, we evaluate all CNNs at the same epoch (epoch 20) to standardize comparison across accuracy metrics (**Table S6**).

#### 3.3 Climate-only Species Distribution Model baselines

Dozens, if not hundreds, of different SDM methods have been proposed over the decades, ranging from simple linear regression to neural network models. Two of the most popular methods, Maximum Entropy (MaxEnt) and random forest (RF), use significantly different methods of prediction and each have been cited thousands of times, warranting their choice as the two baseline SDMs for this study[48]. Concretely, we use the popular *dismo* package for species distribution modeling[49] and compare against two popular SDM approaches: Maxent and downsampled single stacked random forest using best practices lined out in ref. [50]. We chose these two models specifically from the dozens of approaches tried in ref. [51] as these two models had consistently the best performance across the hundreds of species in their dataset, besides ensembling approaches. We also attempted to compare against ensembling approaches and run the popular *biomod* ensembling algorithm, but the algorithm was too slow and memory-intensive for us to be able to run it on all species in our dataset.

We used WorldClim 2.0 bioclimatic variables[13] normalized to mean 0 standard deviation 1 as outlined above. Consistent with ref. [50], we removed all but one bioclim variable with a pearson correlation coefficient higher than 0.8, leaving ten variables in total for modeling including Mean Diurnal Range, Max Temperature of Warmest Month, Minimum Temperature of Coldest Month, Annual Precipitation, Precipitation of Wettest Month, Precipitation of Driest Month, Precipitation Seasonality, Precipitation of Wettest Quarter, Precipitation of Warmest Quarter, and Precipitation of Coldest Quarter.

For each species, we generated 50,000 background samples again consistent with ref. [50] by generating a circular overlay across all points in the training dataset where the radius of each overlay is the median distance between observations in the dataset. For the extrapolation experiments using the spatial cross-validation approach outlined in Section 1.3.2, we removed all background samples within the spatially withheld portion of the state. Finally, we used the same background points for both the random forest and Maxent models to ensure a proper comparison.

##### 3.3.1 Maximum Entropy Baseline

For Maxent, we use a stacked single SDM approach to get predictions across the 2,221 species by generating individual models for each species then aggregating the predictions *post-hoc* (see the intro to Section 3 for distinction). We use the Maxent implementation from the R package *dismo*[49] using the aforementioned background samples and all presences in a given train split of the main dataset, including the ‘nothreshold’ option consistent with ref. [50]. Although studies exist that run MaxEnt on the genus level, we nevertheless opted to only run MaxEnt to model species distribution, not genus or family. To build the joint prediction across all species for MaxEnt, we then aggregated the predicted score of each individual species from their corresponding projection. Maxent failed to run on 83 species which we exclude when calculating accuracy metrics. Fitting times and memory requirements for the uniform data split can be found in **Table S7**. Times are reported running MaxEnt serially per-species, utilizing one core of an Intel Xeon Gold 6240R CPU, but the actual processing was run in parallel using 24 Intel Xeon Gold 6240R CPUs, leading to a total wall clock time of 5.69 hours. We report serial wall clock time for MaxEnt since it cannot fit species simultaneously like the CNN-based approaches and thus running MaxEnt across so many species on a small compute setup (i.e.: 10 GiB memory, 4 CPUs) would take a considerable amount of time compared to the CNN-based approach which is very lightweight and can run on a small, inexpensive 8 GiB NVIDIA M60 GPU with 10 GiB memory and 4 CPUs in under 20 hours. Therefore, we felt the fairest time comparison would be to report wall clock time in an expected low-resource environment where one does not have access to large and expensive compute to widely parallelize the SDM-fitting process. For the extrapolation experiments model fitting on all ten splits of the dataset varied widely due to differences in cluster-specific latency, ranging from 139 to 50 hours to run and ranging from 90 seconds per-species to almost four minutes per species. Due to this cluster-specific variance, these fitting times are thus not reported.

##### 3.3.2 Random Forest Baseline

For the Random Forest baseline, we also take a stacked single SDM approach and use *dismo*[49]. For each species, we fit a Random Forest with 1,000 trees using equal bootstrapping of positive and negative samples with replacement as outlined in ref. [50]. All other options were the *dismo* default settings. 66 species did not properly fit for Random Forest, and those were excluded from subsequent analyses. Fitting times and memory requirements for the uniform data split can be found in **Table S7**. For the extrapolation experiments model fitting on all ten splits of the dataset varied widely due to differences in cluster-specific latency, ranging from 139 to 6 hours to run and ranging from 12 seconds per-species to almost four minutes per species. Due to this cluster-specific variance, these fitting times are thus not reported.

##### 3.3.3 Climate-only Multilayer Perceptron baseline

To compare the difference in remote sensing data versus standard bioclimatic data for species distribution modeling, we also considered how well a standard fully-connected multilayer perceptron (MLP) trained using bioclimatic variables would perform as an SDM. MLPs trained on environmental data are a standard choice for SDMs, although Maxent and random forest have been more popular in recent years[51]. To do so, we implemented a four layer, fully-connected MLP with BatchNorm (**Table S5**). Specifically, the model's architecture was inspired by ref. [52] and consisted of two fully-connected layers with 1,000 neurons each, followed by a dropout layer with a 0.25 dropout rate, then by two layers with 2,000 neurons each, before predicting species, genus, and family. Batch size, total number of epochs, memory usage and training time can be found in **Table S5**. It should be noted that this MLP is *not* convolutional and learns from the raw value of environmental variables, rather than two-dimensional patterns of remote sensing data, like the CNN models do. The climate-only MLP was also trained using the Scaled BCE loss function discussed in Section 3.2.1.

### 4. Individual species case studies

While the ability to successfully predict the presence of thousands of species at once is in itself an impressive feat, also of importance is *how* such an SDM can be useful for a variety of downstream ecological tasks. By modeling each individual species of a community, we can begin to detect not just individual species, but ideally patterns of the entire community as well. To begin to explore the power of a multi-species modeling approach, here we present two case studies of well-known vegetation communities and their species and demonstrate how *deepbiosphere* can detect expected species-level range dynamics across a variety of ecoregions.

#### 4.1 Validating *deepbiosphere's* predictions for individual species

Since *deepbiosphere* is a joint SDM, it can be used to detect and model individual species at a time, along with the broader community. However, how to validate the accuracy of the model for a given species' range is difficult without on the ground, gold-label observations. Barring onsite validation, we opted to instead compare how well *deepbiosphere* could classify species presence for two large charismatic tree species—Redwoods and Valley Oak—compared to humans attempting the same task. Specifically, we compared *deepbiosphere's* predictions to humans labeling the exact same aerial imagery that *deepbiosphere* predicted from. We specifically chose these two species as they are observable directly from the NAIP imagery so that human annotators could distinguish their specific canopies and the human annotations could be reasonably assumed to represent some proxy of ground-truth presence and absence. Even though on-the-ground observations to corroborate the model's predictions don't exist for either species, fine-scaled vegetation maps for both species' ranges do exist and can be used as a proxy for both species' range. Using these vegetation maps, we can cross-validate the human annotations to *deepbiosphere's* predictions with a second independent data source.

##### 4.1.1 Generating high-resolution species range maps with CNN SDMs

*Deepbiosphere*, like all classification-based CNN models, takes in images of ideally a set dimensionality (for our work, 256 x 256 pixels) and makes a single prediction for this image (in our formulation, a single prediction consists of a predicted presence of 2,221 plant species). In other words, *deepbiosphere* can produce a prediction of all 2,221 species for any arbitrary 256 x 256-pixel image. In order to get a map of predictions at a set ground-level resolution using this model, one simply has to slide this so-called local receptive field down every K pixels to generate a map of K-pixel resolution, a technique similar to the striding operation in the convolutional layers of a CNN. Then, depending on the resolution of the imagery used to make the predictions, one can determine the ground-level resolution of the predictions themselves. For example, to generate a 50 m ground-level resolution map from 1m NAIP imagery, a stride of 50 should be employed. Alternately, to generate a 30 m ground-level resolution map from 60 cm NAIP imagery, a stride of 50 will also suffice. A visual explanation of the striding procedure can be found in **Fig. S8**.

##### 4.1.2 Human annotation protocols

In order to generate ground-truth human annotations at a similar resolution to *deepbiosphere's*, a user study was implemented in Google Sheets where human annotators classified the same aerial imagery as *deepbiosphere* by percent cover. Annotators were not domain experts and the only training received *a priori* were three already classified example images. (**Figs. S9, S14C**) These already classified images were taken from the same 2012 NAIP acquisition as what *deepbiosphere* was trained with to ensure standard data quality. These examples were deliberately chosen to be outside of the case study area but within the species' core ranges and were annotated on a scale of 0% cover to 100% cover, with five possible categories. Annotators were given the NAIP imagery partitioned into 256 × 256m blocks, which were labeled A-Z and 1-30 to correspond with the appropriate cell in Google Sheets. In total, three annotators annotated the redwoods case study and two annotators annotated the oaks case study.

##### 4.1.3 Redwoods case study

For the redwoods case study, both *deepbiosphere* and *MaxEnt* were evaluated with models not trained with any examples from the 10th latitudinal cross-validation block, meaning neither model was fitted using any observation within at least 10 km of the case study area. For the *Inception* baseline, the model trained with the uniform dataset split was used for evaluation, meaning that the model did see examples from the case study area during training. To validate the predicted presence of redwoods across vegetation types and models, the National Park Service's 2017 vegetation mapping and classification project was used, specifically the alliance-level mapped to the 30 vegetation classes used for accuracy assessment (see Section 6.2.1 for details)[53]. The co-occurring species used for the understory analysis were chosen based on constancy values reported in ref. [53] for Manual of California (MoC) vegetation associations that crosswalked to either the *Sequoia sempervirens*-(Other) YG Mixed Forest class or the *Sequoia sempervirens* Mature Forest class. Species were considered to be associated if they had a constancy value of >=60% for each MoC vegetation association that crosswalked to the given class, and <=40% constancy value for the crosswalked MoC associations for the other class.

For comparing the predictions of understory herbaceous species in the redwoods case study, the difference in *O. oregana* versus *I. douglasiana* predictions was calculated by subtracting the predicted probability of *I. douglasiana* presence from the predicted probability of *O. oregana*, per-pixel. For pixel-wise probability comparisons, a non-strided prediction map for each SDM (for *deepbiosphere*, this corresponds to generating a map at 256m resolution) was generated to minimize spatial autocorrelation. For the redwoods case study, the vegetation type of each pixel was labeled by determining the vegetation class from the NPS map with the largest area overlapping the pixel. Pixels were considered "inside" a vegetation type if they were labeled with the corresponding vegetation class(es) and "outside" otherwise.

##### 4.1.4 Oaks case study

For the oaks case study, the USDA Forest Service's South Coast existing vegetation map[54] (REGIONAL_DOMINANCE_TYPE key) was used to validate species predictions, crosswalking vegetation types to species using the CALVEG class descriptions from zone 7[55]. A species was considered present within a given regional dominance type if said species' name was mentioned in the corresponding CALVEG vegetation description. The final species to CALVEG mappings are as follows: *Ceanothus cuneatus*: CC, CQ, EX; *Quercus lobata*: QL; *Bromus diandrus*: HG; *Quercus berberidifolia*: CQ; *Arctostaphylos glandulosa*: CQ, SD, *Adenostoma fasciculatum*: QA, CC, CQ, SS, EX. Other CALVEG classes were present in the study area, but either were associated with developed and agricultural land use or were too small in area to map to a sufficient number of pixels. For comparing predicted presence inside versus outside CALVEG zones, *deepbiosphere* predictions made at 256m resolution were used to minimize spatial autocorrelation. For a given species, pixels were marked as "inside" if a given pixel intersected at least one of the associated CALVEG classes for that species, and was marked as 'outside' otherwise.

### 5. Mapping spatiotemporal changes with *deepbiosphere*

On top of modeling individual species, *deepbiosphere's* joint species predictions can also be evaluated in aggregate to explore more macroecological trends in community composition, such as spatial and temporal turnover. Historically, these processes have been difficult to observe directly without detailed species checklist data, limiting the resolution at which these metrics can be generated[56,57]. However, combined SDMs for thousands of species have been proposed previously to measure certain important ecological phenomena, such as imperiled species index[29]. Here, we showcase that *deepbiosphere* could potentially be used for similar purposes, especially for detecting community turnover in space and time at high-resolution.

#### 5.1 Detecting spatial community turnover

In order to interpret *deepbiosphere's* predictions of species presence as community turnover, a novel edge detection algorithm inspired by common edge detection filters from computer vision was used (see **Fig. S1**9 for visual walk-through). Specifically, the averaged one-neighbor euclidean norm is calculated per-pixel from a map of all 2,221 species predictions (See Section 4.1.1 for map generation details) to then generate a map of averaged similarity to neighbor pixels. This algorithm essentially measures the average distance of the predictions of a given pixel to all its nearest neighbor pixels, summarizing how similar or different a given pixel's predicted species list is from nearby areas. Another interpretation of this statistic is as a measure of the average rate of change of species composition within a local area.

The Euclidean norm was chosen as the statistic to measure between pixels as it is a metric that encodes both magnitude and direction. Therefore, it will capture both changes in a given species' presence across pixels through the directional component (e.g: a species highly predicted in one pixel has much lower predicted probability in the other pixel) and capture changes in the raw number of species predicted through the magnitude component (e.g.: many species are predicted with high probability in one pixel, but then very few are predicted with high probability in the other pixel). Generalizing, for all non-edge pixels in the given extent of *deepbiosphere's* presence predictions, we take the Euclidean norm from the central pixel to its neighboring pixels, then take the average across its eight neighbors to generate the final spatial community change value. Below, $\hat{y}_c$ is the predicted probability vectors for all 2,221 species at the given pixel of interest and $\hat{y}_{1N}$ are the predicted probability vectors at all eight one-neighbor pixels (see **Fig. S1**9 for visual walk-through).

$$\text{average one-neighbor Euclidean norm} = \frac{\sum_{i=1}^{8} \|\hat{y}_c - \hat{y}_{1i}\|}{8}$$

This averaged one-neighbor Euclidean norm metric we refer to as the *spatial community change*. Directly validating spatial community change is exceedingly difficult, as such data rarely exists at scale. As a proxy, the number of unique alliance-level vegetation classes in a given pixel is used to approximate how rapidly habitat transitions are occurring in a given location. The idea is that more vegetation classes intersecting in the pixel means that the area is likely an ecotone, and thus should have a higher spatial turnover of species. To validate this approach, a case study from north Marin County of California was used as this region has a highly varied landscape, plus it sits at the transition point between two ecoregions of California, and is thus a known site of ecosystem transition. Finally, the area's vegetation was recently classified down to the association level in 2021[58] (**Fig. S17C**). The number of intersecting alliance-level vegetation classes were counted per pixel using Geopandas' "intersects" function[59] and the spatial community change metric was correlated to the number of vegetation classes using the modified *t test* from SpatialPack, again using the centroid of each pixel as the coordinates per-sample[60]. To confirm that predicting changes in community composition is not as simple as simply predicting the change in greeness or infrared absorption, the average Euclidean distance between the channel values of NAIP imagery was used as a baseline and correlated to the number of intersecting vegetation classes as described above (**Fig. S17D**). To further confirm that predicting changes in community composition is also not as simple as simply recapitulating the number of species in a given area, the number of unique species observed in a given pixel using the biodiversity dataset was also correlated with the spatial community turnover from *deepbiosphere*$_7$ (**Fig. S17F**).

### 5.2 Detecting temporal community turnover

One major benefit to using remote sensing data to train SDMs as compared to climatological datasets like WorldClim is that remote sensing data often has a shorter time period between acquisitions[61]. For example, WorldClim provides bioclimatic variables for one fixed time point for the current historical period, along with future averages in 20 year intervals[13]. Capturing temporal ecological changes due to natural events such as drought, fire, or flooding is simply not possible at such a coarse temporal resolution. Meanwhile, NAIP data is acquired every two years, while a full Landsat acquisition of the earth happens roughly every 16 days. With this fine-grained temporal resolution, ecological changes year-to-year, season-to-season, and even day-to-day are potentially detectable.

Fire is a natural part of many of the ecosystems of California, with a series of unique plant communities that succeed one another following a fire event. However, almost a century of fire suppression has disrupted the cyclic nature of these successional changes in community composition in ecosystems adapted for fire, altering these different steps in the succession and even leading to more intense fires in California[62,63]. To this point, the Rim Fire that occurred in the Sierra mountain foothills was chosen as the case study. At the time of the blaze, the Rim Fire was California's second-largest fire on record. More specifically, the blaze occurred in the Hetch Hetchy Valley in the western Sierra mountains, and occurred in the fall of 2013, giving an entire growing season between the fire and the acquisition of the next round of NAIP imagery in summer of 2014. In order to detect temporal change, a metric similar to the spatial community change discussed in Section 5.1 was employed. Specifically, the Euclidean distance between *deepbiosphere's* predicted species probabilities made from NAIP imagery acquired in 2012 and 2014 was used to approximate the magnitude of temporal community change.

$$\text{temporal Euclidean distance} = ||\hat{y}_{2012} - \hat{y}_{2014}||$$

This temporal Euclidean distance metric we refer to as the *temporal community change*. We used the community change metric outlined above, with an example of the underlying data for this algorithm found in **Fig. S21**.

As with spatial community change, directly detecting temporal community change is difficult. However, scientists did collect hyperspectral data within the bounds of the fire at high resolution, enabling the estimation of the differenced normalized burn ratio (dNBR), a popular metric of fire severity[64]. Normalized burn ratio (NBR) is an empirical measurement of burn severity calculated by taking the difference between a near infrared wavelength—which captures photosynthesis intensity—and a shortwave infrared wavelength—which captures heat absorption from char—which is typically measured using hyperspectral spectrometers such as AVIRIS or MASTER sensors[65]. To calculate dNBR, the change in NBR from data acquired before the fire to data acquired after is calculated. For this analysis, MASTER sensor data was chosen as it had more coverage over the fire than the AVIRIS sensor data, covering roughly half of the fire's extent. Specifically, dNBR calculated using NBR acquired in June of 2014 was used for validation, as that was the closest time point to the acquisition of the corresponding NAIP imagery. In order to reach the same resolution as the empirical burn severity data, we used a 35 pixel stride as outlined in Section 4.1.1.

Deepbiosphere's predicted temporal community change metric was correlated to dNBR using Pearson's correlation corrected for spatial autocorrelation using the Dutilleul correction from the SpatialPack R package[60,66]. Before correlation, both dNBR and *deepbiosphere's* temporal community change predictions were upsampled to 256 m resolution as the spatial correction calculation is very computationally intensive. The Pearson's *r* does not change substantially between correlations calculated at 35m versus 256m resolution, thus only the 256 m results are reported.

As with the spatial community change, the Euclidean distance between NAIP imagery acquired in 2012 and 2014 was calculated to confirm that detecting fire severity is not as simple as recapitulating the difference in the infrared and green bands of the NAIP imagery used to train *deepbiosphere*. NAIP imagery was upscaled to 35 m resolution, normalized, and mean-centered in the same way that imagery is prepared during *deepbiosphere* model training. The correlation between the distance in NAIP pixels and dNBR was also generated at 35 m and 256 m resolution and as with the temporal community change, the Pearson's R does not change substantially between correlations calculated at 35 m versus 256 m resolution.

Finally, we also compared the change in predicted probabilities for two charismatic species with different fire response strategies—Aspen and Golden Yarrow—to show how *deepbiosphere* appears to capture expected changes in abundance before versus after the fire (**Figs. S22, S23**). To do so, the signed difference in probability for each species' predicted presence before versus after the fire was compared.

### 6. Using *deepbiosphere* for downstream spatial mapping tasks

Many important biodiversity conservation tasks and broader mapping efforts rarely possess the volume of data necessary to train a deep neural network like *deepbiosphere* from scratch. Since the species present in this biodiversity dataset span many ecosystems and plant species, both native and introduced, the resulting predictions of species presence may be useful as a predictor variable for a variety of these data-sparse tasks, enabling the seamless generation of these maps from the few data that do exist.

#### 6.1 Using *deepbiosphere* as a feature extractor for downstream maps

To use *deepbiosphere* predictions for a wide variety of downstream applications, we employ *deepbiosphere* as a feature extractor by sequentially feeding in remote sensing imagery and extracting the 2,048 outputs from the final layer of the CNN (Linear: 2-23 in **Table S3**, rightmost gray bar in **Fig.**

**1B**) to use as features for a downstream classification. In other words, for a given downstream task such as crop type mapping, for each example in the downstream training dataset, the latitude and longitude of the example is taken and a 256 x 256 m of NAIP imagery is cropped, centered at the location of the example (see **Fig. S2, S28** for visual examples). Then, that image is fed into *deepbiosphere* and the outputs of the last hidden layer are cached for each example. Finally, a downstream classifier such as random forest or logistic regression is fitted using the downstream training set labels as output and the *deepbiosphere* outputs as input. For accuracy analysis, the same process is repeated but for held out test examples. For generating entire maps, a given section of NAIP imagery is taken and for each strided image in the area, the outputs of the last layer of *deepbiosphere* from that image are taken and classified using the downstream classifier.

#### 6.2 Generating an alliance-level vegetation map of Redwoods National & State Parks

To use *deepbiosphere's* features to generate a vegetation map for Redwoods National and State Parks (RNSP), field-based vegetation type measurements from the 2017 NPS mapping effort were used. Field plots with no supplied map class and plots with duplicate latitude and longitudinal values were removed, leaving a total of 1,198 field-validated vegetation classes for training and testing. To properly compare to the accuracy assessment performed by RNPS, we first cross-walked the provided association-level vegetation type (see section 2 of ref. [53] for details on class type designation from the plot-level relevé forms) to the generalized alliance level. For field plots mapped to the Manual of California Vegetation associations, we used the cross-walk provided in Appendix B. of ref. [53] to also map those plots to the generalized alliance level. We further grouped the alliance-level classes into the 30 vegetation classes used in the RNPS accuracy assessment (see 6.2.1 of ref. [53] for details on the map classes; see sections 5 and 6 of ref. [53] for justification on classes that were removed) and filtered field plots to only include those mapped to the 30 accuracy assessment classes.

For each field plot, we then cropped a 256 x 256 pixel image from NAIP imagery, centered at the field plot in question (see **Fig. S2** for an example). The images were then fed to $deepbiosphere_{10}$ and the 2,048 features from the last layer were extracted. To decrease the downstream classifier's complexity, we only retained features with a non-zero value for at least one field plot, leaving a total of 955 features for fitting. Of the 1,198 field plots provided by the RNSP, 423 were marked as training plots, 489 as testing plots, and 286 as additional plots not used for fitting the final RNSP vegetation map (**Fig. S24B**). The RNSP vegetation map was built with 462 field plots (the additional plots came from locations overlapping the 286 additional plots which were removed as duplicates in our analysis) while we utilize both the 423 training plots and 286 additional plots. To minimize overfitting and improve accuracy and ensure that there were more training points than training features, we augmented the 709 field plots with an additional 300 points bootstrapped from the accuracy assessment test set across ten cross-validation trials, leaving a total of 1,009 field plots for fitting downstream vegetation classifier. We compare across three standard downstream classifiers: random forest, multilayer perceptron and logistic regression using the 189 remaining accuracy assessment test plots per-fold. For each downstream classifier, we use the standard scikit-learn version 1.1.1 implementation with default parameters except for logistic regression, where we use the 'liblinear' solver[25]. As a trivial baseline, we also generate random Gaussian noise and use it to fit the same downstream classifiers from the field data. To generate a full vegetation map of RNSP, we selected the downstream random forest classifier with the highest test set accuracy (63.49% single-label accuracy), then extracted final layer predictions from *deepbiosphere* using the same striding approach as before to achieve a per-pixel resolution of

50m. The features for each pixel were then classified using the downstream random forest classifier to generate a full alliance-level vegetation map of the entire region.

#### 6.3 Generating a crop type map of San Joaquin Valley

For cropland type mapping, we decided to compare against a previous state-of-the-art unsupervised deep learning approach called Tile2Vec[67]. Utilizing the same San Joaquin Valley study area as the original Tile2Vec paper (**Fig. S28B**), we partitioned the 2016 USDA cropland data layer map for the San Joaquin Valley[68] into the same train-test-validation split used by ref. [67] (**Fig. S28D**) and randomly sampled 1,000 geographic locations within the training blocks, ensuring that locations were farther than 128 m away from the boundary edge to minimize examples bleeding over into adjacent blocks. We also removed the top row of training blocks to ensure that no location had been previously seen by *deepbiosphere*$_5$ during training (**Fig. S28A**, maroon band). For each of these 1,000 locations, a 256 x 256 pixel image was cropped from 60 cm resolution 2016 NAIP imagery (**Fig. S28E**), which is of much higher resolution and a different year than the 1m resolution 2012 NAIP imagery used to train *deepbiosphere*$_5$ (**Fig. S28F**). A 50 x 50 pixel image was also extracted for the Tile2Vec network, as *TileNet* was initially trained with much smaller images than *deepbiosphere* (**Fig. S28E**, white box). For both the 256-pixel and 50-pixel image, the cropland class with the largest area in the image was extracted from the 2016 USDA cropland data layer for each image, respectively (**Fig. S28G**). Since the size of the prediction window is very different between *deepbiosphere* and *TileNet*, occasionally the cropland labels will differ for the two networks (i.e.: in the example from **Fig. S28G**, the class with the largest area is almonds for *deepbiosphere* while the class with the largest area is developed / open space for *TileNet*). The same analysis was also performed with 10,000 training examples, and those results are reported in **Table S13**.

We downloaded the *TileNet* architecture and trained weights from the open Github repository (https://github.com/ermongroup/tile2vec) and then fed the cropped NAIP images to their respective deep neural network. The features from the last layer were then extracted (2,048 features for *deepbiosphere* and 512 features for *TileNet*). Three types of downstream classifiers—random forest, multilayer perceptron and logistic regression—were fitted using the 1,000 features and cropland classes, respectively. For each downstream classifier, we use the standard scikit-learn version 1.1.1 implementation with default parameters[25]. We also fit the same suite of classifiers using 1,000 subsetted training examples to fully compare to ref. [67]. To compare the network's accuracies, we extracted features in the same manner as above for 1,000 random locations sampled from test blocks, then compared accuracies across ten different instantiations of the same downstream classifier. To generate a full cropland map of the region, we again used the highest accuracy random forest classifier from across the ten trials (66.9% single-label accuracy) and classified the final layer predictions extracted from *deepbiosphere* using the 2016 NAIP imagery with a 50-pixel stride, generating 30m resolution cropland predictions.

SUPPLEMENTAL FIGURES

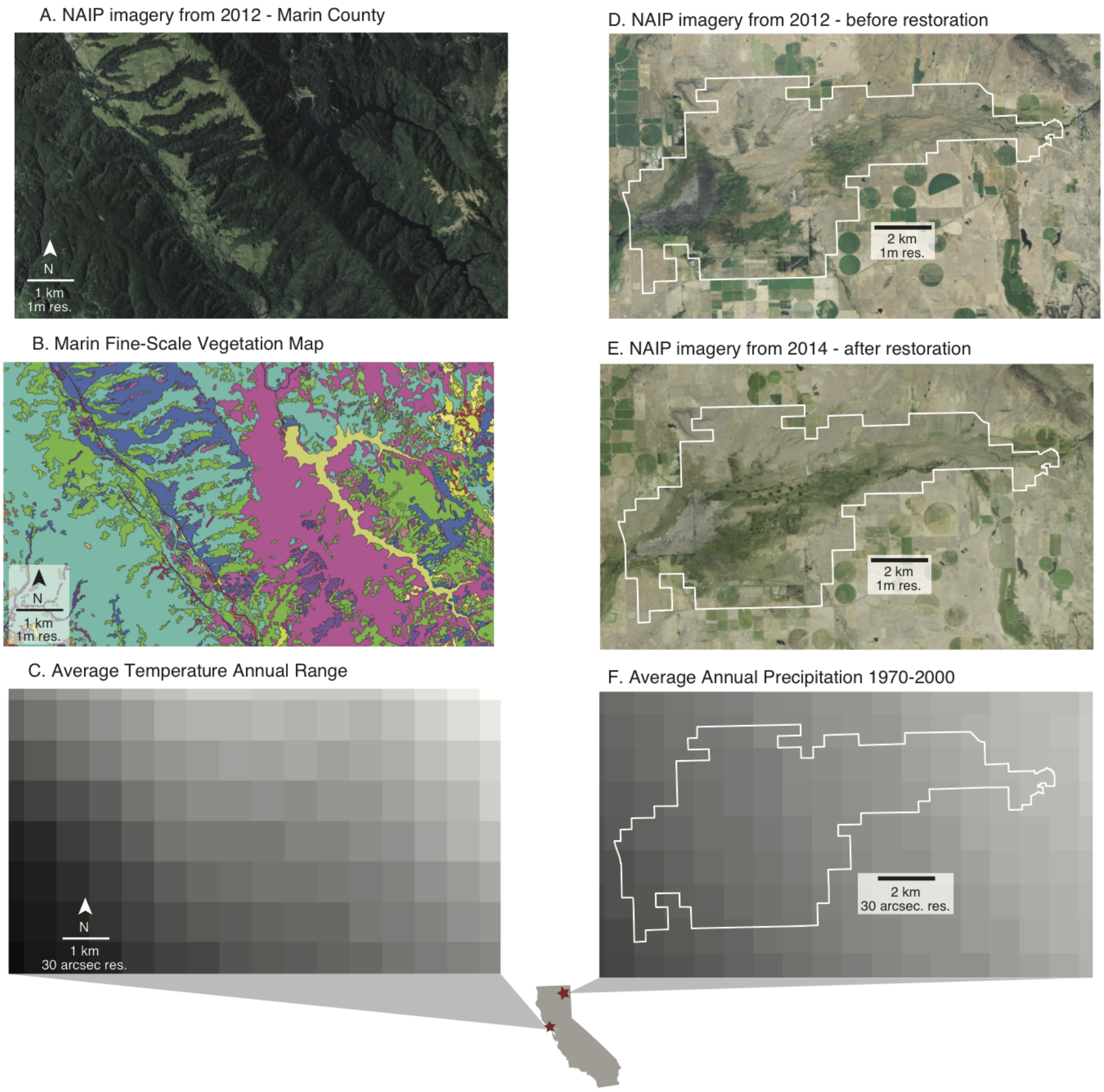


**Fig. S1 | Comparison of the spatial and temporal resolution of remote sensing imagery vs. bioclimatic data.**
**(A)** National Aerial Imagery Program (NAIP) imagery data acquired in 2012 for northern Marin County in northwest California. From this imagery, different ecological features of this forest-grassland-lake transition area are clearly visible, including the light green of the invasive annual grasslands in the middle interspersed with patches of coast live oak. **(B)** Fine-scale vegetation map of the example area from ref [58]. According to this vegetation map, this region contains almost 50 unique habitats and land use categories, fragmented into small units that are clearly visually distinguishable from the remote sensing imagery in (A). **(C)** The 7th WorldClim 2.0 bioclim variable—temperature annual range[13]—has a low spatial resolution and does not capture the local habitat changes in (B) that are clearly visible from remote sensing imagery in (A). It should be noted that the projection for the BioClim variable is a geographic coordinate system, not a projected coordinate system and does not preserve distance; thus each represented pixel is not fully 1x1 km and is rather 30 arc-seconds. **(D)** National Aerial Imagery Program (NAIP) imagery data acquired in 2012 at a degraded ephemeral freshwater wetland at Ash Creek Wildlife Area in northern California (park bounds outlined in white). The degradation of the wetlands is clearly visible as the large light brown grassland intrusion cutting through the middle of the much greener and darker-colored wetland (center of image) **(E)** NAIP imagery acquired in 2014, after a restoration project was undertaken to restore the degraded wetland through a series of pond and plug procedures (visible as small dark dots in the center of image). The restored

wetlands are clearly visible as a now-continuous band of green vegetation traversing the majority of the wildlife area. **(F)** 30 arc-second version of the WorldClim 2.0 12th bioclimatic variable for Ash Creek Wildlife Area[13]. The WorldClim bioclimatic variables are averaged across a three decade timespan (1970-2000) and thus fail to meaningfully capture this rapid temporal ecosystem change.

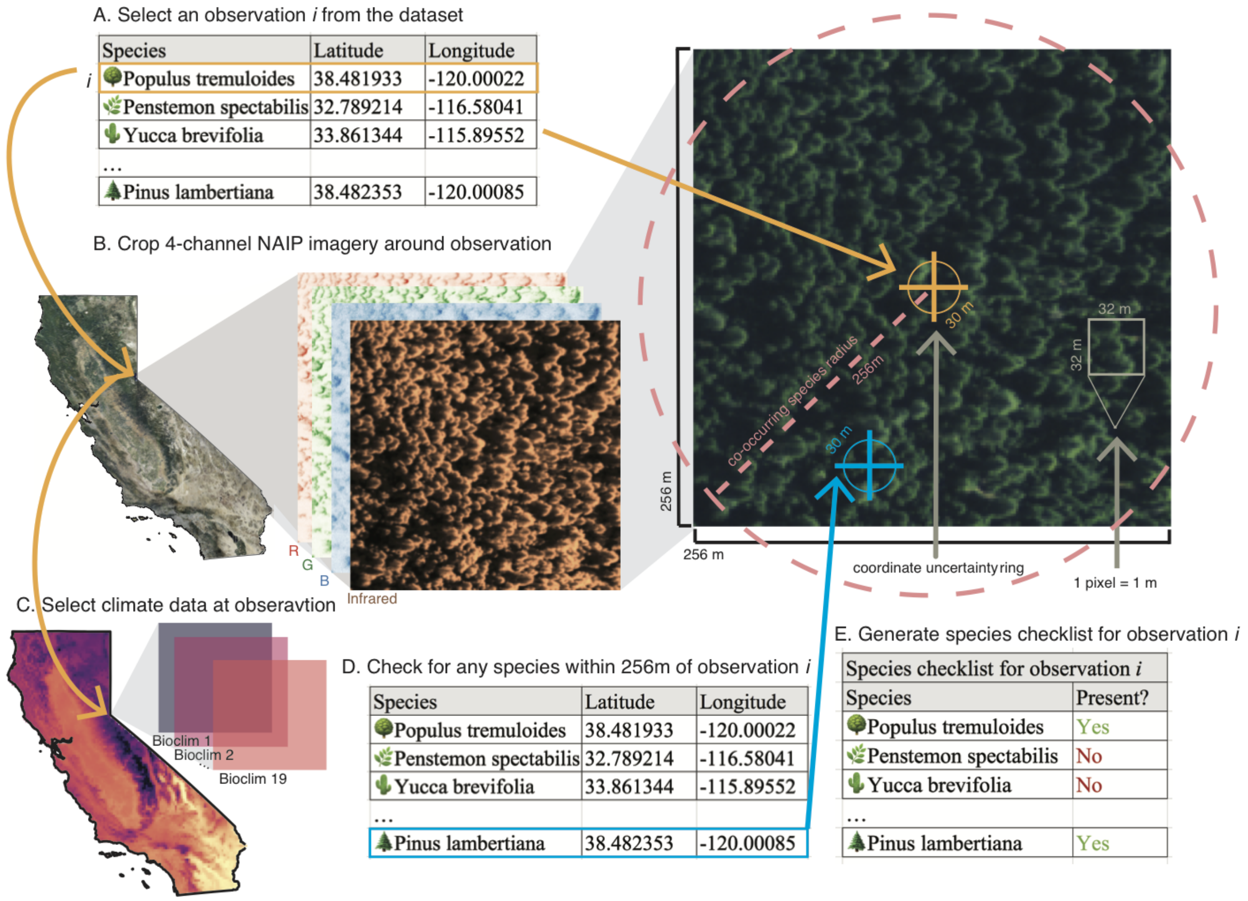


**Fig. S2 | Illustration of building the biodiversity dataset**
**(A)** First, a species observation is selected from the dataset. This process occurs for every unique observation in the dataset **(B)** For this selected observation, a 256 × 256 pixel image (pixel resolution is 1 m) is generated from the NAIP imagery, centered at the geographic location of the observation (orange cross). Observation locations may have upwards of a 30 m radius of geographic uncertainty (orange circle) but still fall well within the image. **(C)** The list of bioclim variable values for that location are generated. **(D)** Next, a list of overlapping species is generated by selecting other observations (blue cross) from the dataset whose coordinates fall within the 256 m radius of the original observation (pink circle) **(E)** The final data products consist of the 4-band remote sensing image (B), the bioclim variables (C) and the partial species checklist (E).

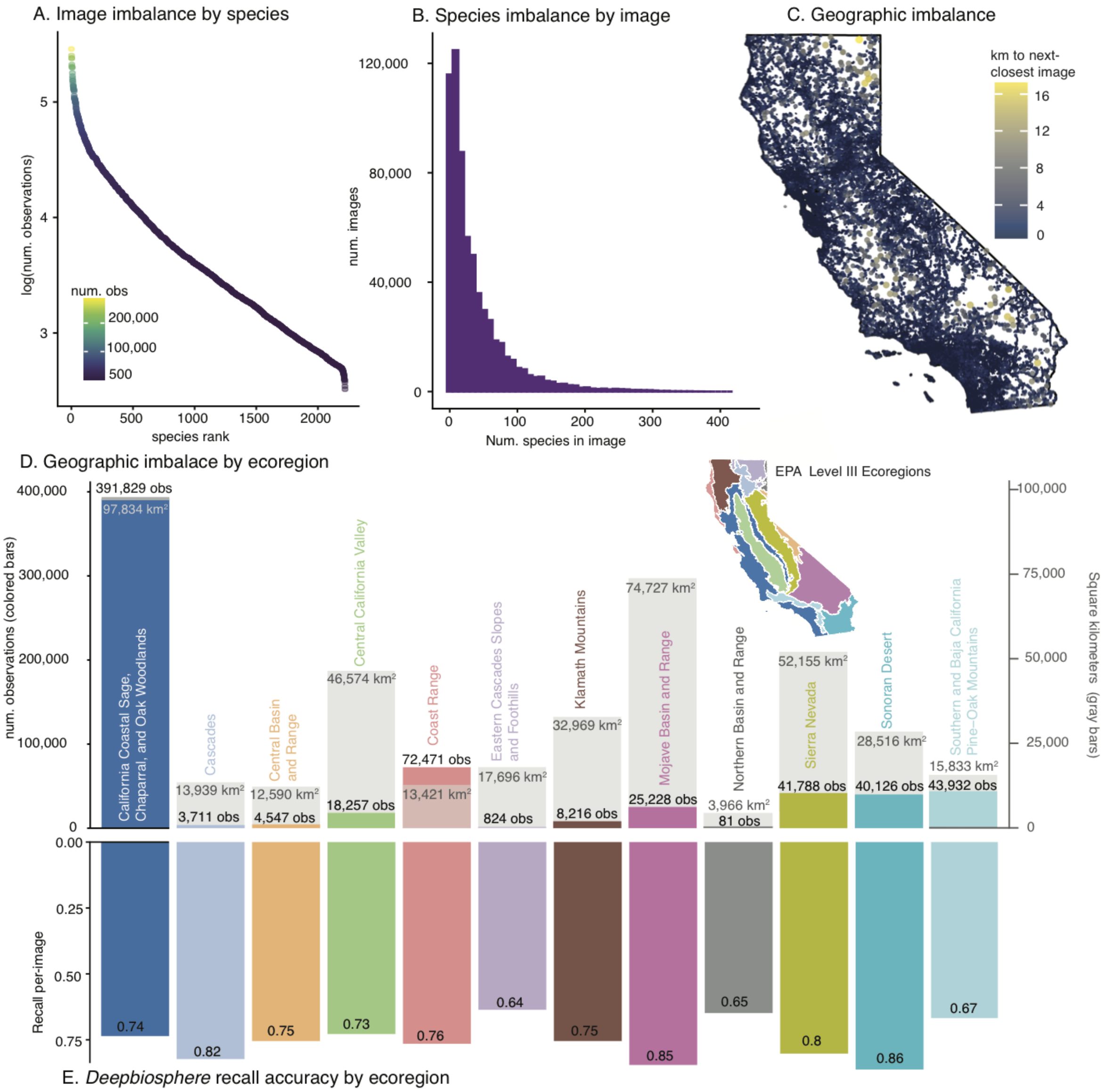


**Fig. S3 | Biases present in the dataset and their effects on accuracy**
**(A)** Many species have few images in the dataset, while a few species have a disproportionate number of observations. This extreme imbalance between species class frequencies can make it difficult for classic machine learning methods that rely on an assumption of an equal number of examples per-class to learn a good representation. **(B)** This histogram highlights the significant imbalance of number of species labeled per-image in the dataset. Most observations have few overlapping species in the observation—likely not fully describing all species actually present in a given site—while only a few observations contain many species and present a more accurate checklist of species presence in a given area. **(C)** Plot of distance to nearest non-overlapping observation across California. Color and size represent the distance in kilometers to the next-closest non-overlapping image in the dataset. As can clearly be seen, observations tend to cluster and distances are not distributed evenly. **(D)** Comparison of number of species occurrences in a given Level III EPA ecoregion versus the area of that ecoregion. Citizen science observations like those used in the dataset tend to cluster around population areas and natural regions where observers can reach. This leads to oversampling in some ecoregions, especially the California Coastal Sage, Chaparral and Oak Woodlands and Coast Range, and significant undersampling in other regions, especially the Cascades and Northern Basin and Range habitats. Colored bars represent the number of observations in the dataset from that region and the grey bars represent the area in square kilometers of the specific ecoregion. Inset is a map of the level III ecoregions of California. **(E)** Average per-image recall accuracy of *deepbiosphere* by ecoregion using uniformly sampled held-out observations (**Fig. S4A**).

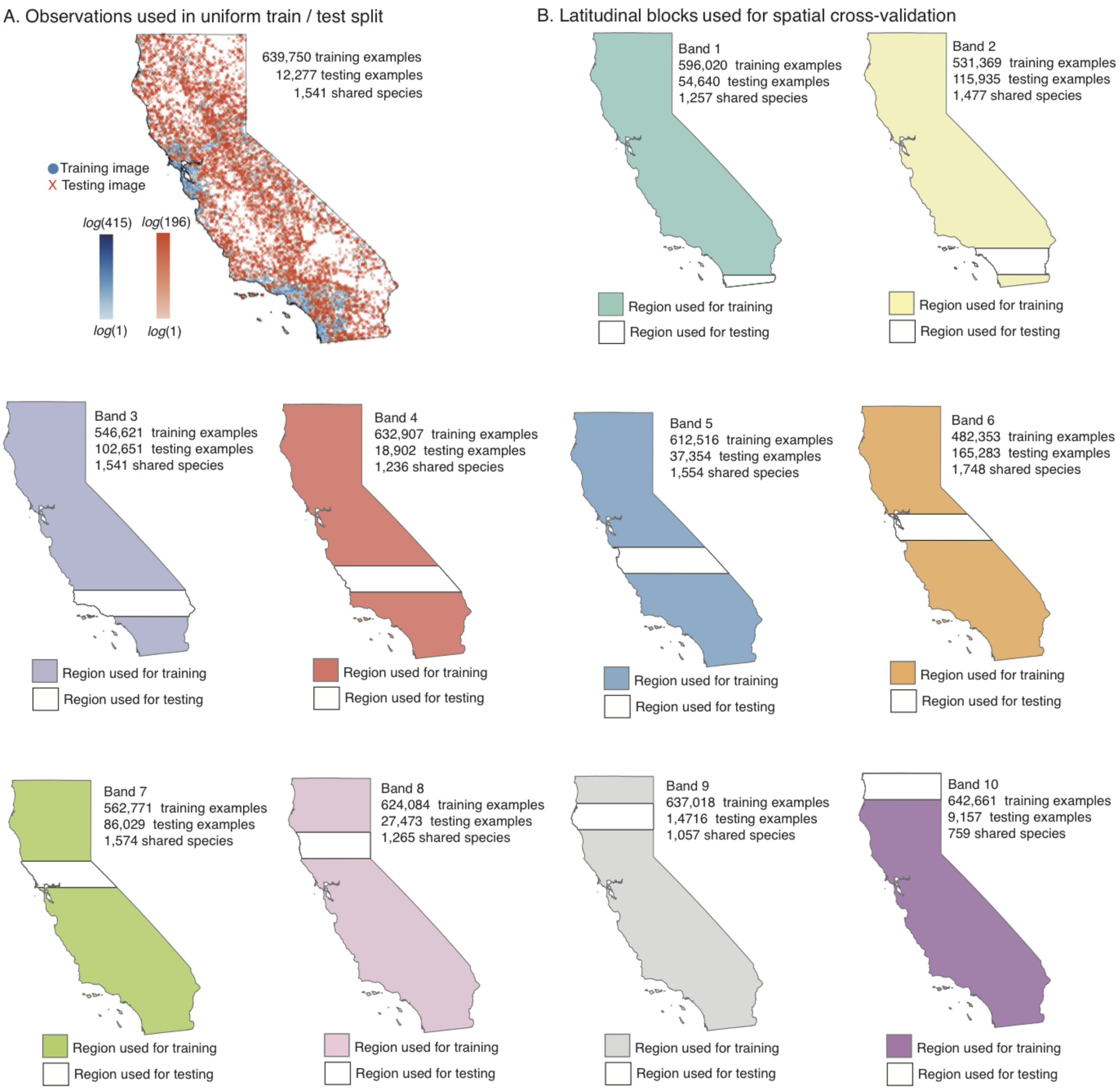


**Fig. S4 | Partitioning the dataset for cross-validation**

**(A)** Location of observations used for uniform, interpolated holdout dataset. Observations were selected for the test set if they were at least 1.2 km away from any observation in the training set to ensure that there was no train/test leakage for bioclim-trained models. Color scales represent the number of unique species present in each observation, log-scaled. **(B)** Location of test blocks for spatial cross-validation. We employed a 10-block spatial holdout procedure to test models' extrapolation ability. Areas that were included in the training set are in color and areas used for the test set are in white. Any images within 1.3km of the start of the test region were removed from the training set in order to prevent potential overlap between bioclimatic variables in the train and test splits (grey boundary). The number of images within each split is also annotated, along with the number of unique species present in both splits.

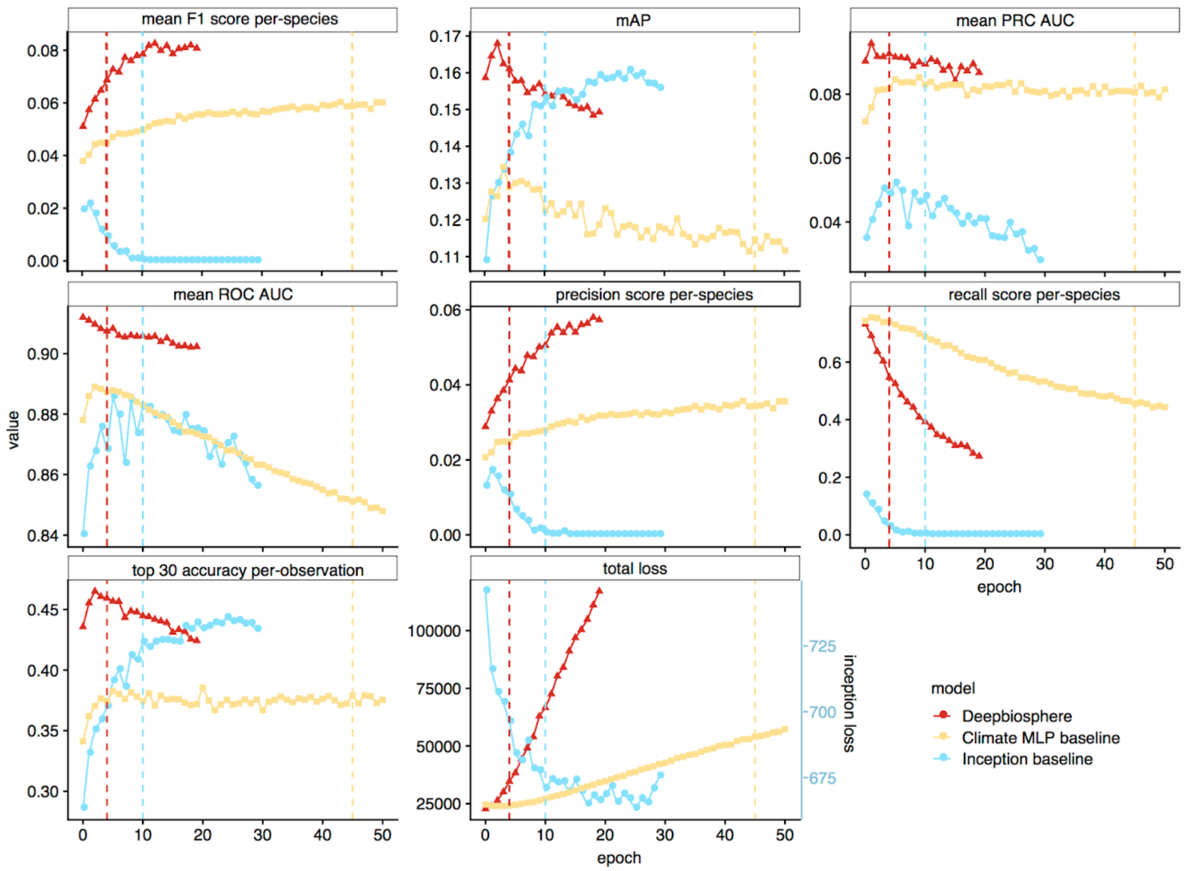


**Fig. S5 | Per-epoch accuracy and loss used to calculate the average optimal epoch for downstream evaluation**
In order to decide which epoch to evaluate the deep learning models, we calculated the max accuracy epoch for seven representative accuracy metrics, plus the total loss on the uniform split of the dataset (**Fig. S4A**), and used the average max accuracy epoch as the epoch of evaluation in all subsequent analyses. Vertical lines represent the average maximal epoch for each model, which for *deepbiosphere* was 4, 10 for *Inception V3*, and 45 for the climate-only *MLP* model.

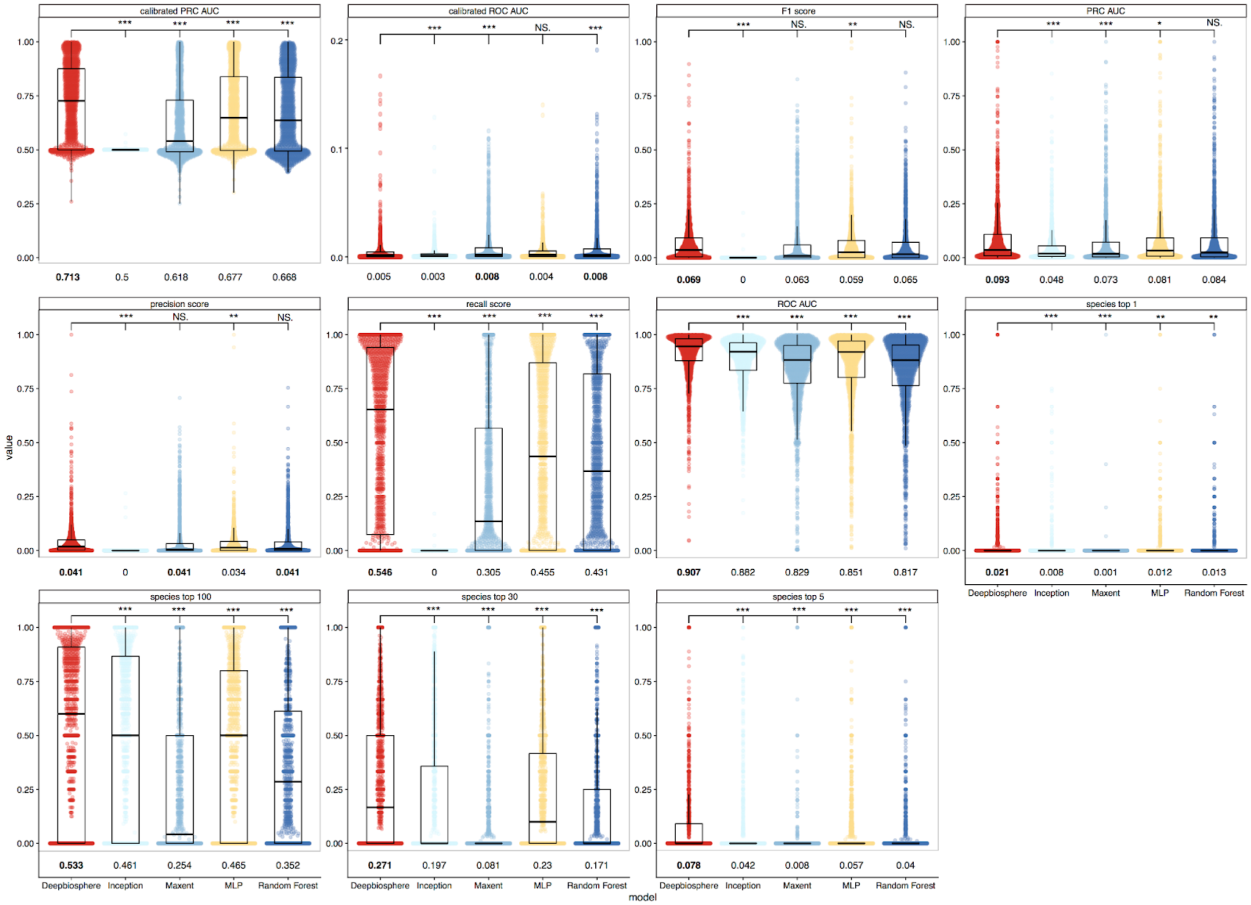


**Fig. S6 | Comparison of per-species accuracy metrics across species**

For the eleven accuracy metrics that can be calculated per-species, we report here the accuracy of *deepbiosphere* to baseline models on the uniform split of the dataset (**Fig. S4A**) using species found both in this train and test set split (1,541 species out of 2,221 total species in the dataset). On average, *deepbiosphere* outperforms the baseline approaches, having a higher accuracy on average for nine out of the eleven metrics reported here. Stars represent unpaired student *t*-test comparing *deepbiosphere's* accuracy per-species to the relevant baseline SDMs with *** indicating a *P*-value of $< 10^{-3}$, ** indicating a *P*-value of $< 10^{-2}$, and * indicating a *P*-value of $< 10^{-1}$. Values annotated below are the average accuracy per-species for that accuracy metric, bolded for the model with highest accuracy.

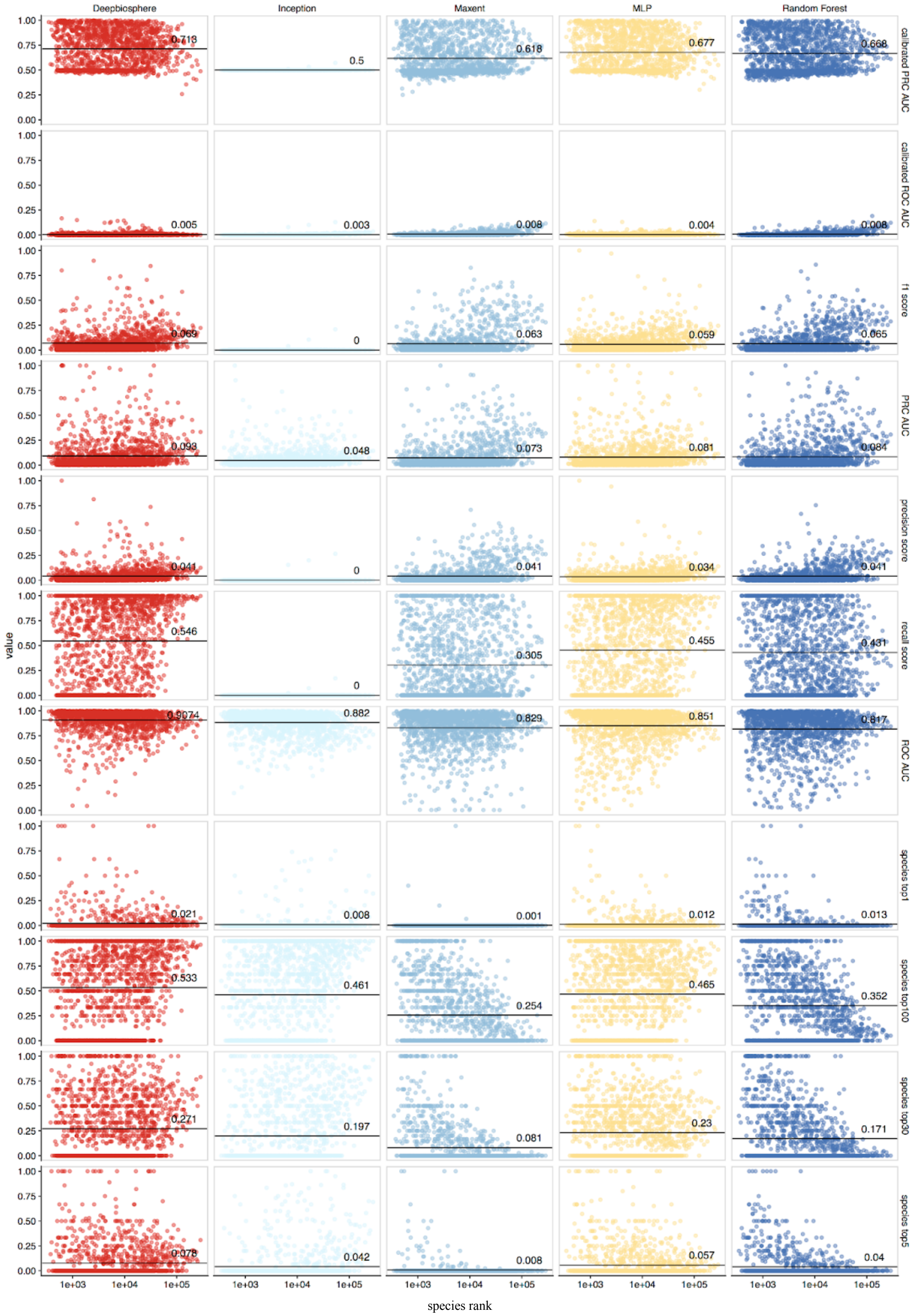


**Fig. S7 | Species abundance to accuracy relationship**

In order to better tease apart the relationship between commonness of observation and model accuracy, here we report the eleven per-species accuracy metrics by the species rank (number of times said species was observed in the whole dataset). Metrics reported here are for model accuracy on the uniform split of the dataset (**Fig. S4A**) using species found both in this train and test set split (1,541 species out of 2,221 total species in the dataset). Additionally, the average for each accuracy metric is reported and plotted as a solid grey line in each plot and annotated.

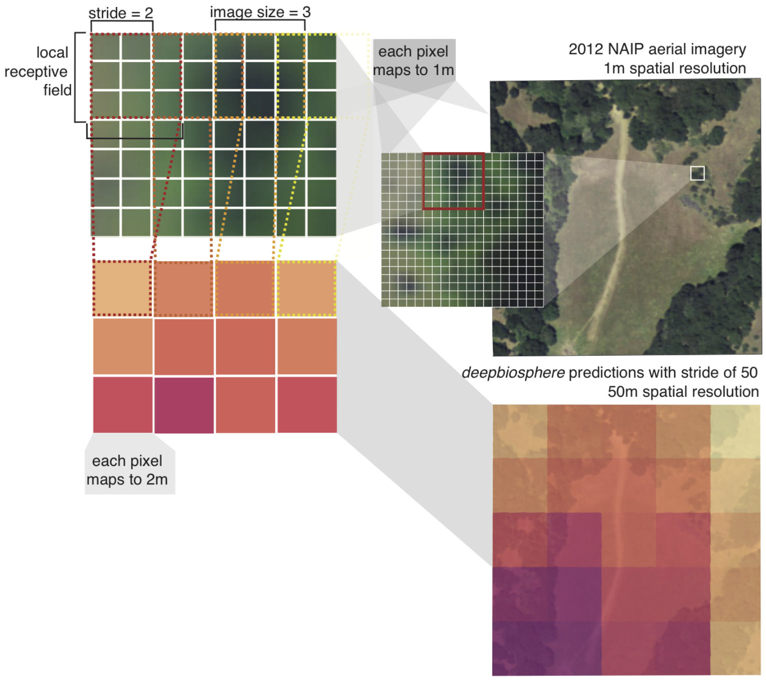


**Fig. S8 | Generating high-resolution predictions with *deepbiosphere***

In order to generate high resolution predictions from *deepbiosphere*, a technique from computer vision called striding is employed. Since *deepbiosphere* is trained with images of size 256 x 256 pixels, we are limited to generating predictions of this size and shape. However, we can get a higher resolution by shifting *deepbiosphere's* local receptive field (the current 256 x 256 pixels used to make a prediction) by K pixels to generate a map of K m resolution. This iterative sliding of the local receptive field is similar to striding, a technique used within the architecture of many CNN models. For example, to generate a 2 m-resolution map from 1 m resolution imagery using a CNN that makes predictions using a 3 x 3 image, we would generate predictions for each 3 x 3 image within the map, striding the predictions by 2, resulting in a final map with a ground resolution of 2 m.

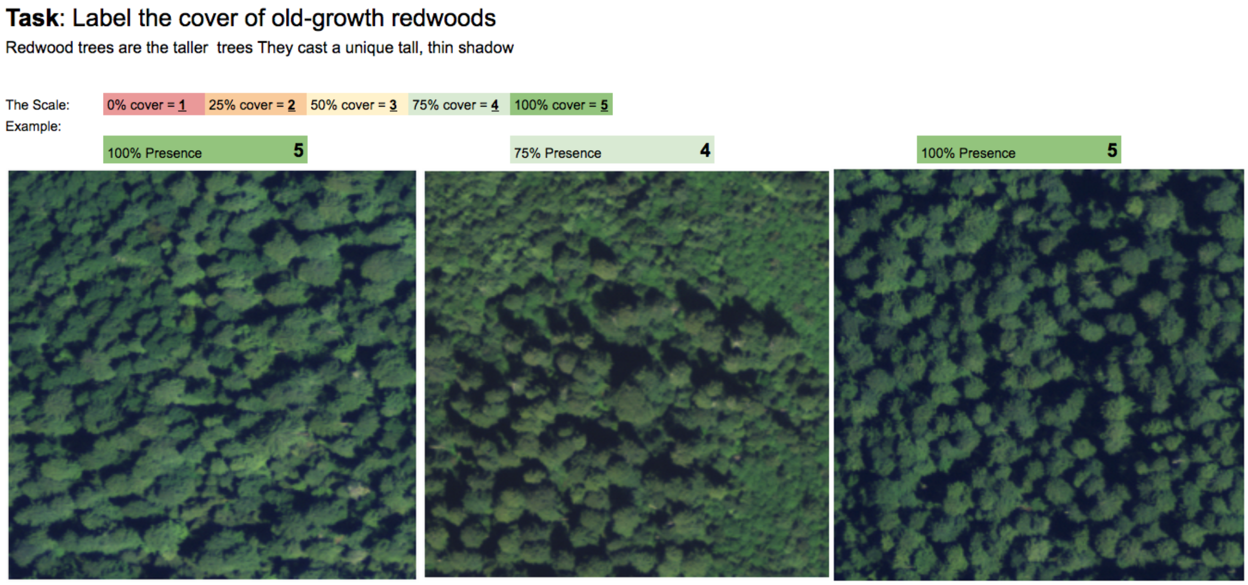


**Fig. S9 | Example images from human redwood labeling task.**
Labelers were given these three examples of old-growth redwoods as positive examples of redwood forest. Details of the three locations can be found in **Table S9**.

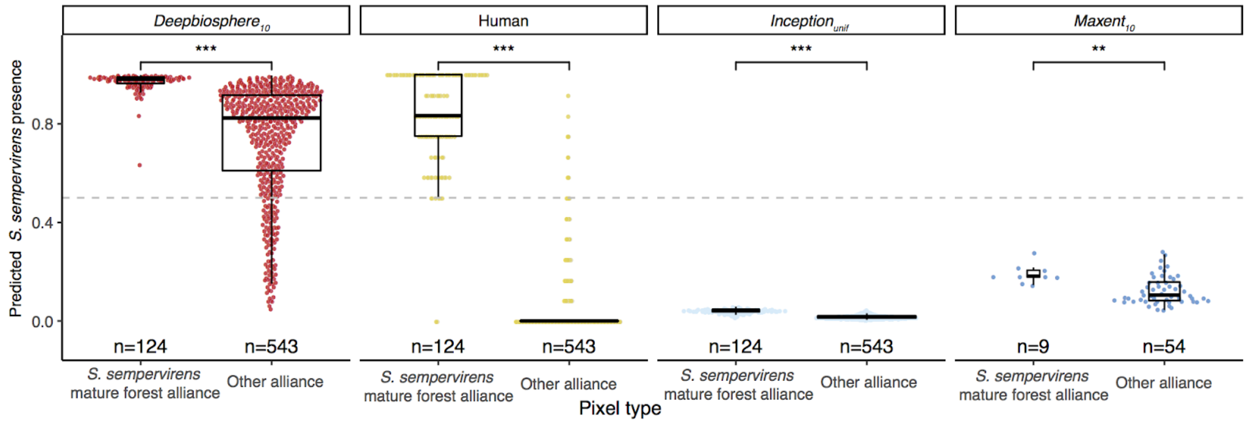


**Fig. S10 | Humans can correctly detect mature redwood groves**

When only looking at mature redwood groves (pixels coded as *S. semperivens* mature forest alliance using the official National Park Service vegetation map for the region, **Fig. S11A**), human annotators trained using the examples in **Fig. S9** do a better job at distinguishing mature redwood-coded pixels than any other model. Specifically, human annotations of redwood cover are the only annotations where, on average, the pixels coded as mature redwoods would be considered present (above 0.5, grey dotted line) while the pixels coded as any other vegetation type would be considered absent (below 0.5). *Deepbiosphere*$_{10}$ predicts non-mature pixels as containing redwoods, likely detecting pixels that contain secondary growth redwood groves, while both the *Inception*$_{unif}$ and *Maxent*$_{10}$ baselines fail to detect any redwoods in the area. While human annotators were highly accurate at detecting old-growth redwoods, the annotation process was slow and cumbersome, taking between 30 minutes to 2.5 hours for annotators to generate a 256 m-resolution estimate of redwood cover, while *deepbiosphere* took only 35 seconds to generate the same resolution predictions. Stars indicate an unpaired student's *t*-test with *P*-values as follows: *** <$10^{-3}$; ** <$10^{-2}$.

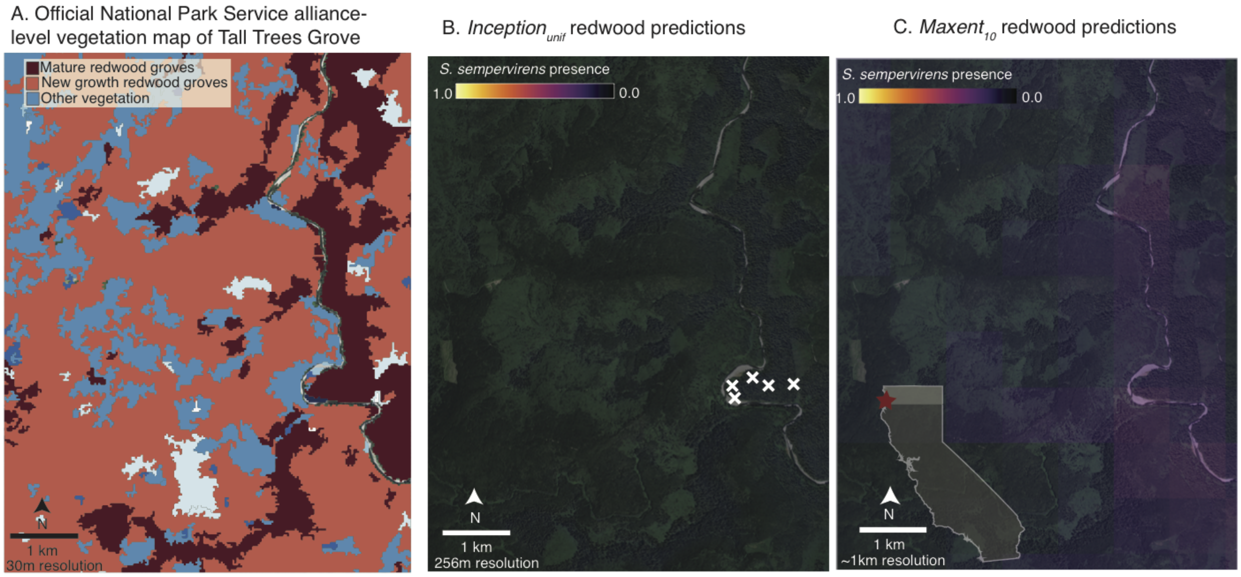


**Fig. S11 | Baseline SDMs cannot detect redwoods in Tall Trees Grove**
**(A)** Official National Park Service alliance-level vegetation map of study area[53]. Most of the cover in the area is predicted to be redwoods with a few other vegetation classes interspersed. **(B)** *Inception$_{unif}$* predictions of redwoods in study area. Since the *Inception* model is trained to predict one species exclusively at a time (courtesy of the softmax transformation in the cross-entropy loss function), the model's outputs cannot be interpreted as species distribution maps reliably since by construction the model will never predict a probability of 0.5 or higher. Even in this example where *Inception$_{unif}$* has been trained using redwood observations from inside the study area (white x), *Inception$_{unif}$* still doesn't predict any pixel as present, even for pixels it has seen as containing redwoods before. **(C)** *Maxent$_{10}$* predictions of redwoods in the study area. *Maxent* also does not predict any pixel as present in the study area, likely because it was not trained with any observations in the 10th spatial cross-validation band (light grey region in inset). While *Maxent* can reliably extrapolate to new regions if those regions have a similar climate profile to the regions used for fitting, the southern and northern populations of redwoods occupy quite dissimilar climates, with the northern population largely covered by the 10th cross-validation band living in a much cooler and at times wetter environment than the populations in the Central Coast and Santa Cruz mountains. Meanwhile, *deepbiosphere$_{10}$* can extrapolate to this new region because while the climate of the two major redwood populations differs dramatically, the 2D shape of the redwoods' crowns does not (see **Fig. S9** for examples), thereby enabling *deepbiosphere$_{10}$* to detect these redwood stands with high precision.

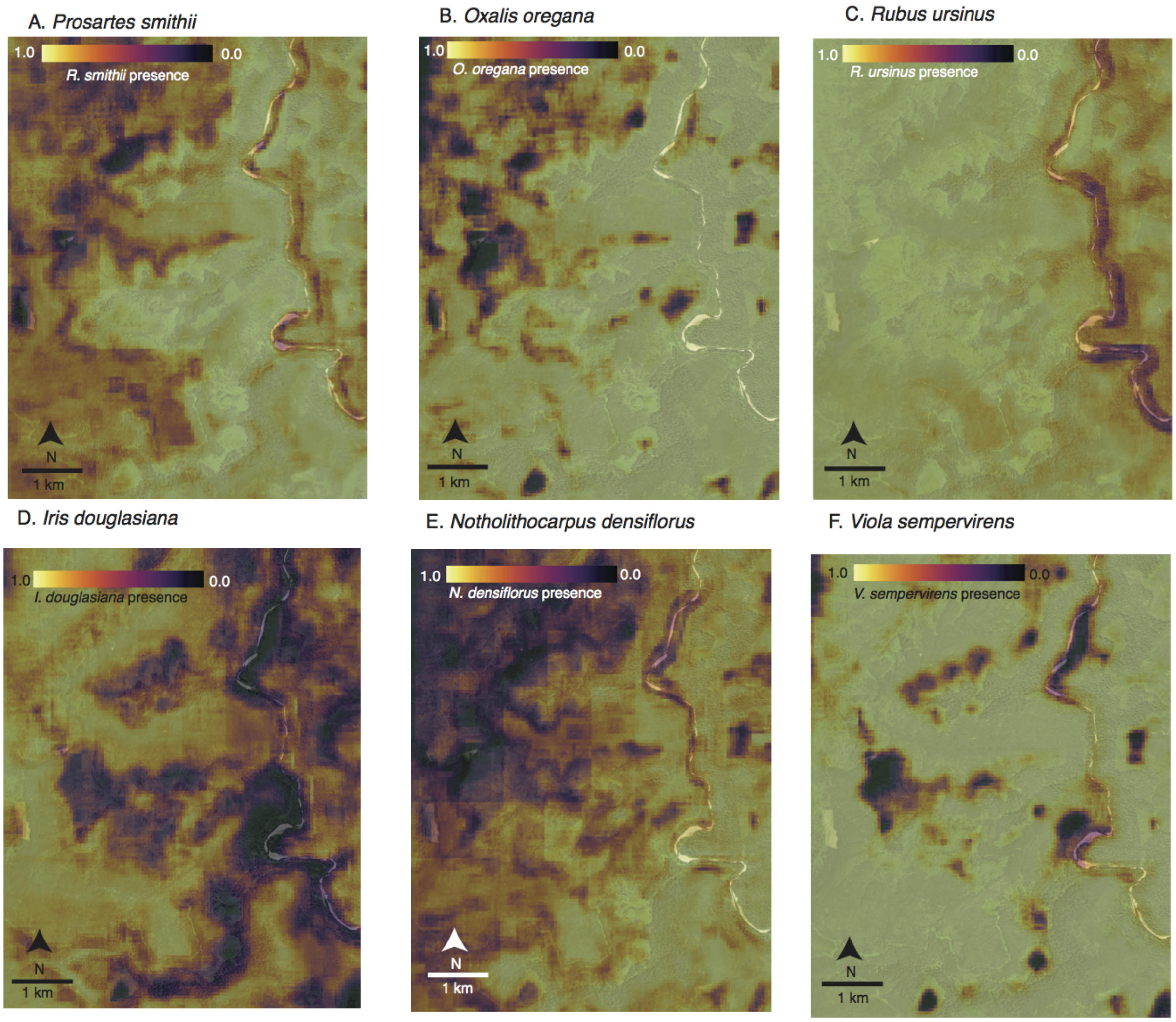


**Fig. S12 | *Deepbiosphere*-generated species presence maps for six redwood co-occurring species**
**(A,B)** Species presence maps for *Oxalis oregana* and *Prosartes smithii*, respectively. In ref. [53], both species were shown to have a high constancy (>=60%) in the mature redwood field plots and a low constancy (<=50%) in secondary growth redwood field plots. This difference in constancy implies that *O. oregana* and *P. smithii* tend to associate more with mature redwoods rather than secondary growth redwoods. **(C,D)** Species presence maps for *Rubus ursinus* and *Iris douglasiana*, respectively. In ref. [53] both species were shown to have a high constancy (>=60%) in the secondary growth redwood field plots and a low constancy (<=50%) in mature redwood field plots. This difference in constancy implies that *R. ursinus* and *I. douglasiana* tend to associate more with secondary redwood groves grown back after the clear-cutting of the 20th century, rather than the old-growth mature redwood groves. **(E,F)** Species presence maps for *Notholithocarpus densiflorus* and *Viola sempervirens*, respectively. In ref. [53], both species were shown to have a high constancy (>=60%) in both secondary growth redwood field plots and in mature redwood field plots. This implies *N. densiflorus* and *V. sempervirens* tend to generally associate with any redwood grove.

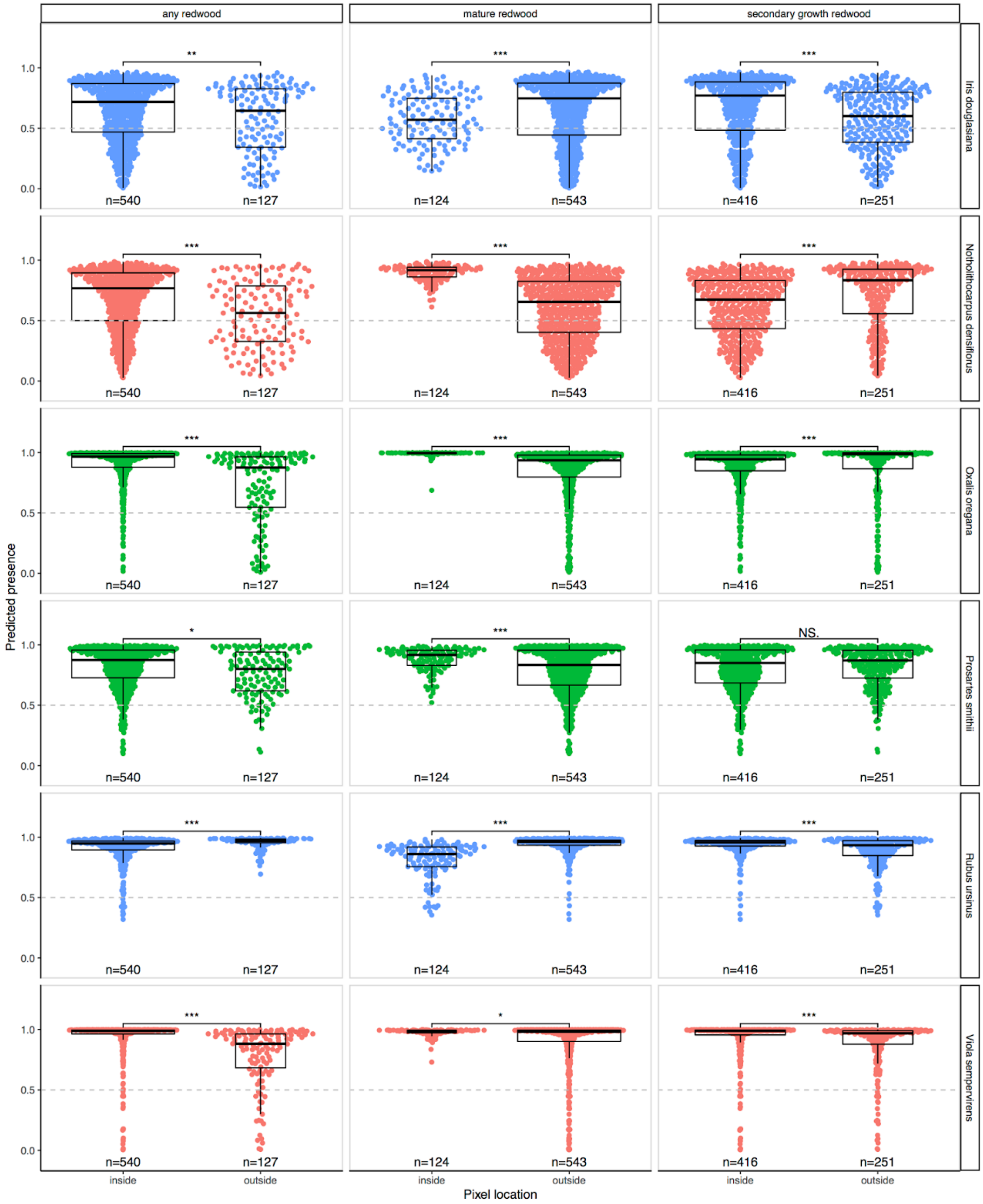


**Fig. S13 | Comparison of co-occurring species presence predictions for associated species**

For six understory species co-occurring with redwoods, here we compare the per-pixel predicted species presence. Species that tend to associate with old-growth redwoods have a higher predicted presence in mature redwood groves and a lower predicted presence in regrowth redwood groves, while conversely species that tend to associate with regrowth tend to have a higher predicted presence in secondary regrowth redwood groves and a lower predicted presence in mature redwood groves. Pixels from the "any redwood" category are considered "inside" if the largest vegetation class from the official NPS alliance-level vegetation map for said cell is either *Sequoia sempervirens*-(Other) YG Mixed Forest or *Sequoia sempervirens* Mature Forest. Pixels from the "mature redwood" category are considered "inside" if the largest vegetation class from the official NPS alliance-level vegetation map for said cell is *Sequoia sempervirens* Mature Forest. Pixels from the "secondary growth" category are considered "inside" if the largest vegetation class from the official NPS alliance-level vegetation map for said cell is *Sequoia sempervirens*-(Other) YG Mixed Forest. Stars indicate an unpaired student's *t*-test with *P*-values as follows: *** $<10^{-3}$; ** $<10^{-2}$; * $<10^{-1}$; N.S. $>= 10^{-1}$

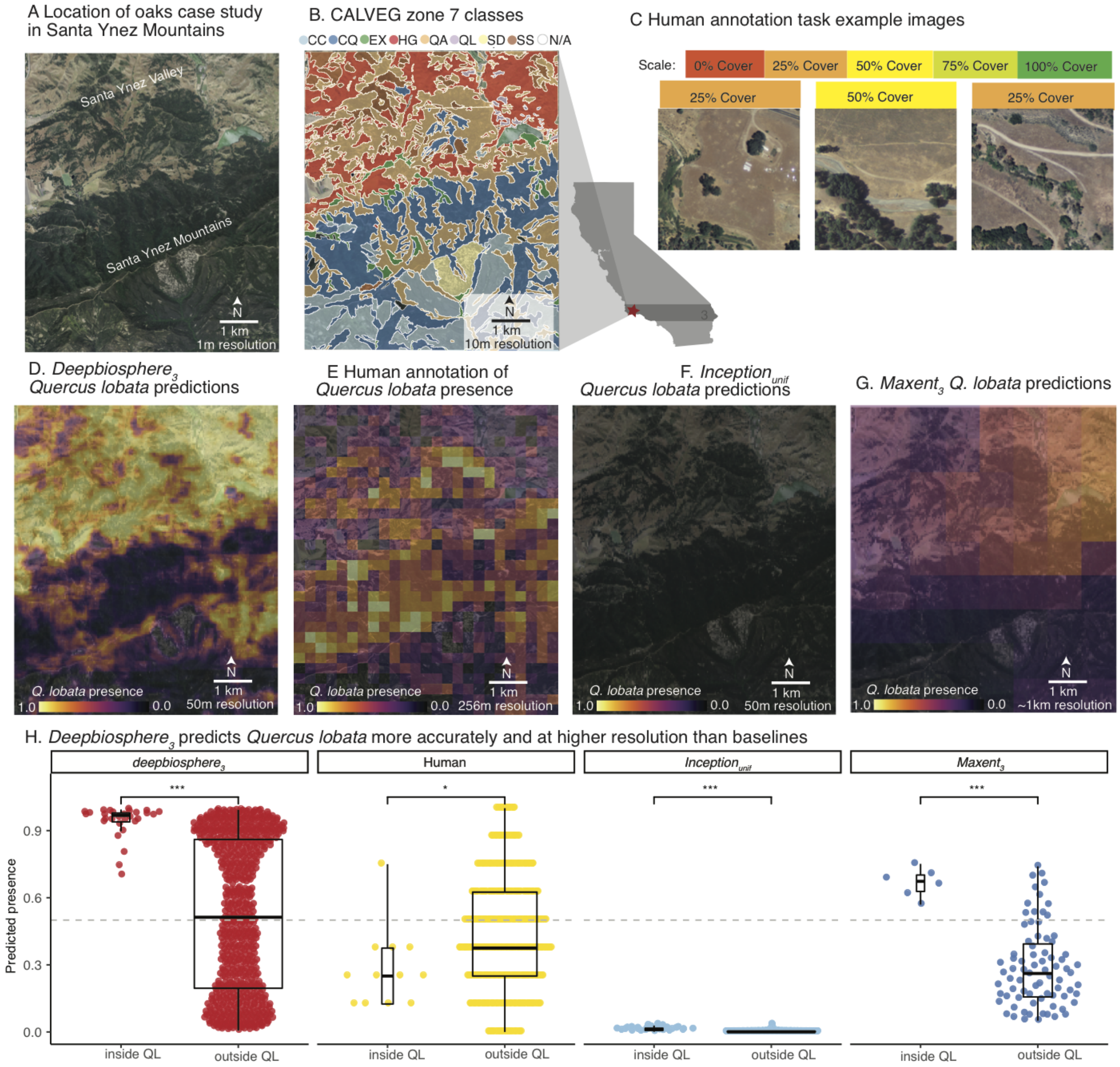


**Fig. S14 | Predictions of dominant species in the Santa Ynez Valley and Mountains of Southwest California.**
**(A)** The Santa Ynez mountains of Southern California have a rich and varied ecological landscape, with many rapid ecosystem transitions from the dry interior valley savanna to the scrubby mountain peaks of the Santa Ynez mountains, making it a good case study to explore how well *deepbiosphere* can capture spatial variation in chaparral ecosytems. **(B)** CALVEG zone 7 alliance-level vegetation classes for study area[54]. Only zones mapping to native vegetation with a sufficient cover in the study area are shown. **(C)** *Quercus lobata* annotation task examples. Detailed information about sites can be found in **Table S9**. **(D)** *Deepbiopshere$_3$* prediction of *Q. lobata* across the study area. Generally speaking, *deepbiosphere$_3$* predicts *Q. lobata* as present in the Santa Ynez valley —its native habitat— and absent in the Santa Ynez mountains. **(E)** Human cover annotations of *Q. lobata* across study area. Human labelers struggled with correctly labeling squares where *Q. lobata* is expected to be found, favoring the chaparral scrub of the Santa Ynez foothills over the valley floor where the oaks are actually found. **(F)** The *Inception$_{unif}$* baseline does not predict *Q. lobata* present anywhere within the study area, a consequence of its softmax-based loss function. **(G)** The *Maxent$_3$* baseline does predict *Q. lobata* as present in most of the valley pixels, but has a much lower resolution than *deepbiosphere$_3$* making it impossible for the model to disambiguate the wooded hills centered in the valley from the grassland portions of the valley. **(H)** Comparison of *Q. lobata* presence annotations per-pixel shows that *deepbiosphere$_3$* and *Maxent$_3$* are the only SDMs that correctly disambiguate pixels annotated as *Q. lobata* habitat (QL) by the CALVEG vegetation map from those not annotated as such. Pixels are labeled as *Q. lobata* habitat if any part of the pixel intersects any CALVEG polygon of class QL. Stars indicate an unpaired student's *t*-test

per-pixel where *** means a *P*-value of $<10^{-3}$ and ** means a *P*-value of $<10^{-2}$. Pixels are calculated at 256 m resolution or lower to minimize spatial autocorrelation.

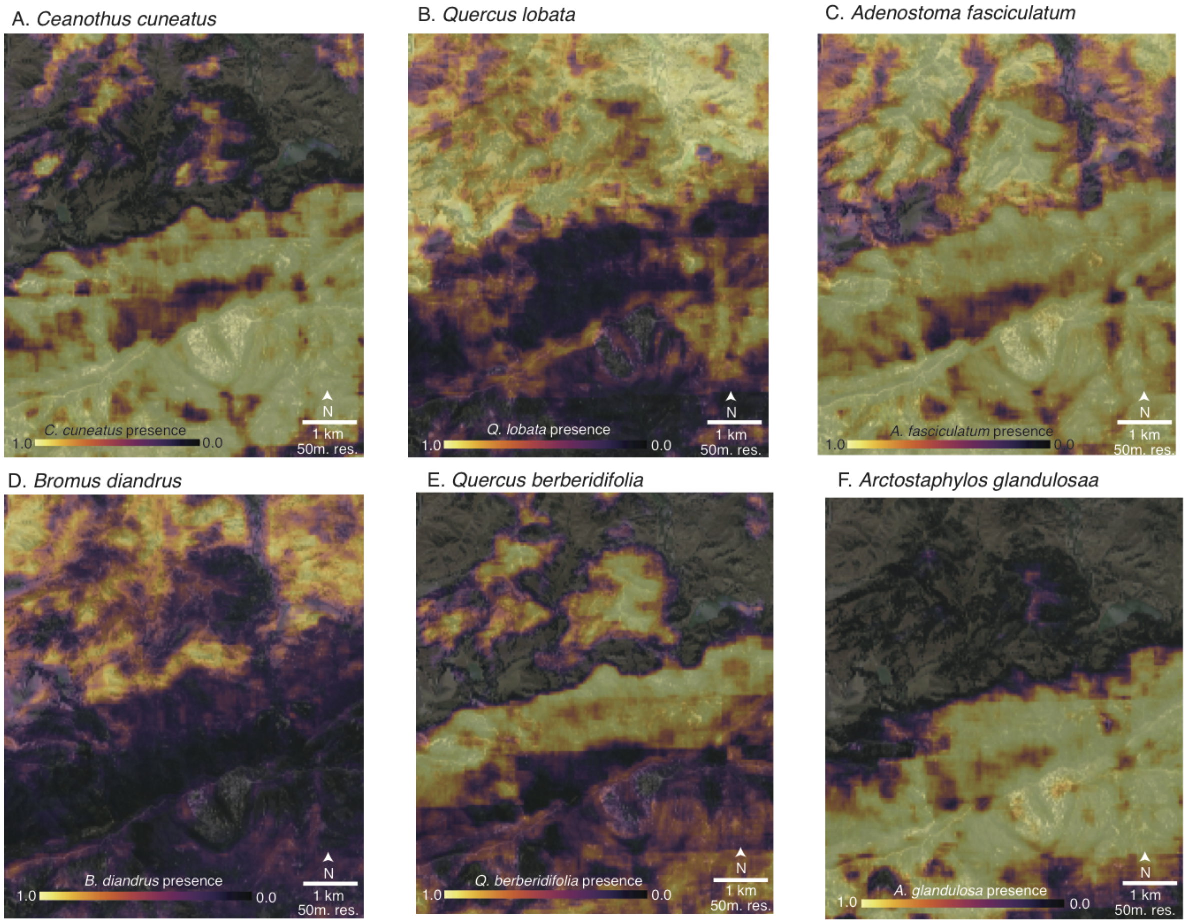


**Fig. S15 | *Deepbiosphere*-generated species presence maps for six chaparral indicator species**
**(A)** Species presence map for *Ceanothus cuneatus* (buckbrush), an indicator or associated species for CALVEG zones *Ceanothus* chaparral alliance (CC), lower montane mixed chaparral alliance (CQ), and coastal mixed hardwood alliance (EX). **(B)** Species presence map for *Quercus lobata* (valley oak), an indicator species for the valley oak alliance (QL). **(C)** Species presence map for *Adenostoma fasciculatum* (chamise), an indicator or associated species for the coast live oak alliance (QA), *Ceanothus* chaparral alliance (CC), coastal mixed hardwood alliance (EX), lower montane mixed chaparral alliance (CQ), and California sagebrush alliance (SS). **(D)** Species presence map for *Bromus diandrus* (great brome), an indicator species for the annual grasses and forbs alliance (HG). **(E)** Species presence map for *Quercus berberidifolia* (scrub oak) an associated species in the lower montane mixed chaparral (CQ) alliance. **(F)** Species presence map for *Arctostaphylos glandulosa* (eastwood manzanita), an associated species in the lower montane mixed chaparral (CQ) and manzanita chaparral (SD) alliances.

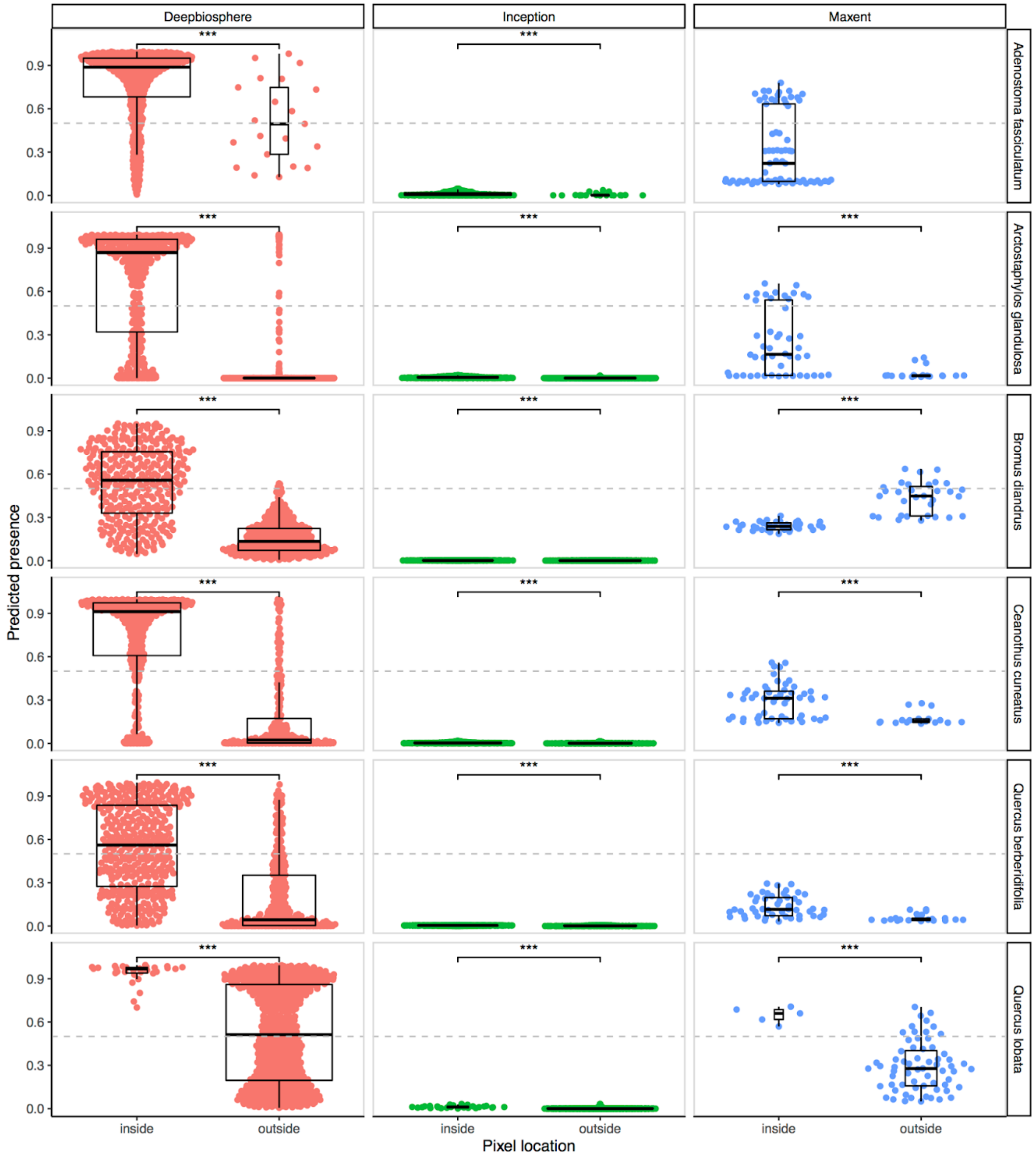


**Fig. S16 | Comparison of predicted presence of indicator species to CALVEG vegetation mapping**

Comparing the predicted probabilities per-pixel generated by *deepbiosphere*$_3$ for each of the six indicator species using the USDA Forest Service's South Coast existing vegetation map[54] shows that on average, *deepbiosphere*$_3$ does a good job of correctly predicting species as present in the pixels located within their associated CALVEG habitats (**Fig. S14B**). Specifically, all species are predicted as present (above 0.5) on average for pixels inside the associated habitats and as absent (below 0.5) on average for pixels outside of the associated habitats. Meanwhile, both *Maxent*$_3$ and *Inception*$_{unif}$ fail this test for all but one case (*Maxent*$_3$ modeling *Q. lobata*). Pixels are labeled as inside habitat if any part of the pixel intersects any CALVEG polygon from the USDA Forest Service's South Coast existing vegetation map[54] where said species is mentioned in the vegetation description for that class[55]. Stars indicate an unpaired student's *t*-test per-pixel where *** means a *P*-value of $<10^{-3}$ and pixels are calculated at 256 m resolution or lower to minimize spatial autocorrelation.

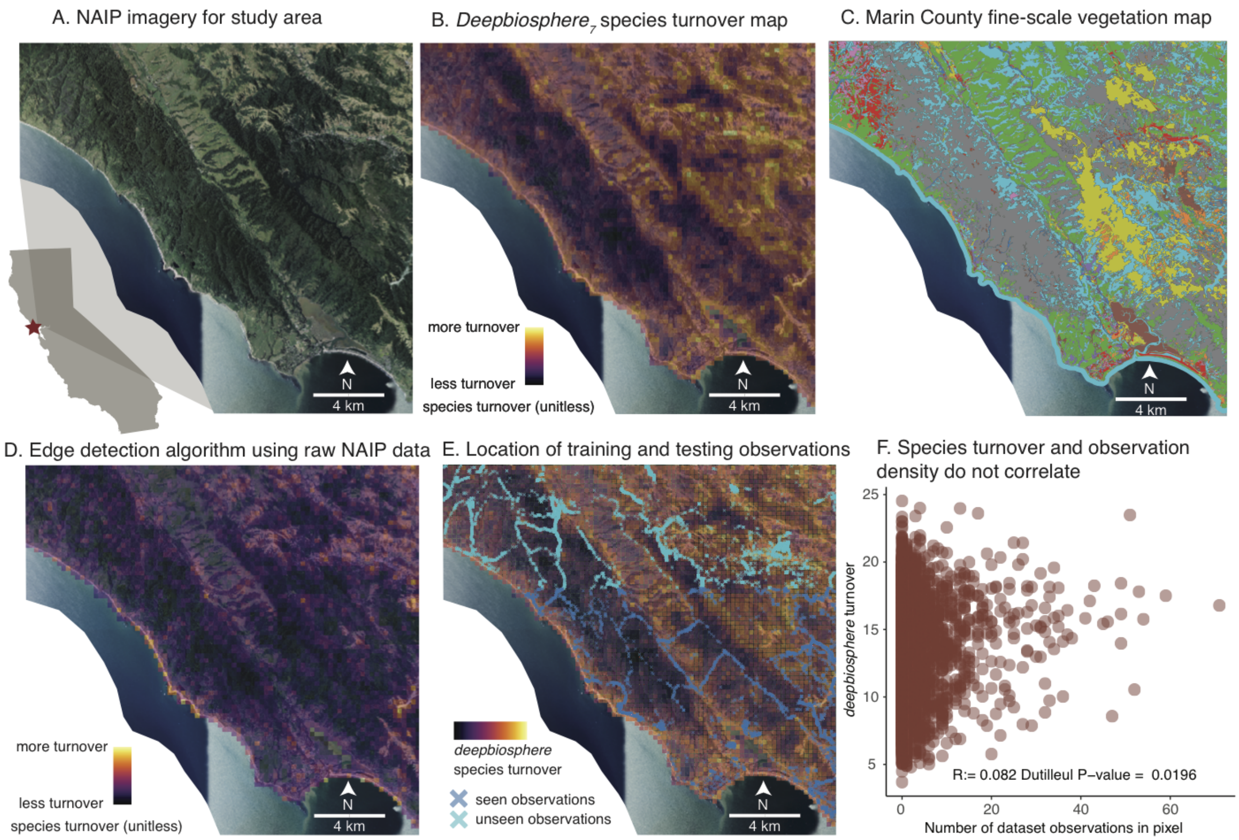


**Fig. S17 | *Deepbiosphere* prediction of spatial community change in northern Marin county.**
**(A)** This region of Marin county is located at the boundary between the Coast Range and Central California Foothills and Coastal Mountains level 1 EPA ecoregions, thus making it an ideal location to study spatial transitions in plant communities. NAIP aerial imagery of the area confirms visually that the landscape is highly varied with beach, forest, and grassland ecosystems visible. **(B)** Using the edge detection algorithm from **Fig. S19**, regions of high species turnover can be visually located. Specifically, pixels corresponding to areas along the beach and between the grassland and forest zones have a higher intensity, which conforms to intuition that boundaries between communities have high spatial community change and rapid species turnover. The spatial community change metric is unitless. **(C)** An independent fine-scale map of vegetation and land use in the area confirms that this area indeed contains many unique ecosystems[58]. The color key can be found in **Fig. S18**. **(D)** Running the same edge detection algorithm but using the raw NAIP bands as input (A) leads to this map of average difference in color between nearby NAIP pixels. Visually, the map looks much different from the *deepbiosphere*-generated version, confirming that *deepbiosphere* is doing far more than making trivial predictions from the greenness of the pixels or other simplistic associations. **(E)** Mapping the location of observations both seen and unseen during training shows that the spatial community change visually also does not appear to naively recapitulate locations it has seen before. **(F)** Correlating the number of observations per-pixel with the spatial community change as predicted by *deepbiosphere*$_7$ confirms that the spatial community change metric is capturing real changes in the community rather than spurious correlations to either the raw pixel greenness or previously seen observations.

- Acacia spp. – Grevillea spp. – Leptospermum laevigatum Semi-Natural Alliance
- Acer macrophyllum Association
- Acer macrophyllum – Alnus rubra Alliance
- Acer negundo / (Rubus ursinus) Association
- Adenostoma fasciculatum Alliance
- Aesculus californica Alliance
- Alnus rhombifolia Alliance
- Ammophila arenaria Semi-Natural Alliance
- Annual Cropland
- Arbutus menziesii Alliance
- Arctostaphylos (bakeri, montana) Alliance
- Arctostaphylos (nummularia, sensitiva) – Chrysolepis chrysophylla Alliance
- Arctostaphylos glandulosa Alliance
- Arid West Freshwater Marsh Group
- Artemisia californica – (Salvia leucophylla) Alliance
- Artemisia pycnocephala Association
- Baccharis pilularis Alliance
- Barren and Sparsely Vegetated
- Bolboschoenus maritimus Alliance
- Calamagrostis nutkaensis Alliance
- Californian Annual & Perennial Grassland Mapping Unit
- Californian Cliff, Scree & Rock Vegetation Group
- Californian Vernal Pool / Swale Bottomland Group
- Ceanothus cuneatus Alliance
- Ceanothus thyrsiflorus Alliance
- Channel
- Conium maculatum – Foeniculum vulgare Semi-Natural Alliance
- Cortaderia (jubata, selloana) Semi-Natural Alliance
- Corylus cornuta / Polystichum munitum Association
- Cotoneaster (lacteus, pannosus) Provisional Semi-Natural Association
- Developed
- Distichlis spicata Alliance
- Eriophyllum staechadifolium – Erigeron glaucus – Eriogonum latifolium Alliance
- Eucalyptus (globulus, camaldulensis) Provisional Semi-Natural Assocation
- Forest Fragment
- Frangula californica ssp. californica – Baccharis pilularis / Scrophularia californica Association
- Fraxinus latifolia Alliance
- Gaultheria shallon – Rubus (ursinus) Alliance
- Genista monspessulana Semi-Natural Association
- Grindelia stricta Provisional Association
- Hesperocyparis macrocarpa Ruderal Provisional Semi-Natural Association
- Hesperocyparis sargentii / Ceanothus jepsonii – Arctostaphylos spp. Association
- Hesperocyparis sargentii Association
- Lupinus arboreus Alliance
- Lupinus chamissonis – Ericameria ericoides Alliance
- Major Road
- Mesembryanthemum spp. – Carpobrotus spp. Semi-Natural Alliance
- Mudflat/Dry Pond Bottom Mapping Unit
- Non-native Forest
- Non-native Herbaceous
- Non-native Shrub
- Notholithocarpus densiflorus Alliance
- Nursery or Ornamental Horticulture Area
- Orchard or Grove
- Perennial Cropland
- Pinus muricata – Pinus radiata Alliance
- Pinus radiata Plantation Provisional Semi-Natural Association
- Pseudotsuga menziesii Mapping Unit
- Pseudotsuga menziesii – Notholithocarpus densiflorus Mapping Unit
- Quercus agrifolia Alliance
- Quercus chrysolepis Alliance
- Quercus durata Alliance
- Quercus garryana Alliance
- Quercus kelloggii Alliance
- Quercus lobata Alliance
- Quercus wislizeni – Quercus chrysolepis (shrub) Alliance
- Rubus armeniacus Semi-Natural Association
- Salix gooddingii – Salix laevigata Alliance
- Salix lasiolepis Alliance
- Salix lucida ssp. lasiandra Association
- Sarcocornia pacifica (Salicornia depressa) Alliance
- Sequoia sempervirens Alliance
- Shrub Fragment
- Spartina foliosa Association
- Toxicodendron diversilobum – (Baccharis pilularis) Association
- Umbellularia californica Alliance
- Vancouverian Freshwater Wet Meadow & Marsh Group
- Vineyard
- Water
- Western North American Freshwater Aquatic Vegetation Macrogroup

**Fig. S18 | Legend for fine-scale Marin vegetation map**
Color key for fine scale vegetation map of northern Marin county[58]. The map contains 80 unique classes, so a circular color scale was utilized to aid visualization.

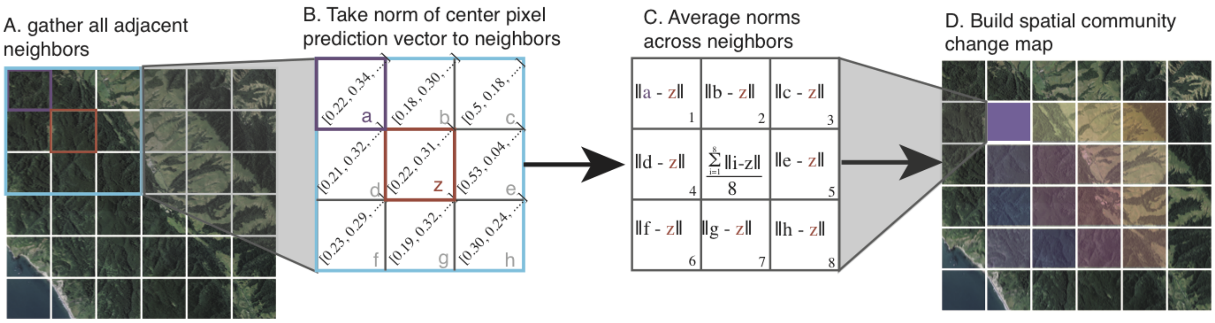


**Fig. S19 | Spatial community change algorithm visual explanation.**
Visual explanation of spatial community change detection algorithm inspired by edge detection filters from computer vision. **(A)** First, the algorithm identifies images adjacent to a given 256 x 256 m square **(B)** Next, using the predicted probabilities from *deepbiosphere,* the distance between each neighboring cell and the central pixel is calculated. **(C)** Then, the norm of these differences is used to generate the average local neighborhood change for the central pixel. (**D**) Convolving this norm-of-neighbors pixel-by-pixel generates the full map of spatial community change at the same resolution as the original map, with a one pixel buffer.

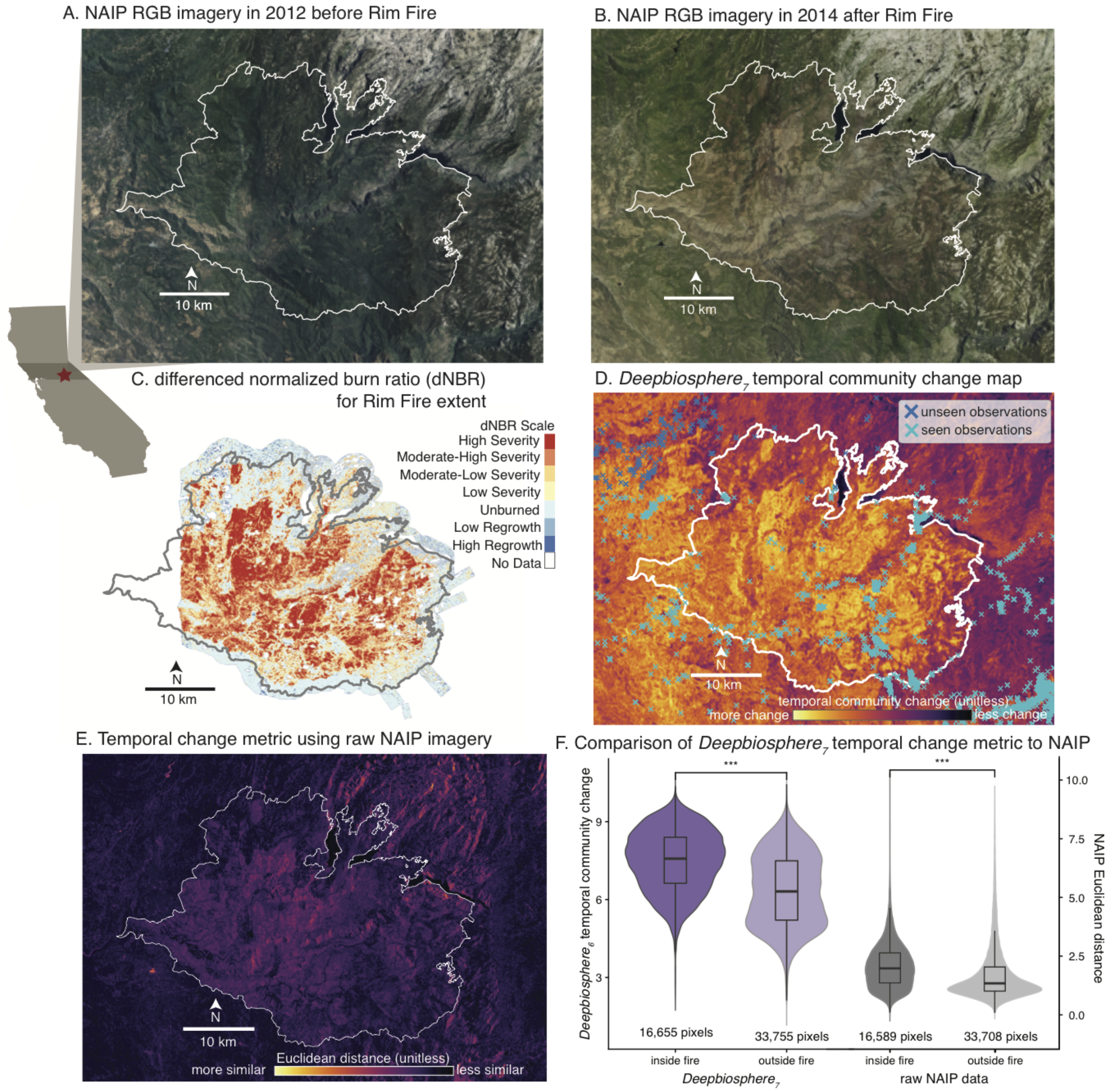


**Fig. S20 | Detecting rapid temporal plant community change after a major California wildfire**

**(A)** NAIP RGB imagery from the Sierra Mountain foothills in eastern California in 2012. This area was the site of a major wildfire in 2013 called the Rim Fire, which at the time was the second-largest wildfire in California history. **(B)** NAIP RGB imagery of 2014 for after the 2013 Rim Fire, with boundary outlined in white. Comparing (A) and (B), the burn scar of the wildfire is very visible within the fire perimeter compared to regions outside the burn scar. **(C)** Differenced normalized burn ratio (dNBR) for part of the fire area from ref. [64]. dNBR estimates the severity of a fire using a normalized scale based on differences in green and near infrared band wavelengths from hyperspectral data collected before vs. after the fire. Only part of the region was imaged before the fire, so dNBR coverage does not extend to the entire area. **(D)** Temporal community change as predicted by *deepbiosphere*$_7$ with observations used to train *deepbiosphere*$_7$ overlaid. **(E)** Temporal Euclidean distance calculated using raw NAIP RGB-I imagery visually does not appear to match the burn severity metric near as well as the *deepbiosphere*-based temporal community change. This is especially interesting given that the infrared and green NAIP bands used to calculate the Euclidean distance are nearly the same wavelength as the bands used to calculate dNBR. **(F)** Comparing pixels from inside vs. outside the fire, both the *deepbiosphere*-based and NAIP-based temporal Euclidean distances are on average higher within versus outside the fire bounds as expected (unpaired student's *t*-test; *P*-values $< 2.2 \times 10^{-16}$).

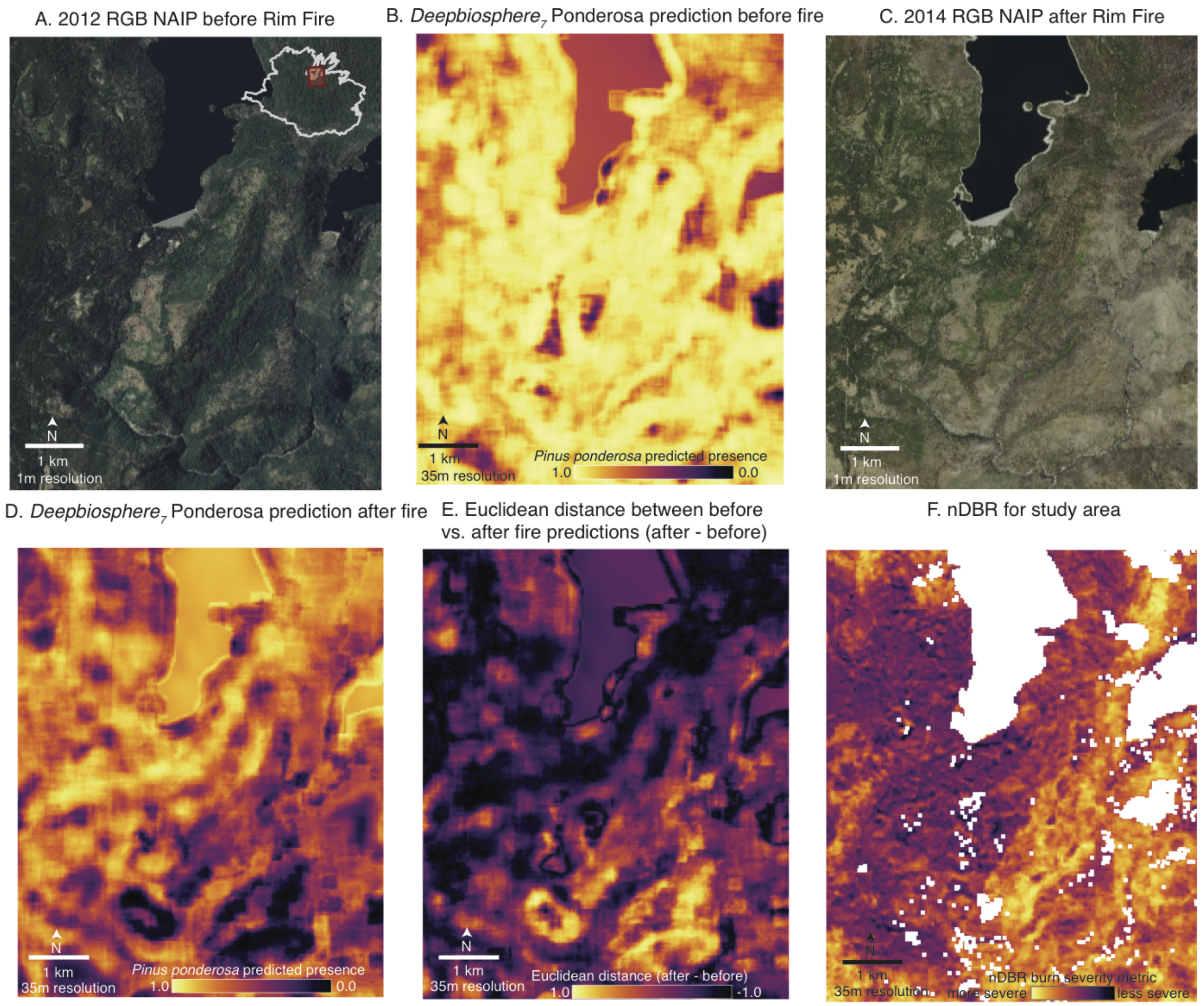


**Fig. S21 | Example of temporal Euclidean distance calculation**
**(A)** NAIP imagery before fire in a subset of the fire's extent (location inside perimeter of fire inset). **(B)** *Deepbiosphere*$_7$ predicted probabilities for *Pinus ponderosa* (ponderosa pine) before the fire, using the imagery in (A). **(C)** NAIP imagery after the fire from the same geographic location. Visually, locations in the bottom right of the image appear to have suffered the most severe burning, while forests on the left hand side of the area have been somewhat spared. **(D)** *Deepbiosphere*$_7$ predicted probabilities for *P. ponderosa* after the fire. Predicted probabilities on the left hand side of the area have not changed substantially while the more severely burned lower right quadrant now has *P. ponderosa* predicted mostly as absent. **(E)** To capture this temporal change in *P. ponderosa* presence, the Euclidean distance between the probabilities in (B) and (D) are calculated per-pixel. For the one-dimensional case (only one species at a time) the Euclidean distance simplifies to the temporal difference between $\hat{p}_{2012}$ and $\hat{p}_{2014}$. Extending this difference calculation across all species in the dataset and taking the subsequent norm is how the temporal community change metric is generated, capturing both the magnitude and direction of probability shifts across species and across time. (F) Here, an empirical metric of burn severity—difference in normalized burn ratio (dNBR)[64]—is displayed for the region.

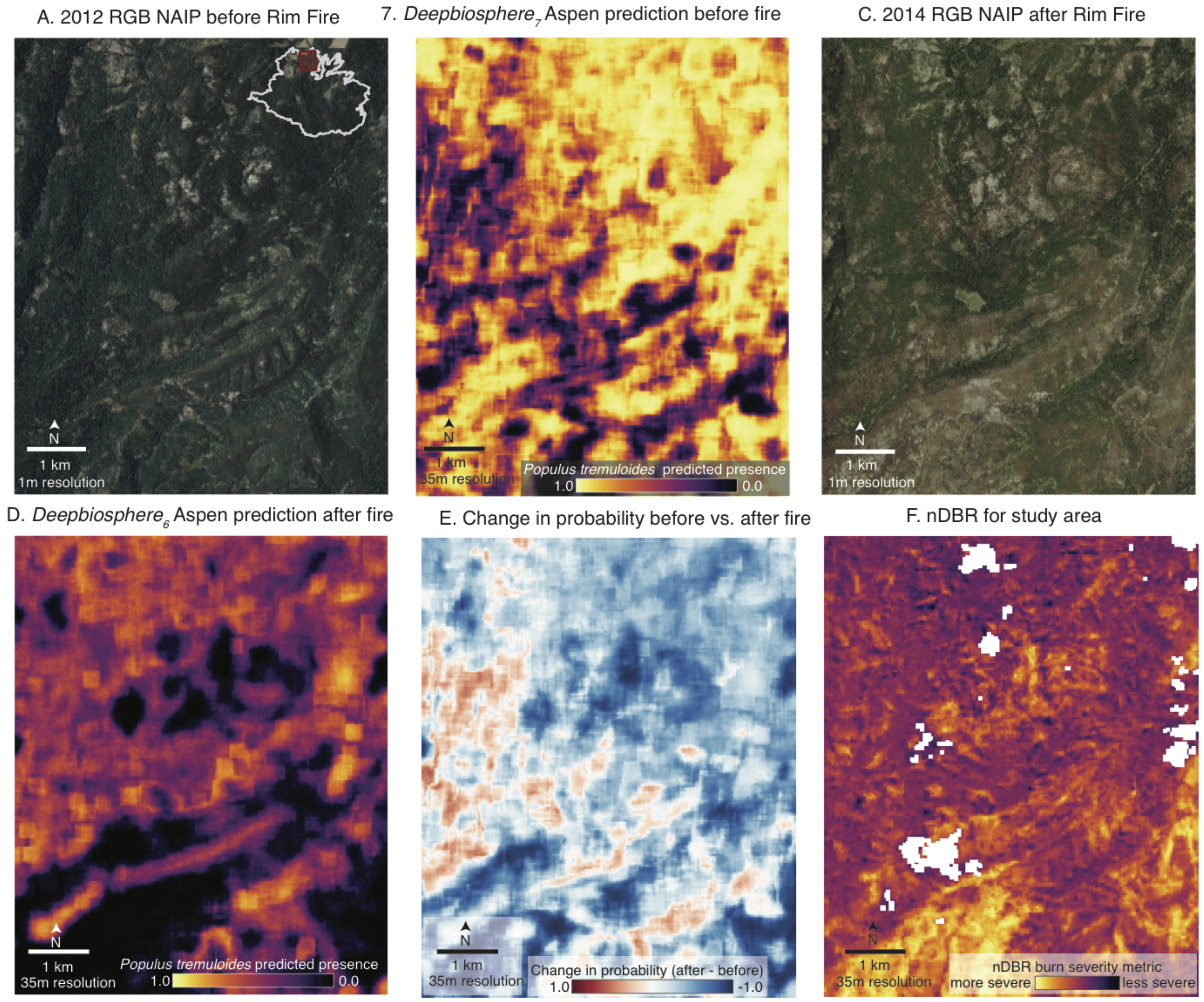


**Fig. S22 | Decrease in predicted *Populus tremuloides* presence before and after the Rim Fire.**
**(A)** NAIP imagery before the fire; example location inset. **(B)** *Deepbiosphere* predictions of Aspen (*P. tremuloides*) presence before the fire, using the imagery from (A). Lighter colors indicate areas *deepbiosphere* predicts Aspen are likely present while darker colors indicate areas of likely absence. **(C)** NAIP imagery from after the fire. **(D)** *Deepbiosphere* predictions of Aspen presence after the fire, using the same scale as (B). **(E)** Predicted change in Aspen presence (calculated by taking the difference between images (D) and (B)). Although Aspen can be found in fire-prone ecosystems, their canopy burns with only the root system surviving and resprouting in the future[69]. Thus we expect an immediate decrease in their predicted presence after a fire until a few years later after which suckers start to re-emerge from the root system. **(F)** The difference in normalized burn ratio (dNBR), which estimates the fire's severity, visually appears to show a more severe burn occurred where predicted Aspen presence has decreased substantially, as would be expected based on Aspen's rhizomatic fire response strategy.

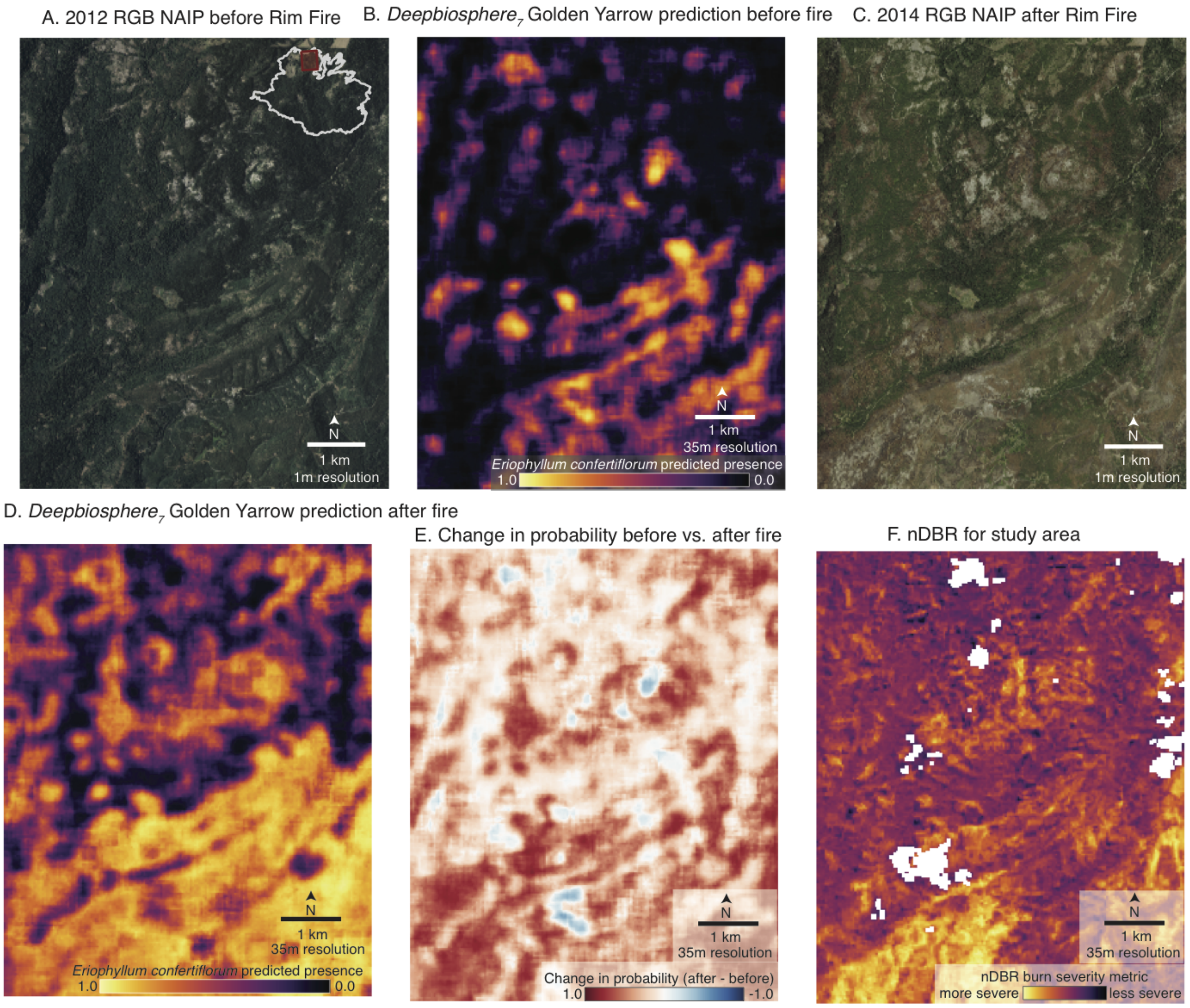


**Fig. S23 | Increase in predicted *Eriophyllum confertiflorum* presence before and after the Rim Fire.**
**(A)** NAIP imagery before the fire; **(B)** *deepbiosphere$_7$* predictions of Golden Yarrow (*Eriophyllum confertiflorum*) presence before the fire, using the imagery from (A). Lighter colors indicate areas *deepbiosphere$_7$* thinks Golden Yarrow are likely present while darker colors indicate areas of likely absence. **(C)** NAIP imagery from after the fire. **(D)** *deepbiosphere$_7$* predictions of Golden Yarrow presence after the fire, using the same scale as (B). **(E)** predicted change in Golden Yarrow presence (calculated by taking the difference between images (D) and (B)). Golden Yarrow is an herbaceous shrub often found in large clumps and is a well known "fire follower" species that follows a recruitment fire adaptation strategy, often rapidly increasing in abundance after a fire event[70]. We thus expect to see a marked increase of Golden Yarrow presence after the fire, aligning with evidence that fire improves Golden Yarrow seed germination rate and increases its abundance[71]. **(F)** The difference in normalized burn ratio (dNBR) for the study area, which estimates the fire's severity.

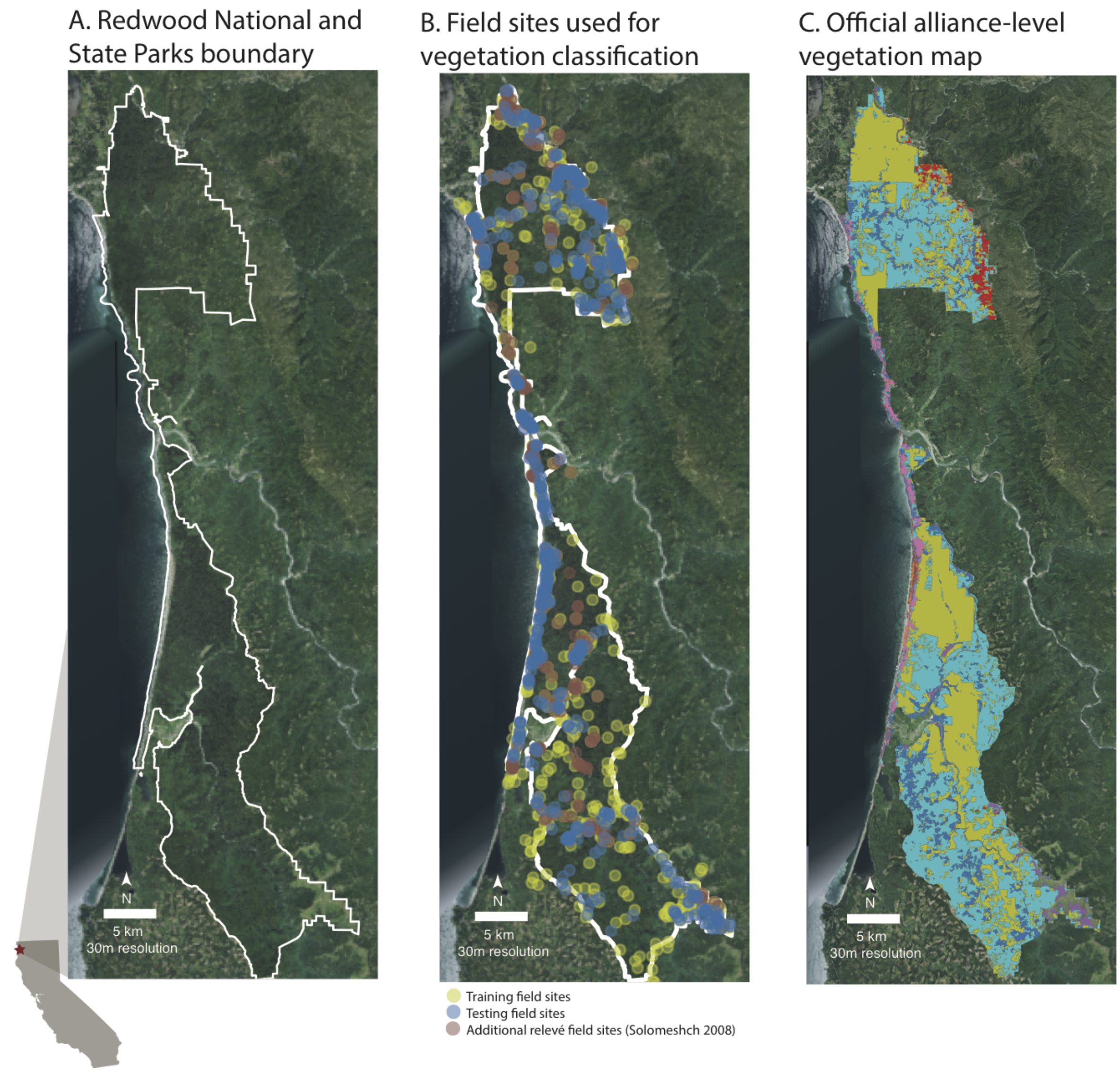


**Fig. S24 | Classifying alliance-level vegetation in Redwoods National & State Parks using *deepbiosphere***
**(A)** NAIP RGB imagery of Redwoods National and State Parks (RNSP) in northwestern California. One of the last remaining vestiges of old-growth redwood's range, RNSP is a critical refuge located in the heart of redwood's native range. The white outline demarcates the joint boundary of the two parks. **(B)** Locations of field plots used by the National Park Service (NPS) for generating an alliance-level vegetation map of the park. 423 field plots (yellow circles) were used to fit the models used to generate the final RNSP vegetation map[53]. An additional 286 field plots (red circles) were used to generate the alliance-level vegetation classes for the park in 2008. Finally, 489 field plots were used to perform an accuracy assessment of the resultant vegetation map. **(C)** The official NPS alliance-level vegetation map of RNSP. Here, the 30 approximately alliance-level map classes used to assess the vegetation map are displayed (see Section 6.2 of ref. [53] for class details).

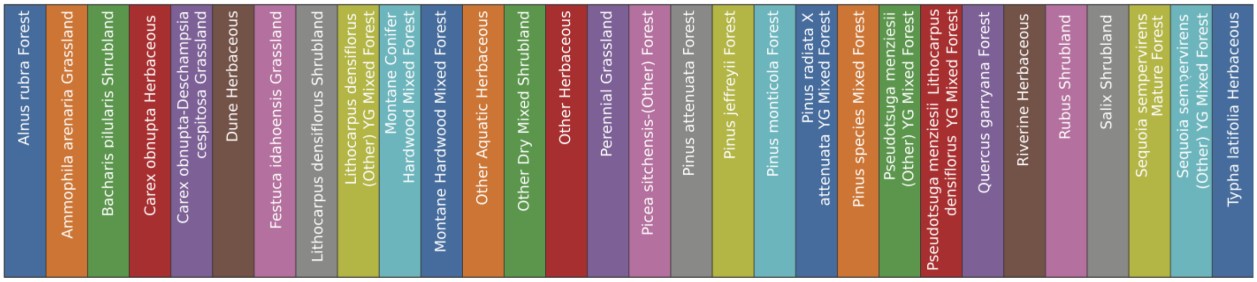


**Fig. S25 | Class map for alliance-level vegetation map of Redwoods National and State Parks**
Color key for alliance-level vegetation map of Redwoods National and State Parks[53].

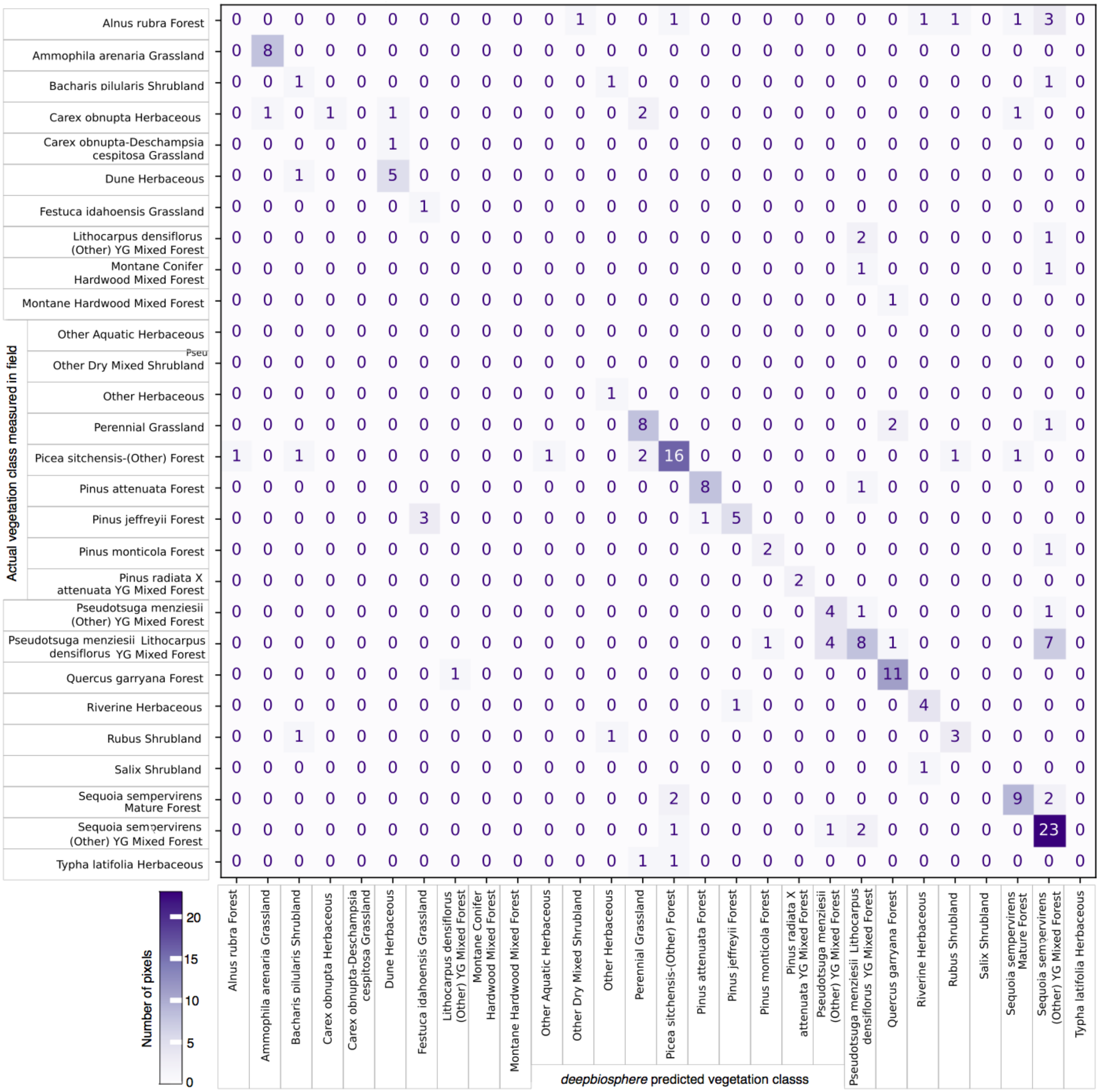


**Fig. S26 | Confusion matrix of *deepbiosphere*-based vegetation classifier on held-out field plots**

Confusion matrix of the predictions of the random forest feature extractor fitted using *deepbiosphere*$_{10}$ features with the highest single-label accuracy fold from the ten-fold cross-validation trials. The single-label accuracy of the classifier was 63.49% for the 189 held-out test field plots fitted using all 955 *deepbiosphere*-based features.

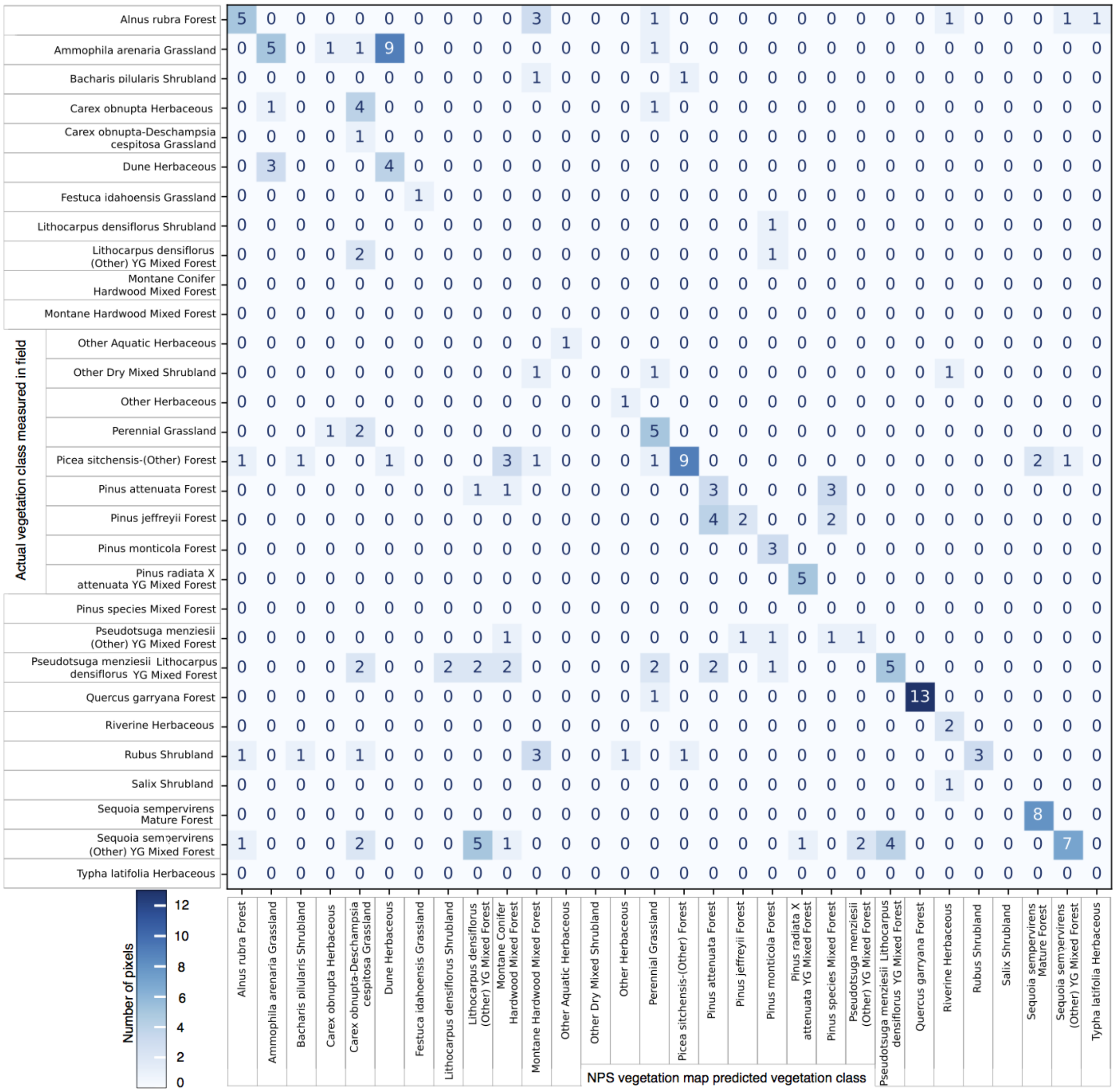


**Fig. S27 | Confusion matrix of official NPS vegetation map on held-out field plots**

Confusion matrix of the predicted vegetation type from the official and proprietary NPS vegetation map using held-out field plots from the fold with the highest single-label accuracy by the *deepbiosphere*-based classifier. The single-label accuracy of the NPS vegetation map was 44.44% for the 189 held-out test field plots.

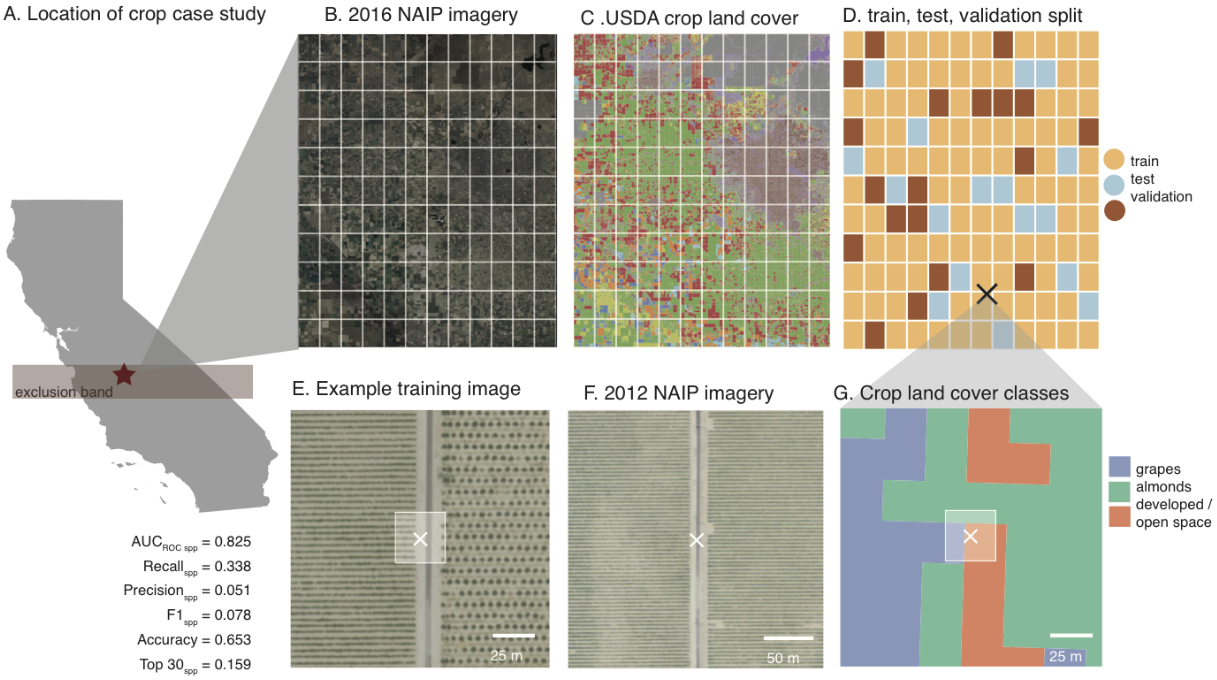


**Fig. S28 | USDA crop-specific land cover classification case study.**
**(A)** Location of Central Valley case study from ref. [67]. The red band signifies the portion of California excluded from training *deepbiosphere*$_5$. Included are a selection of accuracy metrics for *deepbiosphere*$_5$ on observations from the held out region. **(B)** 2016 NAIP imagery of 60cm resolution for the case study with dataset split blocks overlaid. **(C)** USDA cropland data layer. Only 69 crop types are present in the study area. Crop types are binned into ten categories and colored randomly (key in **Fig. 29**). **(D)** Partitioning of case study into train, test, and validation blocks from ref. [67]. Approximately 73% of the blocks are for training and 13% are for testing. The example location is denoted by a black X. **(E)** Example image from the 2016 NAIP imagery used to test both deepbiosphere and *TileNet*[67]. *Deepbiosphere* uses 256 x 256 pixel images to make predictions while TileNet (white box) uses 50 x 50 pixel images. The per-pixel resolution is 60 cm for this image. *TileNet* was trained using this specific year of imagery, while *deepbiosphere*$_5$ was trained with 1 m resolution NAIP imagery from 2012. **(F)** Example image for the same location but from the 2012 NAIP acquisition, the imagery *deepbiosphere*$_5$ was trained with. The resolution is 1 m per-pixel instead of 60 cm, leading to a much different image, and in some cases different crops planted in the field. This means that the 2016 NAIP imagery used in this case study is out of distribution for *deepbiosphere*$_5$, making it a harder prediction task for the model. **(G)** The crop land cover classes for the given example training image. For each example, the class with the largest area is used as the ground truth label for that image, as done in ref. [67]. Since the size of the images used to train *deepbiosphere*$_5$ and *TileNet* are quite different, the maximum area crop land cover class is separately calculated for both *deepbiosphere*$_5$ and *TileNet* respectively. Occasionally the ground truth labels will differ, as in this example where almonds is the class with largest area for the *deepbiosphere*-sized image while open space is the largest area class for *TileNet*, but this approach provides the fairest comparison for each respective model.

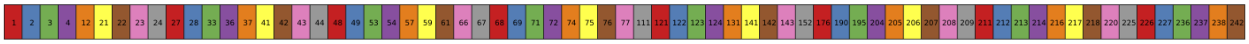


**Fig. S29 | Color key for cropland datalayer map**
Number values map to cropland classes from USDA cropland data layer[68].

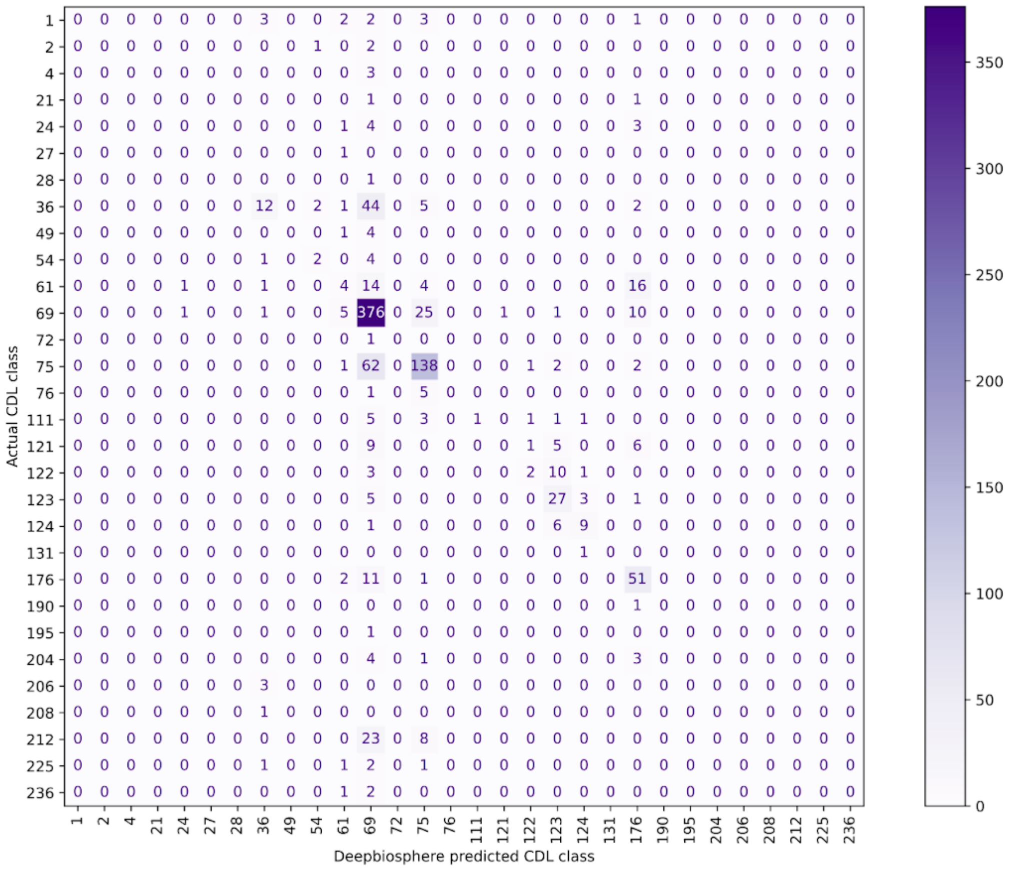


**Fig. S30 | Confusion matrix of *deepbiosphere*-based crop type predictions to USDA cropland data layer**
Confusion matrix of the predictions of the random forest feature extractor fitted using *deepbiosphere*$_5$ features with the highest single-label accuracy fold from the ten-fold cross-validation trials. The single-label accuracy of the classifier was 62.2% comparing the maximum class type of the official USDA cropland data layer map[68] as ground truth for each of the unseen 1,000 test examples. Some classes like grapes (class 69) are well predicted while others like oranges (212) are not.

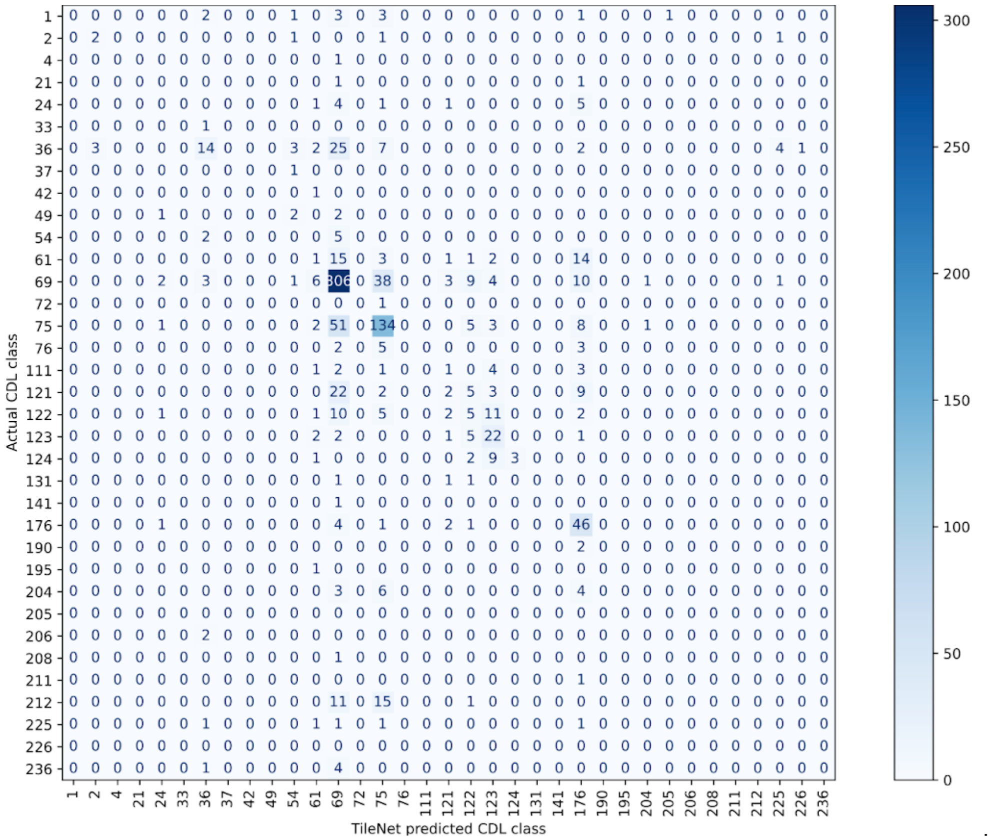


**Fig. S31 | Confusion matrix of *TileNet*-based crop type predictions to USDA cropland data layer**

Confusion matrix of the predictions of a random forest feature extractor fitted using *TileNet*[67]. The single-label accuracy of the classifier was 53.5% comparing the maximum class type of the official USDA cropland data layer map[68] as ground truth for each of the unseen 1,000 test examples. The classifier predicts some classes well like grapes (class 69), while it fails with others like developed/open space (121).

#### SUPPLEMENTAL TABLES

| | | |
|---|---|---|
| Plant biodiversity dataset | Observation date range | 1/1/2015 - 5/1/2022 |
| | Number of unique vascular species | 2,221 |
| | Total plant diversity present | 29.216% |
| | Number of unique genera | 878 |
| | Number of unique families | 153 |
| | Number of unique observations | 652,027 images |
| | Number of linked observations | 614,727 images |
| | Threshold for species inclusion | 500 linked observations |
| | Shannon diversity index | 6.825 |
| | Gini inequality index | 0.334 |
| | Simpson index | 1.000 |
| NAIP Aerial Imagery | Spatial resolution of imagery | 1 meter ground sample distance |
| | Bands used | Blue (428-492 nm) |
| | | Green (533-587 nm) |
| | | Red (608-662 nm) |
| | | Near-Infrared (883-887 nm) |
| | Year of observation | 2012 |
| | Number of unique images | 11,095 |

**Table S1 | Key metrics of dataset.**
Relevant metrics for biodiversity dataset, including number of observations, images, and summary statistics for observations. Included is also information about the National Agriculture Imagery Program aerial imagery used in model training.

| Layer (type:depth-idx) | Output Shape | # Params | Layers Cont'd | Output Shape Cont'd | # Params Cont'd |
|---|---|---|---|---|---|
| ├─Sequential: 1-1 | [65, 2048, 8, 8] | -- | │ │ └─Bottleneck: 3-13 | [65, 1024, 16, 16] | 1,183,104 |
| │ └─SpaceToDepthModule: 2-1 | [65, 64, 64, 64] | -- | │ │ └─Bottleneck: 3-14 | [65, 1024, 16, 16] | 1,183,104 |
| │ └─Sequential: 2-2 | [65, 64, 64, 64] | -- | │ │ └─Bottleneck: 3-15 | [65, 1024, 16, 16] | 1,183,104 |
| │ │ └─Conv2d: 3-1 | [65, 64, 64, 64] | 36,864 | │ │ └─Bottleneck: 3-16 | [65, 1024, 16, 16] | 1,183,104 |
| │ │ └─InPlaceABN: 3-2 | [65, 64, 64, 64] | 128 | │ │ └─Bottleneck: 3-17 | [65, 1024, 16, 16] | 1,183,104 |
| │ └─Sequential: 2-3 | [65, 64, 64, 64] | -- | │ │ └─Bottleneck: 3-18 | [65, 1024, 16, 16] | 1,183,104 |
| │ │ └─BasicBlock: 3-3 | [65, 64, 64, 64] | 82,304 | │ │ └─Bottleneck: 3-19 | [65, 1024, 16, 16] | 1,183,104 |
| │ │ └─BasicBlock: 3-4 | [65, 64, 64, 64] | 82,304 | │ │ └─Bottleneck: 3-20 | [65, 1024, 16, 16] | 1,183,104 |
| │ │ └─BasicBlock: 3-5 | [65, 64, 64, 64] | 82,304 | │ └─Sequential: 2-6 | [65, 2048, 8, 8] | -- |
| │ └─Sequential: 2-4 | [65, 128, 32, 32] | -- | │ │ └─Bottleneck: 3-21 | [65, 2048, 8, 8] | 6,039,552 |
| │ │ └─BasicBlock: 3-6 | [65, 128, 32, 32] | 246,720 | │ │ └─Bottleneck: 3-22 | [65, 2048, 8, 8] | 4,462,592 |
| │ │ └─BasicBlock: 3-7 | [65, 128, 32, 32] | 312,000 | │ │ └─Bottleneck: 3-23 | [65, 2048, 8, 8] | 4,462,592 |
| │ │ └─BasicBlock: 3-8 | [65, 128, 32, 32] | 312,000 | ├─Sequential: 1-2 | [65, 2048] | -- |
| │ │ └─BasicBlock: 3-9 | [65, 128, 32, 32] | 312,000 | │ └─FastAvgPool2d: 2-7 | [65, 2048] | -- |
| │ └─Sequential: 2-5 | [65, 1024, 16, 16] | -- | ├─Linear: 1-3 | [65, 2221] | 4,550,829 |
| │ │ └─Bottleneck: 3-10 | [65, 1024, 16, 16] | 1,086,848 | ├─Linear: 1-4 | [65, 878] | 1,799,022 |
| │ │ └─Bottleneck: 3-11 | [65, 1024, 16, 16] | 1,183,104 | ├─Linear: 1-5 | [65, 153] | 313,497 |
| │ │ └─Bottleneck: 3-12 | [65, 1024, 16, 16] | 1,183,104 | | | |
| Total params: 36,012,596 | | | | | |
| Batch size: 65 images | | | | | |
| Input size (MB): 68.16 | | | | | |
| Forward/backward pass size (MB): 9,682.84 | | | | | |
| Params size (MB): 144.05 | | | | | |
| Estimated Total Size (MB): 9895.05 | | | | | |
| GPU: NVIDIA GRID M60-8Q | | | | | |

**Table S2 | Model summary of *TResNet* architecture**
Summary of training statistics and parameters of the remote sensing-only CNN architecture. The right-hand side columns are the continuation of the model summary. Summary generated using torchinfo version 1.7.0[72].

| Layer (type:depth-idx) | Output Shape | # Params | Layers Cont'd | Output Shape Cont'd | # Params Cont'd |
|---|---|---|---|---|---|
| ├─Sequential: 1-1 | [150, 2048, 8, 8] | -- | │ │ └─Bottleneck: 3-22 | [150, 2048, 8, 8] | 4,462,592 |
| │ └─SpaceToDepthModule: 2-1 | [150, 64, 64, 64] | -- | │ │ └─Bottleneck: 3-23 | [150, 2048, 8, 8] | 4,462,592 |
| │ └─Sequential: 2-2 | [150, 64, 64, 64] | -- | ├─Sequential: 1-2 | [150, 2048] | -- |
| │ │ └─Conv2d: 3-1 | [150, 64, 64, 64] | 36,864 | │ └─FastAvgPool2d: 2-7 | [150, 2048] | -- |
| │ │ └─InPlaceABN: 3-2 | [150, 64, 64, 64] | 128 | ├─Sequential: 1-3 | [150, 2048] | -- |
| │ └─Sequential: 2-3 | [150, 64, 64, 64] | -- | │ └─Linear: 2-8 | [150, 2048] | 4,196,352 |
| │ │ └─BasicBlock: 3-3 | [150, 64, 64, 64] | 82,304 | │ └─BatchNorm1d: 2-9 | [150, 2048] | 4,096 |
| │ │ └─BasicBlock: 3-4 | [150, 64, 64, 64] | 82,304 | │ └─ReLU: 2-10 | [150, 2048] | -- |
| │ │ └─BasicBlock: 3-5 | [150, 64, 64, 64] | 82,304 | ├─Sequential: 1-4 | [150, 2048] | -- |
| │ └─Sequential: 2-4 | [150, 128, 32, 32] | -- | │ └─Linear: 2-11 | [150, 1000] | 20,000 |
| │ │ └─BasicBlock: 3-6 | [150, 128, 32, 32] | 246,720 | │ └─ELU: 2-12 | [150, 1000] | -- |
| │ │ └─BasicBlock: 3-7 | [150, 128, 32, 32] | 312,000 | │ └─Linear: 2-13 | [150, 1000] | 1,001,000 |
| │ │ └─BasicBlock: 3-8 | [150, 128, 32, 32] | 312,000 | │ └─ELU: 2-14 | [150, 1000] | -- |
| │ │ └─BasicBlock: 3-9 | [150, 128, 32, 32] | 312,000 | │ └─Linear: 2-15 | [150, 2000] | 2,002,000 |
| │ └─Sequential: 2-5 | [150, 1024, 16, 16] | -- | │ └─ELU: 2-16 | [150, 2000] | -- |
| │ │ └─Bottleneck: 3-10 | [150, 1024, 16, 16] | 1,086,848 | │ └─Dropout: 2-17 | [150, 2000] | -- |
| │ │ └─Bottleneck: 3-11 | [150, 1024, 16, 16] | 1,183,104 | │ └─Linear: 2-18 | [150, 2000] | 4,002,000 |
| │ │ └─Bottleneck: 3-12 | [150, 1024, 16, 16] | 1,183,104 | │ └─ELU: 2-19 | [150, 2000] | -- |
| │ │ └─Bottleneck: 3-13 | [150, 1024, 16, 16] | 1,183,104 | │ └─Linear: 2-20 | [150, 2048] | 4,098,048 |
| │ │ └─Bottleneck: 3-14 | [150, 1024, 16, 16] | 1,183,104 | │ └─BatchNorm1d: 2-21 | [150, 2048] | 4,096 |
| │ │ └─Bottleneck: 3-15 | [150, 1024, 16, 16] | 1,183,104 | │ └─ELU: 2-22 | [150, 2048] | -- |
| │ │ └─Bottleneck: 3-16 | [150, 1024, 16, 16] | 1,183,104 | ├─Sequential: 1-5 | [150, 2048] | -- |
| │ │ └─Bottleneck: 3-17 | [150, 1024, 16, 16] | 1,183,104 | │ └─Linear: 2-23 | [150, 2048] | 8,390,656 |
| │ │ └─Bottleneck: 3-18 | [150, 1024, 16, 16] | 1,183,104 | │ └─ReLU: 2-24 | [150, 2048] | -- |
| │ │ └─Bottleneck: 3-19 | [150, 1024, 16, 16] | 1,183,104 | ├─Linear: 1-6 | [150, 2221] | 4,550,829 |
| │ │ └─Bottleneck: 3-20 | [150, 1024, 16, 16] | 1,183,104 | ├─Linear: 1-7 | [150, 878] | 1,799,022 |
| │ └─Sequential: 2-6 | [150, 2048, 8, 8] | -- | ├─Linear: 1-8 | [150, 153] | 313,497 |
| │ │ └─Bottleneck: 3-21 | [150, 2048, 8, 8] | 6,039,552 | | | |
| Total params: 59,730,844 | | | | | |
| Batch size: 150 images | | | | | |
| Input size (MB): 157.30 | | | | | |
| Forward/backward pass size (MB): 22,364.51 | | | | | |
| Params size (MB): 238.92 | | | | | |
| Estimated Total Size (MB): 22,760.73 | | | | | |
| GPU: NVIDIA A100 | | | | | |

**Table S3 | Model summary of *Joint TResNet* architecture**
Summary of training statistics and parameters of the *deepbiosphere* architecture. The right-hand side columns are the continuation of the model summary. Summary generated using torchinfo version 1.7.0[72].

| Layer (type:depth-idx) | Output Shape | # Params | Layers Cont'd | Output Shape Cont'd | # Params Cont'd |
|---|---|---|---|---|---|
| ├─BasicConv2d: 1-1 | [100, 32, 254, 254] | -- | │ │ └─Conv2d: 3-87 | [100, 192, 30, 30] | 215,040 |
| │ └─Conv2d: 2-1 | [100, 32, 254, 254] | 1,152 | │ │ └─BatchNorm2d: 3-88 | [100, 192, 30, 30] | 384 |
| │ └─BatchNorm2d: 2-2 | [100, 32, 254, 254] | 64 | │ └─BasicConv2d: 2-55 | [100, 192, 30, 30] | -- |
| ├─BasicConv2d: 1-2 | [100, 32, 252, 252] | -- | │ │ └─Conv2d: 3-89 | [100, 192, 30, 30] | 147,456 |
| │ └─Conv2d: 2-3 | [100, 32, 252, 252] | 9,216 | │ │ └─BatchNorm2d: 3-90 | [100, 192, 30, 30] | 384 |
| │ └─BatchNorm2d: 2-4 | [100, 32, 252, 252] | 64 | ├─InceptionC: 1-14 | [100, 768, 30, 30] | -- |
| ├─BasicConv2d: 1-3 | [100, 64, 252, 252] | -- | │ └─BasicConv2d: 2-56 | [100, 192, 30, 30] | -- |
| │ └─Conv2d: 2-5 | [100, 64, 252, 252] | 18,432 | │ │ └─Conv2d: 3-91 | [100, 192, 30, 30] | 147,456 |
| │ └─BatchNorm2d: 2-6 | [100, 64, 252, 252] | 128 | │ │ └─BatchNorm2d: 3-92 | [100, 192, 30, 30] | 384 |
| ├─MaxPool2d: 1-4 | [100, 64, 125, 125] | -- | │ └─BasicConv2d: 2-57 | [100, 160, 30, 30] | -- |
| ├─BasicConv2d: 1-5 | [100, 80, 125, 125] | -- | │ │ └─Conv2d: 3-93 | [100, 160, 30, 30] | 122,880 |
| │ └─Conv2d: 2-7 | [100, 80, 125, 125] | 5,120 | │ │ └─BatchNorm2d: 3-94 | [100, 160, 30, 30] | 320 |
| │ └─BatchNorm2d: 2-8 | [100, 80, 125, 125] | 160 | │ └─BasicConv2d: 2-58 | [100, 160, 30, 30] | -- |
| ├─BasicConv2d: 1-6 | [100, 192, 123, 123] | -- | │ │ └─Conv2d: 3-95 | [100, 160, 30, 30] | 179,200 |
| │ └─Conv2d: 2-9 | [100, 192, 123, 123] | 138,240 | │ │ └─BatchNorm2d: 3-96 | [100, 160, 30, 30] | 320 |
| │ └─BatchNorm2d: 2-10 | [100, 192, 123, 123] | 384 | │ └─BasicConv2d: 2-59 | [100, 192, 30, 30] | -- |
| ├─MaxPool2d: 1-7 | [100, 192, 61, 61] | -- | │ │ └─Conv2d: 3-97 | [100, 192, 30, 30] | 215,040 |
| ├─InceptionA: 1-8 | [100, 256, 61, 61] | -- | │ │ └─BatchNorm2d: 3-98 | [100, 192, 30, 30] | 384 |
| │ └─BasicConv2d: 2-11 | [100, 64, 61, 61] | -- | │ └─BasicConv2d: 2-60 | [100, 160, 30, 30] | -- |
| │ │ └─Conv2d: 3-1 | [100, 64, 61, 61] | 12,288 | │ │ └─Conv2d: 3-99 | [100, 160, 30, 30] | 122,880 |
| │ │ └─BatchNorm2d: 3-2 | [100, 64, 61, 61] | 128 | │ │ └─BatchNorm2d: 3-100 | [100, 160, 30, 30] | 320 |
| │ └─BasicConv2d: 2-12 | [100, 48, 61, 61] | -- | │ └─BasicConv2d: 2-61 | [100, 160, 30, 30] | -- |
| │ │ └─Conv2d: 3-3 | [100, 48, 61, 61] | 9,216 | │ │ └─Conv2d: 3-101 | [100, 160, 30, 30] | 179,200 |
| │ │ └─BatchNorm2d: 3-4 | [100, 48, 61, 61] | 96 | │ │ └─BatchNorm2d: 3-102 | [100, 160, 30, 30] | 320 |
| │ └─BasicConv2d: 2-13 | [100, 64, 61, 61] | -- | │ └─BasicConv2d: 2-62 | [100, 160, 30, 30] | -- |
| │ │ └─Conv2d: 3-5 | [100, 64, 61, 61] | 76,800 | │ │ └─Conv2d: 3-103 | [100, 160, 30, 30] | 179,200 |
| │ │ └─BatchNorm2d: 3-6 | [100, 64, 61, 61] | 128 | │ │ └─BatchNorm2d: 3-104 | [100, 160, 30, 30] | 320 |
| │ └─BasicConv2d: 2-14 | [100, 64, 61, 61] | -- | │ └─BasicConv2d: 2-63 | [100, 160, 30, 30] | -- |
| │ │ └─Conv2d: 3-7 | [100, 64, 61, 61] | 12,288 | │ │ └─Conv2d: 3-105 | [100, 160, 30, 30] | 179,200 |
| │ │ └─BatchNorm2d: 3-8 | [100, 64, 61, 61] | 128 | │ │ └─BatchNorm2d: 3-106 | [100, 160, 30, 30] | 320 |
| │ └─BasicConv2d: 2-15 | [100, 96, 61, 61] | -- | │ └─BasicConv2d: 2-64 | [100, 192, 30, 30] | -- |
| │ │ └─Conv2d: 3-9 | [100, 96, 61, 61] | 55,296 | │ │ └─Conv2d: 3-107 | [100, 192, 30, 30] | 215,040 |
| │ │ └─BatchNorm2d: 3-10 | [100, 96, 61, 61] | 192 | │ │ └─BatchNorm2d: 3-108 | [100, 192, 30, 30] | 384 |
| │ └─BasicConv2d: 2-16 | [100, 96, 61, 61] | -- | │ └─BasicConv2d: 2-65 | [100, 192, 30, 30] | -- |
| │ │ └─Conv2d: 3-11 | [100, 96, 61, 61] | 82,944 | │ │ └─Conv2d: 3-109 | [100, 192, 30, 30] | 147,456 |
| │ │ └─BatchNorm2d: 3-12 | [100, 96, 61, 61] | 192 | │ │ └─BatchNorm2d: 3-110 | [100, 192, 30, 30] | 384 |
| │ └─BasicConv2d: 2-17 | [100, 32, 61, 61] | -- | ├─InceptionC: 1-15 | [100, 768, 30, 30] | -- |
| │ │ └─Conv2d: 3-13 | [100, 32, 61, 61] | 6,144 | │ └─BasicConv2d: 2-66 | [100, 192, 30, 30] | -- |
| │ │ └─BatchNorm2d: 3-14 | [100, 32, 61, 61] | 64 | │ │ └─Conv2d: 3-111 | [100, 192, 30, 30] | 147,456 |
| ├─InceptionA: 1-9 | [100, 288, 61, 61] | -- | │ │ └─BatchNorm2d: 3-112 | [100, 192, 30, 30] | 384 |
| │ └─BasicConv2d: 2-18 | [100, 64, 61, 61] | -- | │ └─BasicConv2d: 2-67 | [100, 192, 30, 30] | -- |
| │ │ └─Conv2d: 3-15 | [100, 64, 61, 61] | 16,384 | │ │ └─Conv2d: 3-113 | [100, 192, 30, 30] | 147,456 |
| │ │ └─BatchNorm2d: 3-16 | [100, 64, 61, 61] | 128 | │ │ └─BatchNorm2d: 3-114 | [100, 192, 30, 30] | 384 |
| │ └─BasicConv2d: 2-19 | [100, 48, 61, 61] | -- | │ └─BasicConv2d: 2-68 | [100, 192, 30, 30] | -- |
| │ │ └─Conv2d: 3-17 | [100, 48, 61, 61] | 12,288 | │ │ └─Conv2d: 3-115 | [100, 192, 30, 30] | 258,048 |
| │ │ └─BatchNorm2d: 3-18 | [100, 48, 61, 61] | 96 | │ │ └─BatchNorm2d: 3-116 | [100, 192, 30, 30] | 384 |
| │ └─BasicConv2d: 2-20 | [100, 64, 61, 61] | -- | │ └─BasicConv2d: 2-69 | [100, 192, 30, 30] | -- |
| │ │ └─Conv2d: 3-19 | [100, 64, 61, 61] | 76,800 | │ │ └─Conv2d: 3-117 | [100, 192, 30, 30] | 258,048 |
| │ │ └─BatchNorm2d: 3-20 | [100, 64, 61, 61] | 128 | │ │ └─BatchNorm2d: 3-118 | [100, 192, 30, 30] | 384 |

| | | |
|---|---|---|
| │ └─BasicConv2d: 2-21 | [100, 64, 61, 61] | -- |
| │ │ └─Conv2d: 3-21 | [100, 64, 61, 61] | 16,384 |
| │ │ └─BatchNorm2d: 3-22 | [100, 64, 61, 61] | 128 |
| │ └─BasicConv2d: 2-22 | [100, 96, 61, 61] | -- |
| │ │ └─Conv2d: 3-23 | [100, 96, 61, 61] | 55,296 |
| │ │ └─BatchNorm2d: 3-24 | [100, 96, 61, 61] | 192 |
| │ └─BasicConv2d: 2-23 | [100, 96, 61, 61] | -- |
| │ │ └─Conv2d: 3-25 | [100, 96, 61, 61] | 82,944 |
| │ │ └─BatchNorm2d: 3-26 | [100, 96, 61, 61] | 192 |
| │ └─BasicConv2d: 2-24 | [100, 64, 61, 61] | -- |
| │ │ └─Conv2d: 3-27 | [100, 64, 61, 61] | 16,384 |
| │ │ └─BatchNorm2d: 3-28 | [100, 64, 61, 61] | 128 |
| ├─InceptionA: 1-10 | [100, 288, 61, 61] | -- |
| │ └─BasicConv2d: 2-25 | [100, 64, 61, 61] | -- |
| │ │ └─Conv2d: 3-29 | [100, 64, 61, 61] | 18,432 |
| │ │ └─BatchNorm2d: 3-30 | [100, 64, 61, 61] | 128 |
| │ └─BasicConv2d: 2-26 | [100, 48, 61, 61] | -- |
| │ │ └─Conv2d: 3-31 | [100, 48, 61, 61] | 13,824 |
| │ │ └─BatchNorm2d: 3-32 | [100, 48, 61, 61] | 96 |
| │ └─BasicConv2d: 2-27 | [100, 64, 61, 61] | -- |
| │ │ └─Conv2d: 3-33 | [100, 64, 61, 61] | 76,800 |
| │ │ └─BatchNorm2d: 3-34 | [100, 64, 61, 61] | 128 |
| │ └─BasicConv2d: 2-28 | [100, 64, 61, 61] | -- |
| │ │ └─Conv2d: 3-35 | [100, 64, 61, 61] | 18,432 |
| │ │ └─BatchNorm2d: 3-36 | [100, 64, 61, 61] | 128 |
| │ └─BasicConv2d: 2-29 | [100, 96, 61, 61] | -- |
| │ │ └─Conv2d: 3-37 | [100, 96, 61, 61] | 55,296 |
| │ │ └─BatchNorm2d: 3-38 | [100, 96, 61, 61] | 192 |
| │ └─BasicConv2d: 2-30 | [100, 96, 61, 61] | -- |
| │ │ └─Conv2d: 3-39 | [100, 96, 61, 61] | 82,944 |
| │ │ └─BatchNorm2d: 3-40 | [100, 96, 61, 61] | 192 |
| │ └─BasicConv2d: 2-31 | [100, 64, 61, 61] | -- |
| │ │ └─Conv2d: 3-41 | [100, 64, 61, 61] | 18,432 |
| │ │ └─BatchNorm2d: 3-42 | [100, 64, 61, 61] | 128 |
| ├─InceptionB: 1-11 | [100, 768, 30, 30] | -- |
| │ └─BasicConv2d: 2-32 | [100, 384, 30, 30] | -- |
| │ │ └─Conv2d: 3-43 | [100, 384, 30, 30] | 995,328 |
| │ │ └─BatchNorm2d: 3-44 | [100, 384, 30, 30] | 768 |
| │ └─BasicConv2d: 2-33 | [100, 64, 61, 61] | -- |
| │ │ └─Conv2d: 3-45 | [100, 64, 61, 61] | 18,432 |
| │ │ └─BatchNorm2d: 3-46 | [100, 64, 61, 61] | 128 |
| │ └─BasicConv2d: 2-34 | [100, 96, 61, 61] | -- |
| │ │ └─Conv2d: 3-47 | [100, 96, 61, 61] | 55,296 |
| │ │ └─BatchNorm2d: 3-48 | [100, 96, 61, 61] | 192 |
| │ └─BasicConv2d: 2-35 | [100, 96, 30, 30] | -- |
| │ │ └─Conv2d: 3-49 | [100, 96, 30, 30] | 82,944 |
| │ │ └─BatchNorm2d: 3-50 | [100, 96, 30, 30] | 192 |
| ├─InceptionC: 1-12 | [100, 768, 30, 30] | -- |
| │ └─BasicConv2d: 2-36 | [100, 192, 30, 30] | -- |
| │ │ └─Conv2d: 3-51 | [100, 192, 30, 30] | 147,456 |
| │ └─BasicConv2d: 2-70 | [100, 192, 30, 30] | -- |
| │ │ └─Conv2d: 3-119 | [100, 192, 30, 30] | 147,456 |
| │ │ └─BatchNorm2d: 3-120 | [100, 192, 30, 30] | 384 |
| │ └─BasicConv2d: 2-71 | [100, 192, 30, 30] | -- |
| │ │ └─Conv2d: 3-121 | [100, 192, 30, 30] | 258,048 |
| │ │ └─BatchNorm2d: 3-122 | [100, 192, 30, 30] | 384 |
| │ └─BasicConv2d: 2-72 | [100, 192, 30, 30] | -- |
| │ │ └─Conv2d: 3-123 | [100, 192, 30, 30] | 258,048 |
| │ │ └─BatchNorm2d: 3-124 | [100, 192, 30, 30] | 384 |
| │ └─BasicConv2d: 2-73 | [100, 192, 30, 30] | -- |
| │ │ └─Conv2d: 3-125 | [100, 192, 30, 30] | 258,048 |
| │ │ └─BatchNorm2d: 3-126 | [100, 192, 30, 30] | 384 |
| │ └─BasicConv2d: 2-74 | [100, 192, 30, 30] | -- |
| │ │ └─Conv2d: 3-127 | [100, 192, 30, 30] | 258,048 |
| │ │ └─BatchNorm2d: 3-128 | [100, 192, 30, 30] | 384 |
| │ └─BasicConv2d: 2-75 | [100, 192, 30, 30] | -- |
| │ │ └─Conv2d: 3-129 | [100, 192, 30, 30] | 147,456 |
| │ │ └─BatchNorm2d: 3-130 | [100, 192, 30, 30] | 384 |
| ├─InceptionAux: 1-16 | -- | -- |
| │ └─BasicConv2d: 2-76 | -- | -- |
| │ │ └─Conv2d: 3-131 | -- | 98,304 |
| │ │ └─BatchNorm2d: 3-132 | -- | 256 |
| │ └─BasicConv2d: 2-77 | -- | -- |
| │ │ └─Conv2d: 3-133 | -- | 2,457,600 |
| │ │ └─BatchNorm2d: 3-134 | -- | 1,536 |
| │ └─Linear: 2-78 | -- | 1,707,949 |
| ├─InceptionD: 1-17 | [100, 1280, 14, 14] | -- |
| │ └─BasicConv2d: 2-79 | [100, 192, 30, 30] | -- |
| │ │ └─Conv2d: 3-135 | [100, 192, 30, 30] | 147,456 |
| │ │ └─BatchNorm2d: 3-136 | [100, 192, 30, 30] | 384 |
| │ └─BasicConv2d: 2-80 | [100, 320, 14, 14] | -- |
| │ │ └─Conv2d: 3-137 | [100, 320, 14, 14] | 552,960 |
| │ │ └─BatchNorm2d: 3-138 | [100, 320, 14, 14] | 640 |
| │ └─BasicConv2d: 2-81 | [100, 192, 30, 30] | -- |
| │ │ └─Conv2d: 3-139 | [100, 192, 30, 30] | 147,456 |
| │ │ └─BatchNorm2d: 3-140 | [100, 192, 30, 30] | 384 |
| │ └─BasicConv2d: 2-82 | [100, 192, 30, 30] | -- |
| │ │ └─Conv2d: 3-141 | [100, 192, 30, 30] | 258,048 |
| │ │ └─BatchNorm2d: 3-142 | [100, 192, 30, 30] | 384 |
| │ └─BasicConv2d: 2-83 | [100, 192, 30, 30] | -- |
| │ │ └─Conv2d: 3-143 | [100, 192, 30, 30] | 258,048 |
| │ │ └─BatchNorm2d: 3-144 | [100, 192, 30, 30] | 384 |
| │ └─BasicConv2d: 2-84 | [100, 192, 14, 14] | -- |
| │ │ └─Conv2d: 3-145 | [100, 192, 14, 14] | 331,776 |
| │ │ └─BatchNorm2d: 3-146 | [100, 192, 14, 14] | 384 |
| ├─InceptionE: 1-18 | [100, 2048, 14, 14] | -- |
| │ └─BasicConv2d: 2-85 | [100, 320, 14, 14] | -- |
| │ │ └─Conv2d: 3-147 | [100, 320, 14, 14] | 409,600 |
| │ │ └─BatchNorm2d: 3-148 | [100, 320, 14, 14] | 640 |
| │ └─BasicConv2d: 2-86 | [100, 384, 14, 14] | -- |

| | | |
|---|---|---|
| │ │ └─BatchNorm2d: 3-52 | [100, 192, 30, 30] | 384 |
| │ └─BasicConv2d: 2-37 | [100, 128, 30, 30] | -- |
| │ │ └─Conv2d: 3-53 | [100, 128, 30, 30] | 98,304 |
| │ │ └─BatchNorm2d: 3-54 | [100, 128, 30, 30] | 256 |
| │ └─BasicConv2d: 2-38 | [100, 128, 30, 30] | -- |
| │ │ └─Conv2d: 3-55 | [100, 128, 30, 30] | 114,688 |
| │ │ └─BatchNorm2d: 3-56 | [100, 128, 30, 30] | 256 |
| │ └─BasicConv2d: 2-39 | [100, 192, 30, 30] | -- |
| │ │ └─Conv2d: 3-57 | [100, 192, 30, 30] | 172,032 |
| │ │ └─BatchNorm2d: 3-58 | [100, 192, 30, 30] | 384 |
| │ └─BasicConv2d: 2-40 | [100, 128, 30, 30] | -- |
| │ │ └─Conv2d: 3-59 | [100, 128, 30, 30] | 98,304 |
| │ │ └─BatchNorm2d: 3-60 | [100, 128, 30, 30] | 256 |
| │ └─BasicConv2d: 2-41 | [100, 128, 30, 30] | -- |
| │ │ └─Conv2d: 3-61 | [100, 128, 30, 30] | 114,688 |
| │ │ └─BatchNorm2d: 3-62 | [100, 128, 30, 30] | 256 |
| │ └─BasicConv2d: 2-42 | [100, 128, 30, 30] | -- |
| │ │ └─Conv2d: 3-63 | [100, 128, 30, 30] | 114,688 |
| │ │ └─BatchNorm2d: 3-64 | [100, 128, 30, 30] | 256 |
| │ └─BasicConv2d: 2-43 | [100, 128, 30, 30] | -- |
| │ │ └─Conv2d: 3-65 | [100, 128, 30, 30] | 114,688 |
| │ │ └─BatchNorm2d: 3-66 | [100, 128, 30, 30] | 256 |
| │ └─BasicConv2d: 2-44 | [100, 192, 30, 30] | -- |
| │ │ └─Conv2d: 3-67 | [100, 192, 30, 30] | 172,032 |
| │ │ └─BatchNorm2d: 3-68 | [100, 192, 30, 30] | 384 |
| │ └─BasicConv2d: 2-45 | [100, 192, 30, 30] | -- |
| │ │ └─Conv2d: 3-69 | [100, 192, 30, 30] | 147,456 |
| │ │ └─BatchNorm2d: 3-70 | [100, 192, 30, 30] | 384 |
| ├─InceptionC: 1-13 | [100, 768, 30, 30] | -- |
| │ └─BasicConv2d: 2-46 | [100, 192, 30, 30] | -- |
| │ │ └─Conv2d: 3-71 | [100, 192, 30, 30] | 147,456 |
| │ │ └─BatchNorm2d: 3-72 | [100, 192, 30, 30] | 384 |
| │ └─BasicConv2d: 2-47 | [100, 160, 30, 30] | -- |
| │ │ └─Conv2d: 3-73 | [100, 160, 30, 30] | 122,880 |
| │ │ └─BatchNorm2d: 3-74 | [100, 160, 30, 30] | 320 |
| │ └─BasicConv2d: 2-48 | [100, 160, 30, 30] | -- |
| │ │ └─Conv2d: 3-75 | [100, 160, 30, 30] | 179,200 |
| │ │ └─BatchNorm2d: 3-76 | [100, 160, 30, 30] | 320 |
| │ └─BasicConv2d: 2-49 | [100, 192, 30, 30] | -- |
| │ │ └─Conv2d: 3-77 | [100, 192, 30, 30] | 215,040 |
| │ │ └─BatchNorm2d: 3-78 | [100, 192, 30, 30] | 384 |
| │ └─BasicConv2d: 2-50 | [100, 160, 30, 30] | -- |
| │ │ └─Conv2d: 3-79 | [100, 160, 30, 30] | 122,880 |
| │ │ └─BatchNorm2d: 3-80 | [100, 160, 30, 30] | 320 |
| │ └─BasicConv2d: 2-51 | [100, 160, 30, 30] | -- |
| │ │ └─Conv2d: 3-81 | [100, 160, 30, 30] | 179,200 |
| │ │ └─BatchNorm2d: 3-82 | [100, 160, 30, 30] | 320 |
| │ └─BasicConv2d: 2-52 | [100, 160, 30, 30] | -- |
| │ │ └─Conv2d: 3-83 | [100, 160, 30, 30] | 179,200 |
| │ │ └─BatchNorm2d: 3-84 | [100, 160, 30, 30] | 320 |
| │ │ └─Conv2d: 3-149 | [100, 384, 14, 14] | 491,520 |
| │ │ └─BatchNorm2d: 3-150 | [100, 384, 14, 14] | 768 |
| │ └─BasicConv2d: 2-87 | [100, 384, 14, 14] | -- |
| │ │ └─Conv2d: 3-151 | [100, 384, 14, 14] | 442,368 |
| │ │ └─BatchNorm2d: 3-152 | [100, 384, 14, 14] | 768 |
| │ └─BasicConv2d: 2-88 | [100, 384, 14, 14] | -- |
| │ │ └─Conv2d: 3-153 | [100, 384, 14, 14] | 442,368 |
| │ │ └─BatchNorm2d: 3-154 | [100, 384, 14, 14] | 768 |
| │ └─BasicConv2d: 2-89 | [100, 448, 14, 14] | -- |
| │ │ └─Conv2d: 3-155 | [100, 448, 14, 14] | 573,440 |
| │ │ └─BatchNorm2d: 3-156 | [100, 448, 14, 14] | 896 |
| │ └─BasicConv2d: 2-90 | [100, 384, 14, 14] | -- |
| │ │ └─Conv2d: 3-157 | [100, 384, 14, 14] | 1,548,288 |
| │ │ └─BatchNorm2d: 3-158 | [100, 384, 14, 14] | 768 |
| │ └─BasicConv2d: 2-91 | [100, 384, 14, 14] | -- |
| │ │ └─Conv2d: 3-159 | [100, 384, 14, 14] | 442,368 |
| │ │ └─BatchNorm2d: 3-160 | [100, 384, 14, 14] | 768 |
| │ └─BasicConv2d: 2-92 | [100, 384, 14, 14] | -- |
| │ │ └─Conv2d: 3-161 | [100, 384, 14, 14] | 442,368 |
| │ │ └─BatchNorm2d: 3-162 | [100, 384, 14, 14] | 768 |
| │ └─BasicConv2d: 2-93 | [100, 192, 14, 14] | -- |
| │ │ └─Conv2d: 3-163 | [100, 192, 14, 14] | 245,760 |
| │ │ └─BatchNorm2d: 3-164 | [100, 192, 14, 14] | 384 |
| ├─InceptionE: 1-19 | [100, 2048, 14, 14] | -- |
| │ └─BasicConv2d: 2-94 | [100, 320, 14, 14] | -- |
| │ │ └─Conv2d: 3-165 | [100, 320, 14, 14] | 655,360 |
| │ │ └─BatchNorm2d: 3-166 | [100, 320, 14, 14] | 640 |
| │ └─BasicConv2d: 2-95 | [100, 384, 14, 14] | -- |
| │ │ └─Conv2d: 3-167 | [100, 384, 14, 14] | 786,432 |
| │ │ └─BatchNorm2d: 3-168 | [100, 384, 14, 14] | 768 |
| │ └─BasicConv2d: 2-96 | [100, 384, 14, 14] | -- |
| │ │ └─Conv2d: 3-169 | [100, 384, 14, 14] | 442,368 |
| │ │ └─BatchNorm2d: 3-170 | [100, 384, 14, 14] | 768 |
| │ └─BasicConv2d: 2-97 | [100, 384, 14, 14] | -- |
| │ │ └─Conv2d: 3-171 | [100, 384, 14, 14] | 442,368 |
| │ │ └─BatchNorm2d: 3-172 | [100, 384, 14, 14] | 768 |
| │ └─BasicConv2d: 2-98 | [100, 448, 14, 14] | -- |
| │ │ └─Conv2d: 3-173 | [100, 448, 14, 14] | 917,504 |
| │ │ └─BatchNorm2d: 3-174 | [100, 448, 14, 14] | 896 |
| │ └─BasicConv2d: 2-99 | [100, 384, 14, 14] | -- |
| │ │ └─Conv2d: 3-175 | [100, 384, 14, 14] | 1,548,288 |
| │ │ └─BatchNorm2d: 3-176 | [100, 384, 14, 14] | 768 |
| │ └─BasicConv2d: 2-100 | [100, 384, 14, 14] | -- |
| │ │ └─Conv2d: 3-177 | [100, 384, 14, 14] | 442,368 |
| │ │ └─BatchNorm2d: 3-178 | [100, 384, 14, 14] | 768 |
| │ └─BasicConv2d: 2-101 | [100, 384, 14, 14] | -- |
| │ │ └─Conv2d: 3-179 | [100, 384, 14, 14] | 442,368 |
| │ │ └─BatchNorm2d: 3-180 | [100, 384, 14, 14] | 768 |
| │ └─BasicConv2d: 2-102 | [100, 192, 14, 14] | -- |
| │ │ └─Conv2d: 3-181 | [100, 192, 14, 14] | 393,216 |

| | | | | | |
|---|---|---|---|---|---|
| │ └─BasicConv2d: 2-53 | [100, 160, 30, 30] | -- | │ │ └─BatchNorm2d: 3-182 | [100, 192, 14, 14] | 384 |
| │ │ └─Conv2d: 3-85 | [100, 160, 30, 30] | 179,200 | ├─AdaptiveAvgPool2d: 1-20 | [100, 2048, 1, 1] | -- |
| │ │ └─BatchNorm2d: 3-86 | [100, 160, 30, 30] | 320 | ├─Dropout: 1-21 | [100, 2048, 1, 1] | -- |
| │ └─BasicConv2d: 2-54 | [100, 192, 30, 30] | -- | ├─Linear: 1-22 | [100, 2221] | 4,550,829 |
| Total params: 30,602,330 | | | | | |
| Batch size: 100 images | | | | | |
| Input size (MB): 104.86 | | | | | |
| Forward/backward pass size (MB): 43,286.00 | | | | | |
| Params size (MB): 122.41 | | | | | |
| Estimated Total Size (MB): 43,513.27 | | | | | |
| GPU: NVIDIA A100 | | | | | |

**Table S4 | Model summary of *Inception V3* architecture**
Summary of training statistics and parameters of the *Inception V3* architecture baseline from ref. [41]. The right-hand side columns are the continuation of the model summary. Summary generated using torchinfo version 1.7.0[72].

| Layer (type:depth-idx) | Output Shape | # Params |
|---|---|---|
| Bioclim_MLP | [1000, 2221] | -- |
| ├─Sequential: 1-1 | [1000, 2000] | -- |
| │ └─Linear: 2-1 | [1000, 1000] | 20,000 |
| │ └─ELU: 2-2 | [1000, 1000] | -- |
| │ └─Linear: 2-3 | [1000, 1000] | 1,001,000 |
| │ └─ELU: 2-4 | [1000, 1000] | -- |
| │ └─Linear: 2-5 | [1000, 2000] | 2,002,000 |
| │ └─ELU: 2-6 | [1000, 2000] | -- |
| │ └─Dropout: 2-7 | [1000, 2000] | -- |
| │ └─Linear: 2-8 | [1000, 2000] | 4,002,000 |
| │ └─ELU: 2-9 | [1000, 2000] | -- |
| ├─Linear: 1-2 | [1000, 153] | 306,153 |
| ├─Linear: 1-3 | [1000, 878] | 1,756,878 |
| ├─Linear: 1-4 | [1000, 2221] | 4,444,221 |
| Total params: 13,532,252 | | |
| Batch size: 1,000 | | |
| Input size (MB): 0.08 | | |
| Forward/backward pass size (MB): 74.02 | | |
| Params size (MB): 54.13 | | |
| Estimated Total Size (MB): 128.22 | | |
| GPU: NVIDIA GRID M60-8Q | | |

**Table S5 | Model summary of *bioclimate MLP* architecture**
Summary of training statistics and parameters of the bioclimate MLP architecture inspired by ref [52]. Summary generated using torchinfo version 1.7.0[72].

| model | loss | LR | BS | epoch | train time | precision per-species | precision per-img. | precision per-species | recall per-img. |
|---|---|---|---|---|---|---|---|---|---|
| *Joint TResNet* | ScaledBCE | 0.0001 | 150 | 19 | 33.9 hours | **0.0195 [0.0-0.0691]** | **0.0357 [0.0-0.087]** | 0.1429 [0.0-0.5] | 0.4 [0.0-0.8] |
| *TResNet* | ScaledBCE | 0.0001 | 65 | 19 | 24.5 hours | 0.017 [0.0-0.0515] | 0.025 [0.0125-0.0612] | **0.2609 [0.0-0.6667]** | **0.5439 [0.25-1.0]** |
| *TResNet* | CE | 0.0001 | 500 | 19 | 17.6 hours | 0.0 [0.0-0.0] | 0.0 [0.0-0.0] | 0.0 [0.0-0.0] | 0.0 [0.0-0.0] |
| *TResNet* | BCE | 0.0001 | 65 | 19 | 91.6 hours | 0.0 [0.0-0.0] | 0.0 [0.0-0.0] | 0.0 [0.0-0.0] | 0.0 [0.0-0.0] |
| *TResNet* | ASL | 0.0005 | 550 | 19 | 13.0 hours | 0.0 [0.0-0.037] | 0.0 [0.0-0.0] | 0.0 [0.0-0.029] | 0.0 [0.0-0.0] |
| *TResNet* | ASLScaled | 0.0005 | 500 | 19 | 20.3 hours | 0.0132 [0.0-0.0472] | 0.0238 [0.0-0.0588] | 0.15 [0.0-0.4915] | 0.4 [0.0-0.8] |
| **model cont'd** | **loss** | **LR** | **BS** | **epoch** | **train time** | **f1 per-species** | **f1 per-img.** | **mAP** | **single-label accuracy** |
| *Joint TResNet* | ScaledBCE | 0.0001 | 150 | 19 | 33.9 hours | **0.0342 [0.0-0.1157]** | **0.0645 [0.0-0.1429]** | **0.1493** | 0.4547 |
| *TResNet* | ScaledBCE | 0.0001 | 65 | 19 | 24.5 hours | 0.0312 [0.0-0.0909] | 0.0476 [0.0244-0.1096] | 0.1381 | **0.5672** |
| *TResNet* | CE | 0.0001 | 500 | 19 | 17.6 hours | 0.0 [0.0-0.0] | 0.0 [0.0-0.0] | 0.1473 | 0.0612 |
| *TResNet* | BCE | 0.0001 | 65 | 19 | 91.6 hours | 0.0 [0.0-0.0] | 0.0 [0.0-0.0] | 0.1053 | 0.0283 |
| *TResNet* | ASL | 0.0005 | 550 | 19 | 13.0 hours | 0.0 [0.0-0.0311] | 0.0 [0.0-0.0] | 0.0885 | 0.0788 |
| *TResNet* | ScaledASL | 0.0005 | 500 | 19 | 20.3 hours | 0.0243 [0.0-0.0833] | 0.0455 [0.0-0.1053] | 0.1243 | 0.4583 |
| **model cont'd** | **loss** | **LR** | **BS** | **epoch** | **train time** | **img. top 1** | **img. top 5** | **img. top 30** | **img. top 100** |
| *Joint TResNet* | ScaledBCE | 0.0001 | 150 | 19 | 33.9 hours | 0.0397 | 0.1403 | **0.4245** | **0.7086** |
| *TResNet* | ScaledBCE | 0.0001 | 65 | 19 | 24.5 hours | 0.0347 | 0.1246 | 0.4098 | 0.681 |
| *TResNet* | CE | 0.0001 | 500 | 19 | 17.6 hours | **0.0415** | **0.1405** | 0.4157 | 0.6937 |
| *TResNet* | BCE | 0.0001 | 65 | 19 | 91.6 hours | 0.0262 | 0.0932 | 0.3104 | 0.5636 |
| *TResNet* | ASL | 0.0005 | 550 | 19 | 13.0 hours | 0.0233 | 0.0806 | 0.2672 | 0.5061 |
| *TResNet* | ScaledASL | 0.0005 | 500 | 19 | 20.3 hours | 0.0332 | 0.1179 | 0.3608 | 0.6142 |
| **model cont'd** | **loss** | **LR** | **BS** | **epoch** | **train time** | **species top 1** | **species top 5** | **species top 30** | **species top 100** |
| *Joint TResNet* | ScaledBCE | 0.0001 | 150 | 19 | 33.9 hours | 0.0 [0.0-0.0] | **0.0 [0.0-0.1111]** | **0.1429 [0.0-0.5]** | 0.5 [0.0-0.8571] |
| *TResNet* | ScaledBCE | 0.0001 | 65 | 19 | 24.5 hours | 0.0 [0.0-0.0] | 0.0 [0.0-0.0741] | 0.125 [0.0-0.4545] | 0.5 [0.0-0.8125] |
| *TResNet* | CE | 0.0001 | 500 | 19 | 17.6 hours | 0.0 [0.0-0.0] | 0.0 [0.0-0.0714] | **0.1429 [0.0-0.5]** | **0.5385 [0.0-0.8333]** |
| *TResNet* | BCE | 0.0001 | 65 | 19 | 91.6 hours | 0.0 [0.0-0.0] | 0.0 [0.0-0.0] | 0.0 [0.0-0.3333] | 0.4 [0.0-0.6667] |
| *TResNet* | ASL | 0.0005 | 550 | 19 | 13.0 hours | 0.0 [0.0-0.0] | 0.0 [0.0-0.0] | 0.0 [0.0-0.2857] | 0.3333 [0.0-0.6] |
| *TResNet* | ScaledASL | 0.0005 | 500 | 19 | 20.3 hours | 0.0 [0.0-0.0] | 0.0 [0.0-0.0435] | 0.0667 [0.0-0.3846] | 0.4545 [0.0-0.7105] |
| **model cont'd** | **loss** | **LR** | **BS** | **epoch** | **train time** | **ROC AUC** | **PRC AUC** | **calibrated ROC AUC** | **calibrated PRC AUC** |
| *Joint TResNet* | ScaledBCE | 0.0001 | 150 | 19 | 33.9 hours | **0.9428 [0.8703-0.9776]** | **0.0323 [0.0083-0.1071]** | **0.006 [0.0003-0.0167]** | 0.5621 [0.4994-0.7146] |
| *TResNet* | ScaledBCE | 0.0001 | 65 | 19 | 24.5 hours | 0.9262 [0.8466-0.9697] | 0.0287 [0.0073-0.0909] | 0.004 [0.0003-0.0103] | **0.6024 [0.4991-0.7755]** |
| *TResNet* | CE | 0.0001 | 500 | 19 | 17.6 hours | 0.9067 [0.7962-0.9616] | 0.0237 [0.0058-0.0716] | 0.0009 [0.0003-0.0051] | 0.5 [0.5-0.5] |
| *TResNet* | BCE | 0.0001 | 65 | 19 | 91.6 hours | 0.8837 [0.7827-0.9454] | 0.0199 [0.0039-0.0674] | 0.0009 [0.0003-0.0051] | 0.5 [0.4999-0.5] |
| *TResNet* | ASL | 0.0005 | 550 | 19 | 13.0 hours | 0.8442 [0.7311-0.9245] | 0.0136 [0.0034-0.0504] | 0.001 [0.0003-0.0198] | 0.4999 [0.4995-0.511] |
| *TResNet* | ScaledASL | 0.0005 | 500 | 19 | 20.3 hours | 0.8895 [0.7858-0.9562] | 0.0212 [0.0049-0.0729] | 0.0047 [0.0004-0.0127] | 0.558 [0.4984-0.6988] |

**Table S6 | Comparing the performance of different loss functions on uniform data split**

Cumulatively, The Joint TResNet model (*deepbiosphere*) had the highest accuracy on 11/20 metrics, the remote sensing-only TResNet trained with the novel Scaled BCE loss had the highest accuracy on 3/20 metrics, and the remote sensing-only TResNet trained with the classic single label cross-entropy (CE) loss had the highest accuracy on 4/20 metrics When controlling for architecture and comparing strictly by loss, the model trained with the novel Scaled BCE loss had the highest accuracy on 12/20 accuracy metrics, while the model trained with CE loss had the highest accuracy on 8/20 metrics. While the CE loss model performed slightly better on the ranking metrics than the Scaled BCE model, it had very low performance on the binary accuracy metrics and middling performance on the discrimination metrics, a consequence of the single-label target of the loss, which is optimized for predicting only one species at at time. Training times vary considerably depending on whether the implementation was implemented by ourselves or using optimized PyTorch libraries[73]. Reported statistics are median [1st quartile - 3rd quartile] calculated at epoch 19 for all models. Abbreviations: LR = learning rate; BS = batch size; img. = image; cal. = calibrated; mAP = mean average precision; ROC = receiver operating characteristic curve; AUC = area under the curve; PRC = precision-recall curve; ScaledBCE = scaled binary cross-entropy loss; BCE = binary cross-entropy loss; CE = cross-entropy loss; ASL = asymmetric focal loss; ScaledASL = scaled asymmetric focal loss.

| model | loss | LR | BS | epoch | train time | precision per-species | precision per-img. | recall per-species | recall per-img. |
|---|---|---|---|---|---|---|---|---|---|
| *Joint TResNet* | ScaledBCE | 0.0001 | 150 | 4 | 8.4 hours | **0.0183 [0.0021-0.0492]** | 0.0177 [0.0088-0.0446] | **0.6538 [0.0741-0.9406]** | **1.0 [0.6364-1.0]** |
| *Bioclim MLP* | ScaledBCE | 0.0001 | 1,000 | 45 | 49.7 mins. | 0.0132 [0.0-0.0424] | **0.0179 [0.009-0.0462]** | 0.4364 [0.0-0.8699] | 0.75 [0.5-1.0] |
| *Inception V3* | CE | 0.01 | 100 | 10 | 14.1 hours | 0.0 [0.0-0.0] | 0.0 [0.0-0.0] | 0.0 [0.0-0.0] | 0.0 [0.0-0.0] |
| *Maxent* | N/A | N/A | N/A | N/A | 52.6 hours | 0.0048 [0.0-0.0323] | 0.0 [0.0-0.0238] | 0.1348 [0.0-0.566] | 0.0 [0.0-0.5] |
| *Random forest* | N/A | N/A | N/A | N/A | 6.7 hours | 0.0086 [0.0-0.04] | 0.0076 [0.0-0.0317] | 0.3684 [0.0-0.8182] | 0.2821 [0.0-0.875] |
| *random* | N/A | N/A | N/A | N/A | N/A | 0.0016 [0.0005-0.0052] | 0.0048 | 0.5 [0.4786-0.5222] | 0.5007 |
| *frequency* | N/A | N/A | N/A | N/A | N/A | 0.0 [0.0-0.0] | 0.0 [0.0-0.0] | 0.0 [0.0-0.0] | 0.0 [0.0-0.0] |

| model | loss | LR | BS | epoch | train time | F1 per-species | F1 per--img. | mAP | single-label accuracy |
|---|---|---|---|---|---|---|---|---|---|
| *Joint TResNet* | ScaledBCE | 0.0001 | 150 | 4 | 8.4 hours | **0.0354 [0.0043-0.0921]** | 0.0347 [0.0174-0.084] | **0.1611** | **0.7764** |
| *Bioclim MLP* | ScaledBCE | 0.0001 | 1,000 | 45 | 49.7 mins. | 0.0254 [0.0-0.0795] | **0.0351 [0.0179-0.087]** | 0.1146 | 0.684 |
| *Inception V3* | CE | 0.01 | 100 | 10 | 14.1 hours | 0.0 [0.0-0.0] | 0.0 [0.0-0.0] | 0.153 | 0.0049 |
| *Maxent* | N/A | N/A | N/A | | 52.6 hours | 0.0089 [0.0-0.059] | 0.0 [0.0-0.0426] | 0.0261 | 0.2761 |
| *Random forest* | N/A | N/A | N/A | | 6.7 hours | 0.0166 [0.0-0.0721] | 0.0152 [0.0-0.0588] | 0.0569 | 0.3943 |
| *random* | N/A | N/A | N/A | | N/A | 0.0031 [0.001-0.0103] | 0.0091 | 0.0065 | 0.5022 |
| *frequency* | N/A | N/A | N/A | | N/A | 0.0 [0.0-0.0] | 0.0 [0.0-0.0] | 0.0384 | 0.0656 |

| model | loss | LR | BS | epoch | train time | img. top 1 | img. top 5 | img. top 30 | img. top 100 |
|---|---|---|---|---|---|---|---|---|---|
| *Joint TResNet* | ScaledBCE | 0.0001 | 150 | 4 | 8.4 hours | 0.0413 | **0.1472** | **0.4594** | **0.7481** |
| *Bioclim MLP* | ScaledBCE | 0.0001 | 1,000 | 45 | 49.7 mins. | 0.0197 | 0.0946 | 0.3792 | 0.6801 |
| *Inception V3* | CE | 0.01 | 100 | 10 | 14.1 hours | **0.0448** | 0.1408 | 0.4224 | 0.7062 |
| *Maxent* | N/A | N/A | N/A | | 52.6 hours | 0.0004 | 0.0053 | 0.0815 | 0.291 |
| *Random forest* | N/A | N/A | N/A | | 6.7 hours | 0.0138 | 0.046 | 0.184 | 0.3709 |
| *random* | N/A | N/A | N/A | | N/A | 0.0005 | 0.0022 | 0.0133 | 0.0452 |
| *frequency* | N/A | N/A | N/A | | N/A | 0.0066 | 0.0274 | 0.0846 | 0.1952 |

| model | loss | LR | BS | epoch | train time | species top 1 | species top 5 | species top 30 | species top 100 |
|---|---|---|---|---|---|---|---|---|---|
| *Joint TResNet* | ScaledBCE | 0.0001 | 150 | 4 | 8.4 hours | 0.0 [0.0-0.0] | **0.0 [0.0-0.0909]** | **0.1667 [0.0-0.5]** | **0.6 [0.0-0.9091]** |
| *Bioclim MLP* | ScaledBCE | 0.0001 | 1,000 | 45 | 49.7 mins. | 0.0 [0.0-0.0] | 0.0 [0.0-0.0] | 0.1 [0.0-0.4167] | 0.5 [0.0-0.8] |
| *Inception V3* | CE | 0.01 | 100 | 10 | 14.1 hours | 0.0 [0.0-0.0] | 0.0 [0.0-0.0] | 0.0 [0.0-0.3571] | 0.5 [0.0-0.8667] |
| *Maxent* | N/A | N/A | N/A | | 52.6 hours | 0.0 [0.0-0.0] | 0.0 [0.0-0.0] | 0.0 [0.0-0.0] | 0.0417 [0.0-0.5] |
| *Random forest* | N/A | N/A | N/A | | 6.7 hours | 0.0 [0.0-0.0] | 0.0 [0.0-0.0] | 0.0 [0.0-0.25] | 0.2857 [0.0-0.6129] |
| *random* | N/A | N/A | N/A | | N/A | 0.0 [0.0-0.0] | 0.0 [0.0-0.0] | 0.0 [0.0-0.019] | 0.0412 [0.0133-0.0667] |
| *frequency* | N/A | N/A | N/A | | N/A | 0.0 [0.0-0.0] | 0.0 [0.0-0.0] | 0.0 [0.0-0.0] | 0.0 [0.0-0.0] |

| model | loss | LR | BS | epoch | train time | ROC AUC | PRC AUC | cal. ROC AUC | cal. PRC AUC |
|---|---|---|---|---|---|---|---|---|---|
| *Joint TResNet* | ScaledBCE | 0.0001 | 150 | 4 | 8.4 hours | **0.9458 [0.879-0.981]** | **0.036 [0.0093-0.1077]** | 0.0014 [0.0001-0.0045] | **0.7266 [0.5-0.8752]** |
| *Bioclim MLP* | ScaledBCE | 0.0001 | 1,000 | 45 | 49.7 mins. | 0.9199 [0.8017-0.971] | 0.0342 [0.0076-0.0914] | **0.0018 [0.0002-0.0054]** | 0.6473 [0.4975-0.8383] |
| *Inception V3* | CE | 0.01 | 100 | 10 | 14.1 hours | 0.9213 [0.835-0.9624] | 0.0195 [0.0061-0.055] | 0.0008 [0.0002-0.0026] | 0.5 [0.5-0.5] |
| *Maxent* | N/A | N/A | N/A | | 52.6 hours | 0.8825 [0.7754-0.9505] | 0.018 [0.0039-0.0725] | **0.0018 [0.0002-0.0083]** | 0.5399 [0.4905-0.7301] |
| *Random forest* | N/A | N/A | N/A | | 6.7 hours | 0.882 [0.7639-0.9515] | 0.0237 [0.0044-0.0925] | 0.0015 [0.0002-0.0072] | 0.6364 [0.4942-0.8356] |

| | | | | | | | | |
|---|---|---|---|---|---|---|---|---|
| *random* | N/A | N/A | N/A | N/A | 0.5005 [0.4883-0.5131] | 0.0024 [0.0011-0.0061] | 0.0008 [0.0002-0.0026] | 0.3751 [0.3697-0.3807] |
| *frequency* | N/A | N/A | N/A | N/A | 0.5 [0.5-0.5] | 0.0016 [0.0005-0.0052] | 0.0007 [0.0002-0.0023] | 0.5 [0.5-0.5] |

**Table S7 | Comparison of *deepbiosphere* to baseline SDMs and previous deep-learning-based approaches**
Cumulatively, *deepbiosphere* (Joint TResNet) had the highest accuracy on 17/20 metrics, the *bioclim MLP* on 3/20 metrics, *InceptionV3* on 1/20 metrics, and *Maxent* on 1/20 metrics. Train / fitting times are also provided. For deep learning-based models the train time was calculated to the epoch of evaluation. For stacked species distribution models, fitting times are aggregated across the 2,221 species assuming fitting was done in serial one species at a time utilizing one core. Reported statistics are median [1st quartile - 3rd quartile] calculated using the epoch of evaluation for the deep learning-based models (**Fig. S5, SM 3.2.6**). Abbreviations: LR = learning rate; BS = batch size; MLP = multilayer perceptron; img. = image; cal. = calibrated; mAP = mean average precision; ROC = receiver operating characteristic curve; AUC = area under the curve; PRC = precision-recall curve; ScaledBCE = scaled binary cross-entropy loss; CE = cross-entropy loss.

| model | loss | LR | BS | epoch | precision per-species | precision per-img. | recall per-species | recall per-img. |
|---|---|---|---|---|---|---|---|---|
| *Joint TResNet* | ScaledBCE | 0.0001 | 150 | 4 | **0.0521 [0.0499-0.0576]** | **0.1079 [0.0886-0.1342]** | **0.3403 [0.3189-0.3634]** | **0.6631 [0.609-0.6988]** |
| *Bioclim MLP* | ScaledBCE | 0.0001 | 1,000 | 45 | 0.0385 [0.0324-0.0421] | 0.0883 [0.0724-0.1203] | 0.2182 [0.1921-0.2378] | 0.4798 [0.4585-0.52] |
| *Maxent* | N/A | N/A | N/A | N/A | 0.036 [0.0305-0.0376] | 0.0422 [0.0403-0.0445] | 0.3263 [0.2639-0.4767] | 0.421 [0.3583-0.6409] |
| *Random forest* | N/A | N/A | N/A | N/A | 0.0358 [0.0331-0.0394] | 0.0382 [0.0361-0.0459] | 0.4452 [0.3382-0.5758] | 0.5085 [0.432-0.6856] |
| *Frequency* | N/A | N/A | N/A | N/A | 0.0017 [0.001-0.0019] | 0.1265 [0.0706-0.1499] | 0.0135 [0.0122-0.0153] | 0.0979 [0.0674-0.1082] |
| **model cont'd** | **loss** | **LR** | **BS** | **epoch** | **F1 per-species** | **F1 per-img.** | **mAP** | **single-label accuracy** |
| *Joint TResNet* | ScaledBCE | 0.0001 | 150 | 4 | **0.0793 [0.075-0.0835]** | **0.1652 [0.1371-0.2001]** | **0.1892 [0.1744-0.2125]** | **0.6844 [0.6367-0.7237]** |
| *Bioclim MLP* | ScaledBCE | 0.0001 | 1,000 | 45 | 0.0581 [0.0495-0.0599] | 0.1356 [0.1089-0.1666] | 0.1137 [0.1003-0.1299] | 0.4931 [0.4683-0.5302] |
| *Maxent* | N/A | N/A | N/A | | 0.0541 [0.0448-0.0607] | 0.0698 [0.0624-0.0754] | 0.0389 [0.03-0.0484] | 0.426 [0.3619-0.6438] |
| *Random forest* | N/A | N/A | N/A | | 0.0574 [0.0488-0.0602] | 0.068 [0.0578-0.0738] | 0.0542 [0.0458-0.062] | 0.5104 [0.4369-0.6868] |
| *Frequency* | N/A | N/A | N/A | | 0.0028 [0.0018-0.0032] | 0.0958 [0.055-0.1001] | 0.0953 [0.0568-0.1006] | 0.1029 [0.07-0.1157] |
| **model cont'd** | **loss** | **LR** | **BS** | **epoch** | **top 1 per-img..** | **top 5 per-img..** | **top 30 per-img..** | **top 100 per-img..** |
| *Joint TResNet* | ScaledBCE | 0.0001 | 150 | 4 | **0.0219 [0.0205-0.0245]** | **0.0927 [0.0855-0.098]** | **0.3525 [0.3333-0.359]** | **0.6545 [0.628-0.666]** |
| *Bioclim MLP* | ScaledBCE | 0.0001 | 1,000 | 45 | 0.0071 [0.0062-0.0081] | 0.0396 [0.035-0.0433] | 0.2076 [0.1889-0.224] | 0.4883 [0.4606-0.4982] |
| *Maxent* | N/A | N/A | N/A | | 0.0004 [0.0001-0.0018] | 0.0037 [0.0019-0.0073] | 0.0454 [0.0399-0.06] | 0.1859 [0.158-0.1941] |
| *Random forest* | N/A | N/A | N/A | | 0.0036 [0.0025-0.0047] | 0.0209 [0.0128-0.023] | 0.0971 [0.0746-0.1096] | 0.2232 [0.2009-0.2817] |
| *Frequency* | N/A | N/A | N/A | | 0.0091 [0.0049-0.0111] | 0.0359 [0.0296-0.0429] | 0.1329 [0.0994-0.1493] | 0.3007 [0.2159-0.312] |
| **model cont'd** | **loss** | **lLR** | **BS** | **epoch** | **top 1 per-species** | **top 5 per-species** | **top 30 per-species** | **top 100 per-species** |
| *Joint TResNet* | ScaledBCE | 0.0001 | 150 | 4 | **0.0079 [0.0072-0.0086]** | **0.0313 [0.0293-0.0364]** | **0.1395 [0.129-0.1597]** | **0.3377 [0.3056-0.3564]** |
| *Bioclim MLP* | ScaledBCE | 0.0001 | 1,000 | 45 | 0.0037 [0.0036-0.005] | 0.0184 [0.0177-0.0198] | 0.0855 [0.081-0.1097] | 0.2329 [0.2143-0.2678] |
| *Maxent* | N/A | N/A | N/A | | 0.0002 [0.0001-0.0016] | 0.0035 [0.0016-0.0068] | 0.0495 [0.0415-0.0538] | 0.1672 [0.1476-0.1808] |
| *Random forest* | N/A | N/A | N/A | | 0.0038 [0.0032-0.0042] | 0.0165 [0.016-0.0176] | 0.0855 [0.0811-0.0901] | 0.2223 [0.2093-0.2318] |
| *Frequency* | N/A | N/A | N/A | | 0.0007 [0.0006-0.0008] | 0.0037 [0.0032-0.004] | 0.0212 [0.0188-0.0232] | 0.0649 [0.0616-0.074] |
| **model cont'd** | **loss** | **LR** | **BS** | **epoch** | **ROC AUC** | **PRC AUC** | **calibrated ROC AUC** | **calibrated PRC AUC** |
| *Joint TResNet* | ScaledBCE | 0.0001 | 150 | 4 | **0.8197 [0.7858-0.8281]** | **0.0722 [0.0683-0.0768]** | **0.0141 [0.012-0.016]** | **0.6004 [0.5862-0.6099]** |
| *Bioclim MLP* | ScaledBCE | 0.0001 | 1,000 | 45 | 0.6873 [0.6822-0.7305] | 0.0543 [0.0492-0.0576] | 0.0129 [0.0117-0.016] | 0.5517 [0.5464-0.562] |
| *Maxent* | N/A | N/A | N/A | | 0.7138 [0.7051-0.7312] | 0.0525 [0.048-0.0597] | 0.0114 [0.0097-0.0122] | 0.5463 [0.521-0.552] |
| *Random forest* | N/A | N/A | N/A | | 0.7019 [0.6946-0.7137] | 0.053 [0.0499-0.0601] | 0.0102 [0.0081-0.0117] | 0.5511 [0.5223-0.5641] |
| *Frequency* | N/A | N/A | N/A | | 0.5 [0.5-0.5] | 0.015 [0.0132-0.0172] | 0.0069 [0.006-0.0081] | 0.4933 [0.4924-0.4939] |

**Table S8 | Comparison of *deepbiosphere* to baseline SDMs across spatial cross-validation bands.**
The epoch of evaluation for deep learning models was determined using the average epoch of highest performance on the uniform train / test split. For deep learning-based models the train time was calculated to the epoch of evaluation. For stacked species distribution models, fitting times are aggregated across the 2,221 species assuming fitting was done in serial one species at a time utilizing one core. Cumulatively, *deepbiosphere* had the highest accuracy on all 20/20 metrics. *Deepbiosphere* = Joint TResNet; MLP = multilayer perceptron; LR = learning rate; BS = batch size; ScaledBCE = scaled binary cross-entropy loss; img. = image; cal. = calibrated; mAP = mean average precision; ROC = receiver operating characteristic curve; AUC = area under the curve; PRC = precision-recall curve; ScaledBCE = scaled binary cross-entropy loss. Reported statistics are median [1st quartile - 3rd quartile].

| **Redwoods case study location:** (41.209, -124.01) | | **Oaks case study location:** (34.533, -120.17) | | |
|---|---|---|---|---|
| Redwoods example locations | Grove – Park | Oaks example locations | Species | Calflora ID |
| (40.3527, -123.9894) | Bull Creek Flats – Humboldt Redwoods | (36.108322, -120.561226) | *Quercus lobata* | po68973 |
| (41.7564, -124.1087) | Grove of Titans – Jedediah Smith Redwoods State Park | (36.151438, -120.770939) | *Quercus lobata* | po68984 |
| (40.6554, -124.0998) | Elk River Trail Grove – Headwaters Preserve | (36.635690, -121.242532) | *Quercus lobata* | po122185 |

**Table S9 | Site details for individual species case studies and human annotation experiments**
Two locations within both species' predicted range on Calscape (https://calscape.org) were selected as case studies. For the redwoods case study (left-hand side), example locations were chosen based on known remaining old-growth redwood groves and Tall Trees Grove in Redwoods National and State Parks was selected as the case study location. For the oaks case study (right-hand side), a list of candidate locations was generated from Calflora *Quercus lobata* occurrence records, with the ultimate case study site being selected from a region with multiple observed oaks from an undersampled region in the biodiversity prediction dataset. For each site, the most centered NAIP imagery tile was selected as the extent of the case study. Each tile is approximately 5 x 6 km in extent.

| species | *Sequoia sempervirens* - tested with 131 observations | | | | |
|---|---|---|---|---|---|
| model | % obs. seen | $ROC_{AUC\ spp}$ | precision | recall | top 5 accuracy |
| $Inception_{unif}$ | 99.24% | 0.79014 | 0 | 0 | **0.92366** |
| $deepbiosphere_{10}$ | 0.0% | **0.85698** | **0.34930** | **0.79567** | 0.00763 |
| $Maxent_{10}$ | 0.0% | 0.82582 | 0.056 | 0.01084 | 0 |
| $MLP_{10}$ | 0.0% | 0.34858 | 0 | 0 | 0 |
| $RF_{10}$ | 0.0% | 0.84368 | 0 | 0 | 0 |

**Species associated with mature redwood forest**

| species | *Oxalis oregana* - tested with 133 observations | | | | | *Prosartes smithii* - tested with 49 observations | | | | |
|---|---|---|---|---|---|---|---|---|---|---|
| model | % obs. seen | $ROC_{AUC\ spp}$ | precision | recall | top 5 accuracy | % obs. seen | $ROC_{AUC\ spp}$ | precision | recall | top 5 accuracy |
| $Inception_{unif}$ | 97.74% | 0.82532 | 0 | 0 | **0.85714** | 100.0% | 0.77792 | 0 | 0 | 0 |
| $deepbiosphere_{10}$ | 0.0% | **0.82862** | **0.39191** | **0.68631** | 0.07519 | 0.0% | **0.82417** | **0.25535** | **0.56270** | 0 |
| $Maxent_{10}$ | 0.0% | 0.78706 | 0.10087 | 0.03452 | 0 | 0.0% | 0.61294 | 0.232094 | 0.27438 | 0 |
| $MLP_{10}$ | 0.0% | 0.61862 | 0.20872 | 0.03988 | 0.02256 | 0.0% | 0.55632 | 0 | 0 | 0 |
| $RF_{10}$ | 0.0% | 0.77491 | 0.56 | 0.00833 | 0 | 0.0% | 0.62661 | 0.19765 | 0.39657 | 0 |

**Species associated with secondary growth redwood forest**

| species | *Rubus ursinus* - tested with 26 observations | | | | | *Iris douglasiana* - tested with 13 observations | | | | |
|---|---|---|---|---|---|---|---|---|---|---|
| model | % obs. seen | $ROC_{AUC\ spp}$ | precision | recall | top 5 accuracy | % obs. seen | $ROC_{AUC\ spp}$ | precision | recall | top 5 accuracy |
| $Inception_{unif}$ | 100.0% | 0.62901 | 0 | 0 | 0.03846 | 100.0% | 0.60923 | 0 | 0 | 0 |
| $deepbiosphere_{10}$ | 0.0% | **0.78660** | 0.11850 | **0.91849** | **0.26923** | 0.0% | 0.72671 | 0.05976 | 0.58108 | 0 |
| $Maxent_{10}$ | 0.0% | 0.48530 | 0.04115 | 0.01456 | 0 | 0.0% | 0.48364 | 0 | 0 | 0 |
| $MLP_{10}$ | 0.0% | 0.66889 | **0.15895** | 0.60990 | 0.07692 | 0.0% | **0.81435** | **0.06628** | **0.99662** | **0.61538** |
| $RF_{10}$ | 0.0% | 0.48570 | 0 | 0 | 0 | 0.0% | 0.54729 | 0 | 0 | 0 |

**Species associated both with mature redwood forest**

| species | *Viola sempervirens* - tested with 80 observations | | | | | *Notholithocarpus densiflorus* - tested with 58 observations | | | | |
|---|---|---|---|---|---|---|---|---|---|---|
| metric | % obs. seen | $ROC_{AUC\ spp}$ | precision | recall | top 5 accuracy | % obs. seen | $ROC_{AUC\ spp}$ | precision | recall | top 5 accuracy |
| $Inception_{unif}$ | 95.0% | 0.75628 | 0 | 0 | 0.05 | 96.55% | **0.74933** | 0 | 0 | **0.06897** |
| $deepbiosphere_{10}$ | 0.0% | **0.84209** | 0.31210 | **0.86478** | **0.15** | 0.0% | 0.71063 | 0.18954 | **0.45876** | 0.03448 |
| $Maxent_{10}$ | 0.0% | 0.67921 | **0.41838** | 0.52865 | 0 | 0.0% | 0.72215 | 0 | 0 | 0 |
| $MLP_{10}$ | 0.0% | 0.39984 | 0.07172 | 0.02674 | 0.0125 | 0.0% | 0.64161 | 0.17065 | 0.17740 | 0.01724 |
| $RF_{10}$ | 0.0% | 0.66408 | 0.24304 | 0.14668 | 0 | 0.0% | 0.77140 | **0.28070** | 0.01808 | 0 |

**Table S10 | Accuracy of various SDMs for predicting previously unseen occurrences in northern California**
Four accuracy metrics for the seven species from the Redwoods National and State Parks case study. Accuracy metrics are calculated using dataset observations from inside the 10th spatial cross-validation block (**Fig. S4B**). The number of observations is denoted alongside the species name. Models with the subscript 10 (i.e. $deepbiosphere_{10}$) were trained without observations from this region. Models with the subscript unif (i.e. $Inception_{unif}$) were trained with points uniformly sampled from across the state, including observations within the region (**Fig. S4A**). The "% obs. seen" column reports this value by indicating how many of the observations used to calculate the accuracy were seen by the model during training. For many species, including redwoods, $deepbiosphere_{10}$ has superior performance on unseen examples and only for Douglas Iris (*Iris douglasiana*) is it consistently outperformed by the MLP climatic model. *Deepbiosphere* = Joint TResNet; MLP = multilayer

perceptron; obs. = observation; cal. = calibrated; ROC = receiver operating characteristic curve; AUC = area under the curve; *spp* = per-species.

| Model | num. examples | RF | MLP | LR |
|---|---|---|---|---|
| *Deepbiosphere*$_{10}$ | 1,009 | **57.9365 [54.8942-59.127]** | **49.2063 [44.9735-50.9259]** | 45.5026 [44.1799-46.4286] |
| NPS vegetation map | 462 | 48.6772 [46.6931-49.2063] | 48.6772 [46.6931-49.2063] | **48.6772 [46.6931-49.2063]** |
| Random baseline | 1,009 | 11.6402 [11.2434-12.8307] | 7.672 [7.0106-8.3333] | 7.672 [7.0106-8.3333] |

**Table S11 | Accuracy of *deepbiosphere*-based vegetation type prediction compared to official map**
Single-label accuracy comparison of predicted alliance-level vegetation type using features extracted from *deepbiosphere*$_{10}$ compared to the official National Park Service alliance-level vegetation map across 10 bootstrapped cross-validation trials. To ensure at least a minimum of 1,000 examples are available to fit the extracted features of *deepbiosphere*$_{10}$, in each trial 300 field plots from the 489 plots reserved by the National Park Service for performing an accuracy assessment are used for fitting the feature extractor, leaving 189 for testing the feature extractor's performance. The National Park Service chose to subset down the total available field plots in the study to the highest quality field plots based on a spectral assessment, thus only 462 plots were used to generate the final National Park Service map. The random baseline consists of fitting each given feature extractor with random gaussian noise of the same dimensions as the feature vector of *deepbiosphere*$_{10}$. This baseline approximates the accuracy of simply predicting the most frequent alliance from the field plots. Reported statistics are median [1st quartile - 3rd quartile] across the ten cross-validation trials.

| | | *deepbiosphere* | *deepbiosphere* | NPS map | *deepbiosphere* | NPS map | *deepbiosphere* | NPS map |
|---|---|---|---|---|---|---|---|---|
| vegetation class | Num. examples | $ROC_{AUC}$ | f1-score | f1-score | precision | precision | recall | recall |
| Alnus rubra Forest | 8 | 0.813 | 0 | **0.588** | 0 | **0.556** | 0 | **0.625** |
| Ammophila arenaria Grassland | 8 | 1 | **0.941** | 0.308 | **0.889** | 0.4 | **1** | 0.25 |
| Bacharis pilularis Shrubland | 3 | 0.965 | **0.286** | 0 | **0.25** | 0 | **0.333** | 0 |
| Carex obnupta Herbaceous | 6 | 0.852 | **0.286** | 0 | **1** | 0 | **0.167** | 0 |
| Carex obnupta-Deschampsia cespitosa Grassland | 1 | 0.697 | 0 | **0.154** | 0 | **0.083** | 0 | **1** |
| Dune Herbaceous | 6 | 0.989 | **0.769** | 0.462 | **0.714** | 0.429 | **0.833** | 0.5 |
| Festuca idahoensis Grassland | 1 | 0.989 | 0.4 | **1** | 0.25 | **1** | **1** | **1** |
| Lithocarpus densiflorus-(Other) YG Mixed Forest | 3 | 0.86 | 0 | **0.143** | 0 | **0.091** | 0 | **0.333** |
| Montane Conifer-Hardwood Mixed Forest | 2 | 0.572 | 0 | **0.2** | 0 | **0.125** | 0 | **0.5** |
| Montane Hardwood Mixed Forest | 1 | 0.375 | 0 | 0 | 0 | 0 | 0 | 0 |
| Other Herbaceous | 1 | 0.989 | **0.5** | **0.5** | **0.333** | **0.333** | **1** | **1** |
| Perennial Grassland | 11 | 0.944 | **0.667** | 0.529 | **0.615** | 0.391 | 0.727 | **0.818** |
| Picea sitchensis-(Other) Forest | 23 | 0.933 | **0.727** | 0.588 | 0.762 | **0.909** | **0.696** | 0.435 |
| Pinus attenuata Forest | 9 | 0.993 | **0.889** | 0.556 | **0.889** | 0.556 | **0.889** | 0.556 |
| Pinus jeffreyii Forest | 9 | 0.982 | **0.667** | 0.615 | 0.833 | **1** | **0.556** | 0.444 |
| Pinus monticola Forest | 3 | 0.993 | 0.667 | **0.75** | **0.667** | 0.6 | 0.667 | **1** |
| Pinus radiata X attenuata YG Mixed Forest | 2 | 1 | **1** | **1** | **1** | **1** | **1** | **1** |
| Pseudotsuga menziesii-(Other) YG Mixed Forest | 6 | 0.81 | **0.533** | 0 | **0.444** | 0 | **0.667** | 0 |
| Pseudotsuga menziesii-Lithocarpus densiflorus YG Mixed Forest | 21 | 0.908 | **0.444** | 0.312 | **0.533** | 0.455 | **0.381** | 0.238 |
| Quercus garryana Forest | 12 | 0.997 | 0.815 | **0.96** | 0.733 | **0.923** | 0.917 | **1** |
| Riverine Herbaceous | 5 | 0.995 | **0.727** | 0.545 | **0.667** | 0.5 | **0.8** | 0.6 |
| Rubus Shrubland | 5 | 0.989 | **0.6** | 0.5 | 0.6 | **0.667** | **0.6** | 0.4 |
| Salix Shrubland | 1 | 0.995 | 0 | 0 | 0 | 0 | 0 | 0 |
| Sequoia sempervirens Mature Forest | 13 | 0.937 | 0.72 | **0.839** | **0.75** | 0.722 | 0.692 | **1** |
| Sequoia sempervirens-(Other) YG Mixed Forest | 27 | 0.91 | **0.676** | 0.389 | 0.561 | **0.778** | **0.852** | 0.259 |
| Typha latifolia Herbaceous | 2 | 0.997 | 0 | **0.667** | 0 | **1** | 0 | **0.5** |
| Num. higher accuracy classes | | | 15 | 10 | 13 | 13 | 15 | 12 |
| | overall accuracy | | 0.635 | 0.481 | | | | |

**Table S12 | Per-class accuracy of *deepbiosphere* vegetation type prediction compared to official map**
Breakdown of accuracy by vegetation class on unseen field plots using the best $deepbiosphere_{10}$ downstream classifier across all ten bootstrapped cross-validation trials compared to the RNSP map predictions. Vegetation classes with no observed plots have been excluded, leaving 27 classes for comparison. Three binary accuracy metrics are reported (precision, recall, F1-score) and one discrimination metric (AUC). Deepbiosphere = Joint TResNet; ROC = receiver operating characteristic curve; AUC = area under the curve; NPS = National Park Service.

| feature classifier | **RF** | **RF** | **MLP** | **MLP** | **LR** | **LR** |
|---|---|---|---|---|---|---|
| num. examples | 1,000 | 10,000 | 1,000 | 10,000 | 1,000 | 10,000 |
| *Deepbiosphere$_5$* | **62.1 [61.6-62.725]** | **66.2 [66.05-66.575]** | **63.6 [63.5-64.175]** | **62.75 [61.975-63.175]** | **64.2 [64.1-64.2]** | **64.1 [64.025-64.1]** |
| *TileNet* | 53.55 [53.3-54.2] | 57.9 [57.8-58.2] | 50.65 [50.15-51.2] | 53.35 [53.125-53.875] | 52.7 [52.7-52.7] | 56.1 [56.025-56.1] |
| Random | 42.0 [41.9-42.075] | 41.9 [41.825-41.9] | 24.75 [22.75-26.15] | 25.35 [25.15-26.025] | 25.5 [25.4-25.575] | 19.8 [19.8-19.9] |

**Table S13 | Comparison of *deepbiosphere* crop type accuracy to state-of-the-art unsupervised learning method**

Single-label accuracy of predicted crop type using the indicated feature classifier fitted with features extracted from *deepbiosphere$_5$* compared to *TileNet* from ref. [67] across ten instantiations of said classifier. Extracted features are identical across the ten trials; only the feature classifiers themselves are re-fitted. The random baseline consists of fitting each given feature classifier using random gaussian noise with the same dimensions as the features used to fit the *deepbiosphere$_5$* feature classifiers. This baseline approximates the accuracy of simply predicting the most frequent crop type from the training locations. Reported statistics are median [1st quartile - 3rd quartile] across ten separate instantiations of said downstream classifier.

| | *deepbiosphere* | *TileNet* | *deepbiosphere* | *TileNet* | *deepbiosphere* | *TileNet* | *deepbiosphere* | *TileNet* | *deepbiosphere* | *TileNet* |
|---|---|---|---|---|---|---|---|---|---|---|
| CDL class | frequency | frequency | $AUC_{ROC}$ | $AUC_{ROC}$ | f1-score | f1-score | precision | precision | recall | recall |
| 1 | 11 | 11 | **0.956** | 0.848 | 0 | 0 | 0 | 0 | 0 | 0 |
| 111 | 12 | 12 | **0.849** | 0.726 | **0.154** | 0 | **1** | 0 | **0.083** | 0 |
| 121 | 21 | 43 | **0.937** | 0.623 | 0 | **0.07** | 0 | **0.143** | 0 | **0.047** |
| 122 | 16 | 37 | **0.937** | 0.771 | **0.19** | 0.139 | **0.4** | 0.143 | 0.125 | **0.135** |
| 123 | 36 | 33 | **0.983** | 0.969 | **0.614** | 0.484 | **0.519** | 0.379 | **0.75** | 0.667 |
| 124 | 16 | 15 | **0.997** | 0.989 | **0.581** | 0.333 | 0.6 | **1** | **0.562** | 0.2 |
| 131 | 1 | 3 | **0.985** | 0.484 | 0 | 0 | 0 | 0 | 0 | 0 |
| 176 | 65 | 55 | 0.947 | **0.948** | **0.63** | 0.548 | **0.526** | 0.407 | 0.785 | **0.836** |
| 190 | 1 | 2 | **1** | 0.708 | 0 | 0 | 0 | 0 | 0 | 0 |
| 2 | 3 | 5 | **0.988** | 0.85 | 0 | **0.4** | 0 | **0.4** | 0 | **0.4** |
| 204 | 8 | 13 | **0.711** | 0.653 | 0 | 0 | 0 | 0 | 0 | 0 |
| 206 | 3 | 2 | **0.988** | 0 | 0 | 0 | 0 | 0 | 0 | 0 |
| 21 | 2 | 2 | **0.487** | 0.466 | 0 | 0 | 0 | 0 | 0 | 0 |
| 212 | 31 | 27 | **0.973** | 0.909 | 0 | 0 | 0 | 0 | 0 | 0 |
| 225 | 5 | 5 | **0.918** | 0.729 | 0 | 0 | 0 | 0 | 0 | 0 |
| 236 | 3 | 5 | **0.651** | 0.564 | 0 | 0 | 0 | 0 | 0 | 0 |
| 24 | 8 | 12 | **0.897** | 0.707 | 0 | 0 | 0 | 0 | 0 | 0 |
| 36 | 66 | 61 | **0.895** | 0.848 | 0.27 | **0.322** | 0.522 | **0.538** | 0.182 | **0.23** |
| 4 | 3 | 1 | **0.821** | 0 | 0 | 0 | 0 | 0 | 0 | 0 |
| 49 | 5 | 5 | **0.851** | 0.488 | 0 | 0 | 0 | 0 | 0 | 0 |
| 54 | 7 | 7 | **0.968** | 0.624 | **0.333** | 0 | **0.4** | 0 | **0.286** | 0 |
| 61 | 40 | 37 | **0.904** | 0.773 | **0.133** | 0.035 | **0.2** | 0.05 | **0.1** | 0.027 |
| 69 | 420 | 384 | **0.909** | 0.844 | **0.748** | 0.711 | **0.643** | 0.642 | **0.895** | 0.797 |
| 75 | 206 | 205 | **0.92** | 0.855 | **0.69** | 0.625 | **0.711** | 0.598 | **0.67** | 0.654 |
| 76 | 6 | 10 | **0.925** | 0.587 | 0 | 0 | 0 | 0 | 0 | 0 |
| | Num. higher accuracy classes | | 24 | 1 | 9 | 3 | 8 | 3 | 7 | 5 |
| | | overall accuracy | 0.622 | 0.535 | | | | | | |

**Table S14 | Per-crop accuracy of *deepbiosphere* compared to state-of-the-art unsupervised learning method**
Breakdown of accuracy by cropland data layer class on unseen locations using the best *deepbiosphere* downstream classifier fitted with 1,000 examples. Cropland classes with fewer than two examples in the test set have been excluded. Frequencies are reported as calculated for the largest area class for the respective receptive field size of the deep learning model. Three binary accuracy metrics are reported (precision, recall, F1-score) and one discrimination metric (AUC). *Deepbiosphere* = Joint TResNet; ROC = receiver operating characteristic curve; AUC = area under the curve; CDL = cropland data layer.
